# Daily rewiring of a neural circuit generates a predictive model of environmental light

**DOI:** 10.1101/2020.08.31.275313

**Authors:** Bryan J. Song, Slater J. Sharp, Dragana Rogulja

## Abstract

Ongoing sensations are compared to internal, experience-based, reference models; mismatch between reality and expectation can signal opportunity or danger, and can shape behavior. The nature of internal reference models is largely unknown. We describe a model that enables moment-to-moment luminance evaluation in flies. Abrupt shifts to lighting conditions inconsistent with the subjective time-of-day trigger locomotion, whereas shifts to appropriate conditions induce quiescence. The time-of-day prediction is generated by a slowly shifting activity balance between opposing neuronal populations, LNvs and DN1as. The two populations undergo structural changes in axon length that accord with, and are required for, conveying time-of-day information. Each day, in each population, the circadian clock directs cellular remodeling such that the maximum axonal length in one population coincides with the minimum in the other; preventing remodeling prevents transitioning between opposing internal states. We propose that a dynamic predictive model resides in the shifting connectivities of the LNv- DN1a circuit.

---

The brain assigns valence to incoming sensory stimuli, allowing responsiveness to be context-dependent. To do this, it must continually check ongoing sensations against expectations, an idea known as predictive coding^1^. Discrepancy between expected and actual outcomes is famously reflected in the activity of dopaminergic neurons^2^, which have been proposed to encode reward prediction error^3^. The mechanisms that give rise to this integrated signal are not yet understood. It has been theorized that dopaminergic neurons are a point of convergence between ongoing sensory signals and expectations generated from prior outcomes^1^. In order to understand the computations underlying predictive coding, it is necessary to identify the sensory and predictive information streams that are integrated by downstream comparators. While sensory circuits have been successfully mapped in many systems, the representation of expectations in the brain remains largely unknown.

Predictable schedules enable planning for future events. Biological mechanisms have evolved to take cues from regular environmental fluctuations and organizing preparatory physiological and behavioral processes. For example, anticipating nightfall means seeking food and shelter before sundown. This is made possible by circadian clocks, conserved molecular oscillators that operate on a ∼24 hour schedule^4^ and are entrained by rhythmic cues such as daily cycles of light^5^ or temperature^6^. Under normal light-cycling conditions, the progression of daytime and nighttime is tracked by the rise and fall of key circadian clock proteins^7^. When external cues are removed (animals are put into constant darkness), molecular clocks continue to cycle^7^. These clocks give proper timing to rhythmic processes such as sleeping and feeding^8^, but there is no evidence that they are used moment-to-moment to assess conditions in the environment. We show that the circadian system assists in prediction evaluation, and describe a mechanism by which it does this: a microcircuit within the network of circadian neurons uses cellular remodeling as a strategy to organize slowly shifting internal predictions. The experimental paradigm we established provides a new way to study how expectations are encoded and evaluated.

## Results

### Locomotor reactivity to light depends on time-of-day

We used a protocol similar to Lu et al^9^, where flies experienced light and darkness alternating every 12 hours for several days (mimicking daytime and nighttime) before spending at least 24 hours in darkness. They were then exposed to light for an hour at different times of day (**Fig. 1a**). When light was presented during the nighttime, wild-type^10^ flies immediately increased locomotion (startle, **Fig. 1a,b** and **Extended Data Video 1**). During subjective daytime (daytime, but in darkness) their reaction was the opposite – they immediately slowed down or stopped moving (**Fig. 1a,b** and **Extended Data Video 1**). The difference in baseline locomotion (**Extended Data Fig. 1a**) does not explain differential responsiveness between day and night, based on the following: locomotion was similar at 8pm and 8am, but diverged when lights turned on (**Fig. 1b**); responsiveness in individuals showed no correlation with baseline locomotion, and weak correlation with sleep status (**Extended Data Fig. 1a,b**); normalizing light-evoked locomotion to baseline did not change the results (**Extended Data Fig. 1c**). Though males generally sleep during the day while females do not^11^, both sexes responded to daytime light by decreasing activity (**Fig. 1a,b**; **Extended Data Fig. 2a,b**).

**Figure 1.**
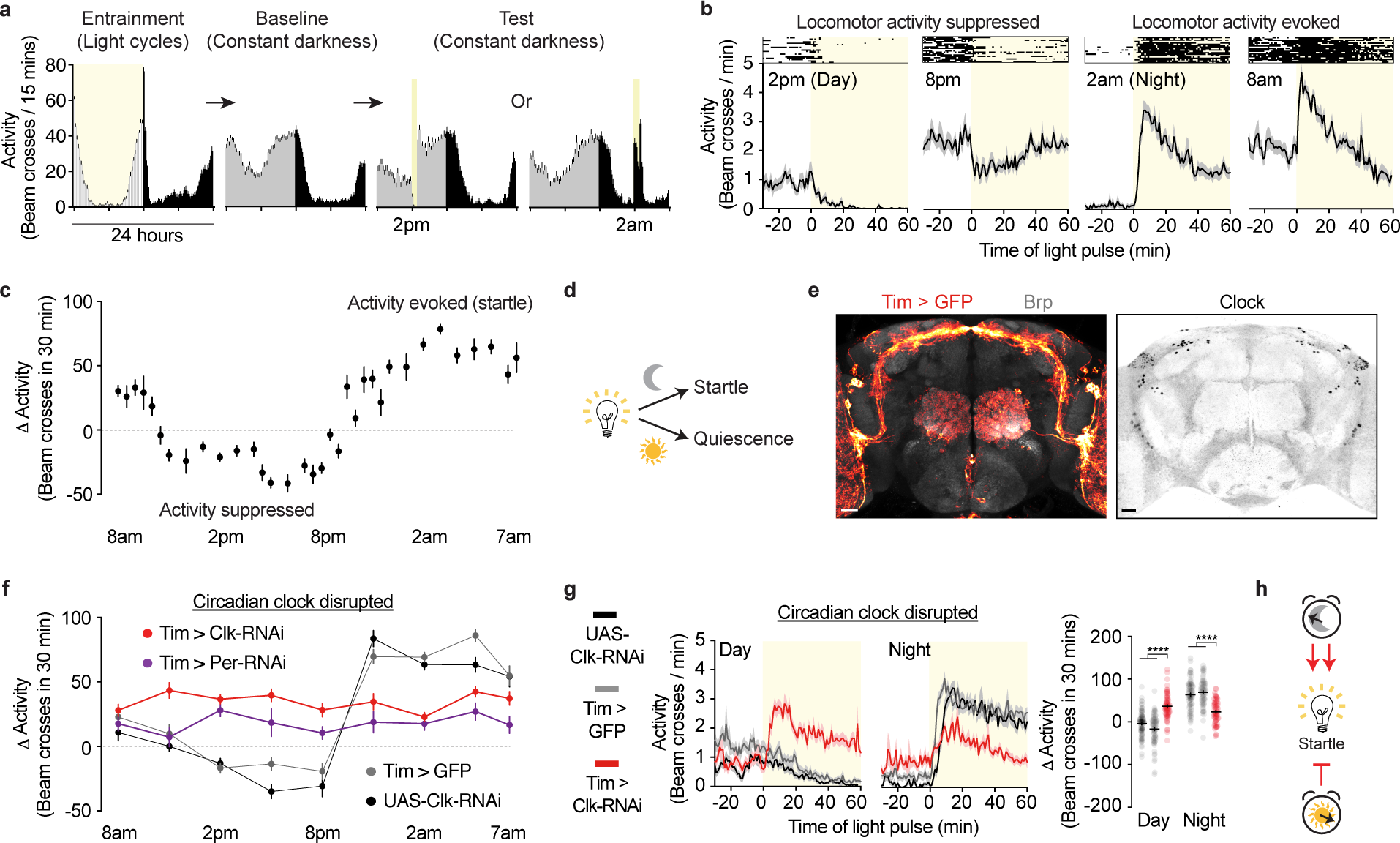
Circadian clocks bidirectionally modulate light responsiveness. **(a)** Experimental protocol and daily locomotor activity of an isogenic wild-type strain, *w^+^iso31*, during entrainment, baseline, and test periods. Vertical bars: 15-minute bins of locomotor activity, where height indicates mean. Error bars, S.E.M. Yellow indicates light; reactivity to light depends on time-of day, even though animals are in complete darkness. **(b)** A closer view of acute locomotor responses to light at different times. Boxed insets show representative activities of individual flies; below, averaged activity of all tested wild-type flies. These data are collected at 1-minute intervals. Shading: error bars (S.E.M in all figures). **(c)** Change in activity evoked by light, in independent wild-type cohorts, across a 24 hour period. **(d)** Schematic of daily light responses in wild-type flies. **(e)** Left, Tim-Gal4 is expressed throughout the circadian network, labeled with GFP. The whole brain is visualized with an antibody against Bruchpilot (Brp), a presynaptic protein. Tim-Gal4 is also expressed in noncircadian neurons in the antennal lobe and glia^60^, which was partially blocked by inclusion of repo-Gal80 (data not shown) in all experiments using this driver. Right, clock neurons labeled with an antibody against Clock. **(f)** Light-evoked changes in activity across 24 hours in controls (black and gray), and in flies in which circadian proteins Clock (Clk, red) or Period (Per, purple) were depleted with RNAi. Independent cohorts were tested at different time points. **(g)** Clock disruption (red) perturbs acute light responsiveness during daytime and nighttime. **(h)** Proposed model for daily switches in light contextualization, regulated by the clock. For all figures, behaviors were analyzed with Two-way ANOVA, Tukey’s post-test, unless otherwise indicated. *p<0.05, **p<0.01, ***p<0.001, ****p<0.0001 for all figures. Extended Data Table 1 shows sample sizes for all figures. For all figures, scale bars: 20 µm.

**Figure 2.**
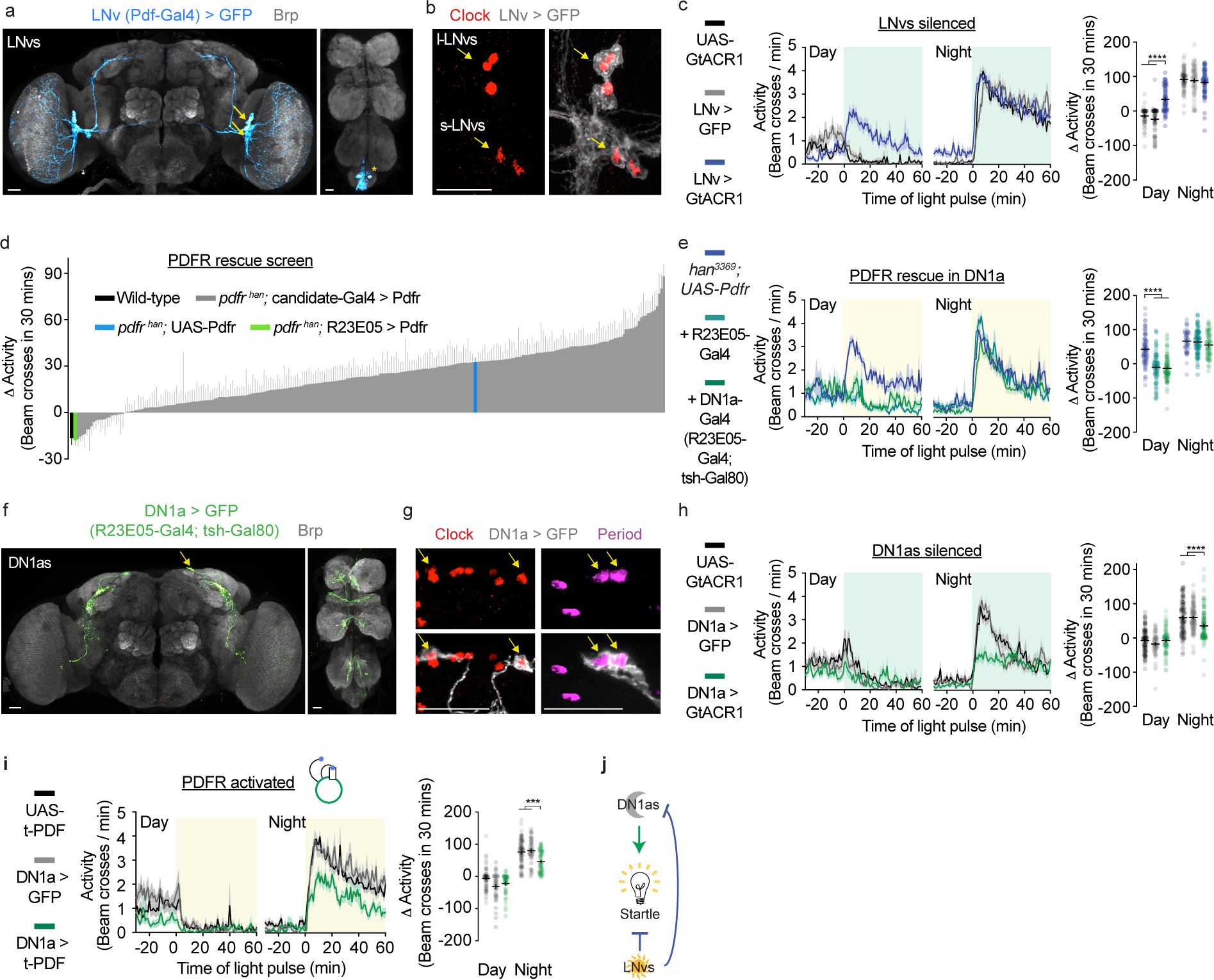
Different clock neurons are required for normal daytime, vs nighttime, reactivity to light. **(a)** Left, LNv neurons in the brain labeled with GFP. Arrows point to LNv cell bodies. Right, non-LNv expression in the ventral nerve cord (VNC) indicated by asterisk. LNv-Gal4 is driven by *pdf* regulatory elements^18, 61^. **(b)** Clock staining in the LNvs. **(c)** Silencing LNvs using the light-gated chloride channel GtACR1 perturbs mid-day but not mid-night light responsiveness. For all optogenetic experiments, the light used for neuronal silencing simultaneously served to probe behavior. **(d)** A *pdfr* genetic rescue screen reveals a role for LNv-to-DN1a transmission in contextualizing daytime light. PDFR was expressed in candidate neuronal populations in *han*^3369^ or *han^5304^ pdfr* mutant backgrounds. **(e)** Activity traces for flies in which PDFR was expressed using R23E05-Gal4 (dark green) or using R23E05-Gal4 with teashirt-Gal80 (DN1a-Gal4, light green) in the *han^3369^ pdfr* mutant background. This experiment was also done in the *han^5304^ pdfr* mutant background (Figure S4A). **(f)** DN1as visualized by GFP (DN1a > GFP). **(g)** Clock and Period staining in DN1as. **(h)** DN1a silencing perturbs mid-night but not mid-day light responsiveness. **(i)** Constitutive activation of PDFR with t-PDF in DN1as attenuates nighttime response to light. **(j)** Model of LNv and DN1a roles in contextualizing light.

Two opposing states of responsiveness lasted ∼12 hours each (**Fig. 1c**), correlating with the schedule of light and darkness previously experienced during entrainment. To test whether time-of-day light responses are indeed instructed by prior experience, we entrained flies to shortened or lengthened light schedules (**Extended Data Fig. 3a**). At the same hour (7pm), a light pulse either suppressed or evoked locomotor activity, depending on whether light had been on at 7pm during entrainment (**Extended Data Fig. 3b-f**). Locomotion always seemed triggered by subjective mismatch (i.e. experiencing different conditions than expected at that time of day). In support of this idea, startle was evoked not only by nighttime light (**Fig. 1a-c**), but also by daytime darkness (**Extended Data Fig. 3g,h**). Locomotor reactivity to light pulses therefore reports internal estimates of daytime vs nighttime with moment-to-moment resolution (**Fig. 1d**).

**Figure 3.**
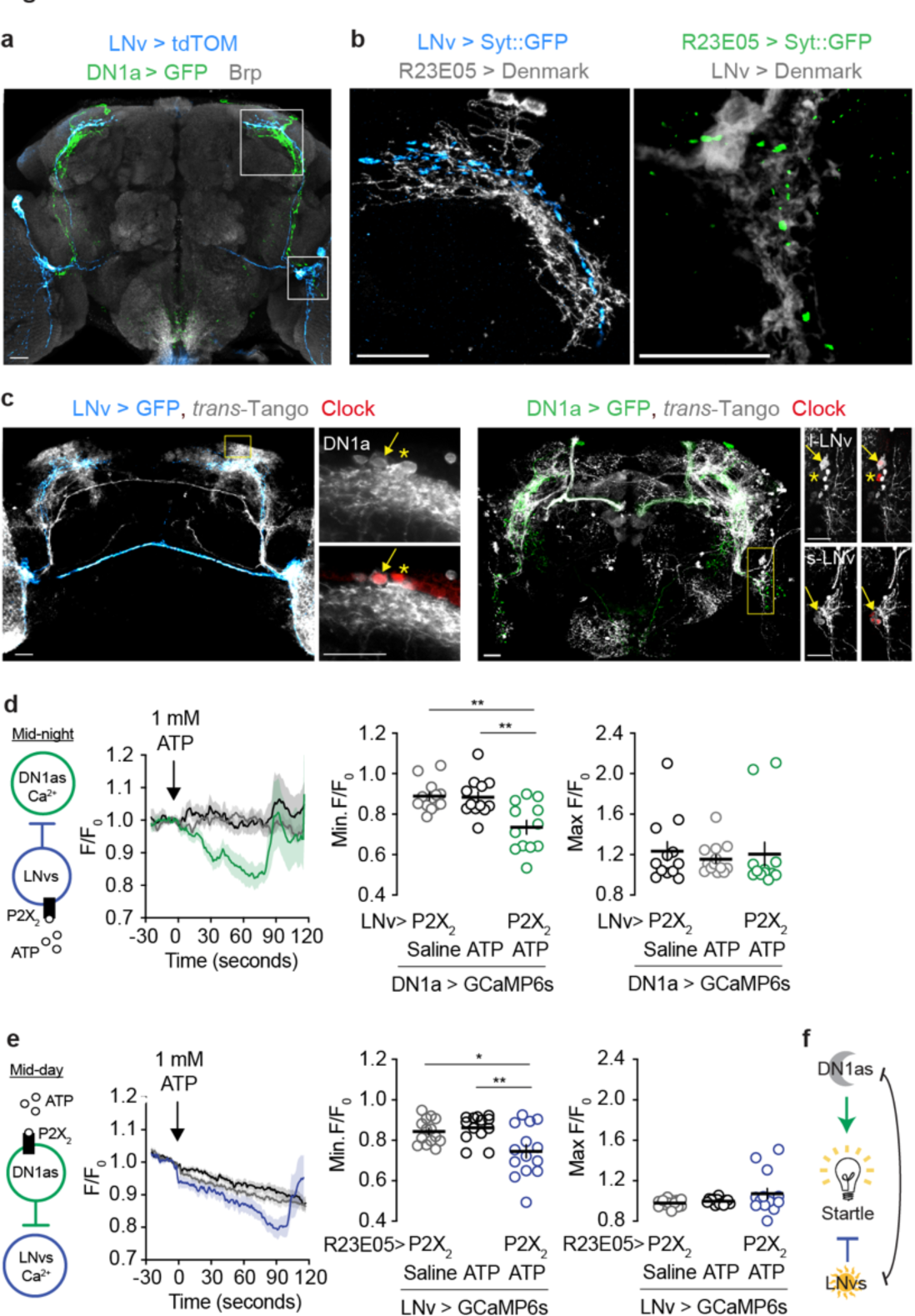
Reciprocal targeting between LNvs and DN1as. **(a)** Expression of GFP in DN1as, and tdTomato in LNvs, reveals overlapping projections. **(b)** LNv axons overlap with DN1a dendrites (left), and *vice versa* (right). Synaptotagmin (Syt): presynaptic marker; Denmark: postsynaptic marker. Left, area outlined on the top in (a). Right, area outlined on the bottom in (A). **(c)** Postsynaptic targets labeled by *trans*-Tango indicate that LNvs and DN1as target each other. Arrows point to target cells expressing Clock. Asterisks: DN1as and LNvs not labeled by *trans*-Tango. **(d)** Chemogenetic activation of LNvs during the nighttime silences DN1as *ex vivo*. **(e)** Chemogenetic activation of DN1as during the daytime silences LNvs *ex vivo*. In all conditions, a steady decline in GCaMP6s fluorescence was seen in LNvs, likely due to photobleaching. For and (e), measurements were taken from flies that were in light-dark cycles (at ∼2pm (ZT6) for daytime, at ∼2am (ZT18) for nighttime). Minimal and maximal changes in calcium are reported for each trial. One-way ANOVA with Tukey’s post-hoc test. (**f**) Model of LNv and DN1a inhibitory circuit connectivities.

### Circadian clocks contextualize environmental light

Based on the timescales involved, we suspected the involvement of circadian clocks, molecular programs that are entrained by environmental cues and that organize daily rhythms in physiology and behavior^12^. When core clock proteins Clock^12^ or Period^13–15^ (Per) were depleted from the network of ∼150 circadian neurons^16, 17^ (**Fig. 1e**, **Extended Data Fig. 4a-d**), flies lost the ability to contextualize light relative to time of day. Instead, they always responded with a startle – but this startle was weak relative to the nighttime startle in controls (**Fig. 1f,g**). The implication is that the circadian system bidirectionally modifies a stereotyped behavioral response to an abrupt change in luminance - during daytime (when light is appropriate) clocks suppress the startle, but they enhance it during nighttime (when light is inappropriate (**Fig. 1h**). Predictions originate from molecular clock oscillations, as mutants with faster clocks^13^ cycled through light-responsive states faster (**Extended Data Fig. 5a,b** and **Extended Data Table 2**).

**Figure 4.**
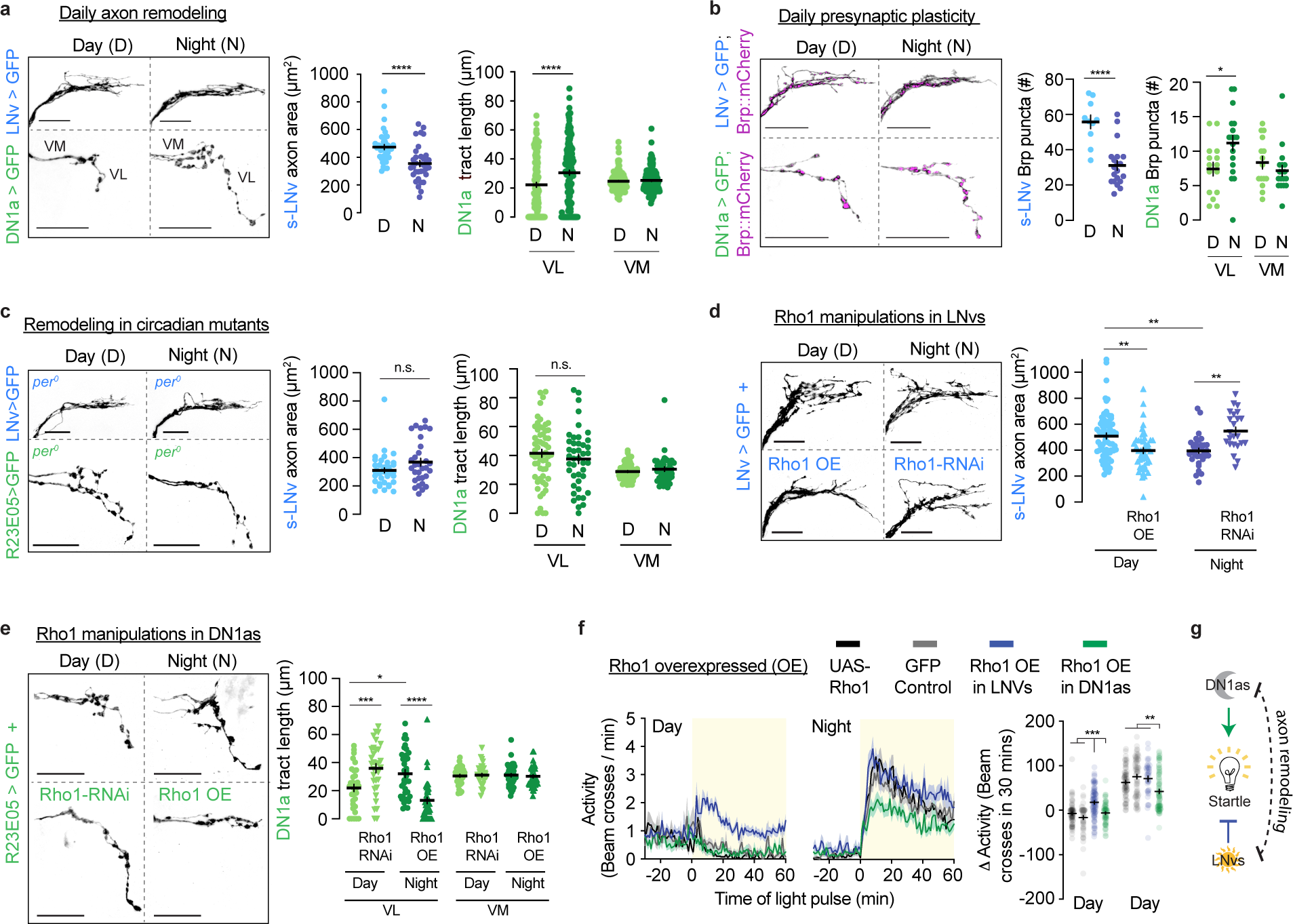
Axon remodeling in LNvs and DN1as is required for normal light reactivity. **(a)** LNvs and DN1as show antiphase oscillations in neurite morphology. Measurements were taken from flies that were in light-dark cycles (at 2pm (ZT6) for daytime, at 2am (ZT18) for nighttime). Quantifications show data for individual brain hemispheres. For this figure, t-Test in LNvs, Two- way ANOVA with Tukey’s post-hoc test in DN1as. Ventromedial (VM) tracts are internal controls for ventrolateral (VL) tracts in DN1as. **(b)** Daily changes in presynaptic site number in LNvs and DN1as, as reported by a synaptic marker Brp:mCherry (magenta) within GFP-labeled neurites (black). **(c)** *Period* mutants lack LNv and DN1a plasticity rhythms. **(d)** Manipulating Rho1 in LNvs (Rho1-RNAi vs Rho1 overexpression (OE)) bi-directionally changes axonal fasciculation. One- way ANOVA with Tukey’s post-hoc test. **(e)** Manipulating Rho1 in DN1as bi-directionally changes axonal fasciculation. One-way ANOVA with Tukey’s post-hoc test. **(f)** Flies with fasciculated (closed) dorsal LNv axons (Rho1 OE in LNvs, blue) have perturbed daytime, but not nighttime, light responsiveness. The opposite is true in flies in which DN1a axons are kept short (Rho1 OE in DN1as, green). ‘GFP control’ is LNv > GFP during the daytime, and DN1a > GFP during the nighttime. **(g)** Schematic summary of LNv-DN1a circuit state transitions that regulate opponent predictions about light.

### Separate clock neuron subpopulations contextualize daytime and nighttime light

To find the neuronal mechanism that organizes predictions about environmental light, we first examined LNvs^12^ (**Fig. 2a,b**). This small group of neurons regulates normal locomotor activity rhythms (i.e. the pattern of activity seen under basal conditions, where periods of high and low activity occur at predictable times of day)^18^. Using the green-light-gated chloride channel GtACR1^19^, we silenced LNvs conditionally, avoiding potential developmental artifacts (Methods). Here light served as both a visual stimulus and effector for GtACR1. LNv silencing caused flies to startle in response to light during subjective daytime (**Fig. 2c**), as if they no longer held the expectation that light during the daytime is appropriate – mimicking the daytime phenotype of flies lacking clocks entirely (**Fig. 1g**). The near-instantaneous nature of optogenetics allows us to conclude that LNvs contextualize daytime light on a moment-by-moment basis. Surprisingly, unlike general clock disruption (**Fig. 1g**), LNv silencing produced no nighttime phenotype (**Fig. 2c**). Daytime-specific phenotypes were also seen with RNAi-mediated depletion of the LNv- specific neuropeptide Pigment Dispersing Factor^20^ (PDF, **Extended Data Fig. 6a**), and with hypomorphic mutations in the PDF receptor (PDFR)^21^ (**Extended Data Fig. 6b,c**). Though knocking down PDF in the small LNv subpopulation^20^ (**Extended Data Fig. 7a,b,d**) was sufficient to disrupt daytime light responsiveness (**Extended Data Fig. 7c,e**), the phenotype was stronger when the small and large LNvs were manipulated simultaneously (**Extended Data Fig. 6a**), so we treated them as a unit.

Because LNvs are considered to be a central pacemaker^18^, a reasonable concern is that LNv- disrupted flies lack circadian rhythms entirely. Several lines of evidence argue for specific, rather than general, loss of clock function. First, LNvs silencing did not affect nighttime responsiveness to light (**Fig. 2c**, **Extended Data Fig. 6a**), unlike when the entire clock network was disabled (**Fig. 1g**). Second, the arrhythmic phenotypes of LNv disruption have been argued to stem from developmental problems^22^, which are avoided with conditional optogenetics. Third, it takes many days for LNv silencing to cause arrhythmicity - most animals are rhythmic during the first two days of constant darkness^18^ (**Extended Data Table 2**), which is when our testing is done. Finally, other subpopulations can support timekeeping in the absence of LNv function^22^.

We conclude that LNvs signal that light is appropriate during daytime but are dispensable for contextualizing light during nighttime.

LNv silencing recapitulates only the daytime phenotype of clock disruption, suggesting that other populations might have analogous function during the nighttime. Since LNvs signal through the peptide PDF, we looked for their targets by restoring expression of the PDF receptor to various neuronal populations in the receptor mutants (Fig. 2d, Extended Data Fig. 8a). For this pdfr rescue screen, we predominantly tested Gal4 lines expressed sparsely in the nervous system, and also looked at lines expressed in sleep- and locomotion-regulating centers, circadian subpopulations, and neurons expressing specific neurotransmitters or peptides (Extended Data Table 3). Restoring PDFR to most neuronal populations, including known LNv targets^23–25^, could not fully suppress the pdfr mutant phenotype (Extended Data Table 3). Only a few of the 274 tested Gal4 lines allowed complete rescue; of these, we focused on the lines with the most restricted expression. When PDF transmission was enabled onto neurons labeled by R23E05-Gal4^26^, normal responsiveness to daytime light (decrease in locomotor activity) was restored (Fig. 2d,e; Extended Data Fig. 8b). R23E05 labels ∼20 neurons in the ventral nerve cord, and ∼10 neurons in the brain (Extended Data Fig. 9a,b). The brain neurons include the four dorsal-anterior clock neurons (DN1as; Fig. 2f,g; Extended Data Fig. 9c). DN1a dendrites and cell bodies are in the dorsomedial protocerebrum, and their axons descend towards the accessory medulla^27, 28^ (Extended Data Fig. 9d). We restricted expression of R23E05-Gal4 to mostly DN1as with addition of teashirt-Gal80 (tsh-Gal80), a Gal4 inhibitor expressed in the ventral nerve cord (Extended Data Fig. 9e). We refer to this intersectionally- derived driver as DN1a-Gal4.

DN1a silencing had the opposite effect from LNv silencing, perturbing only the *nighttime* response to light (**Fig. 2h**). The phenotype – a less robust startle - was similar to the nighttime phenotype of clock-disrupted flies (**Fig. 1g**) suggesting that without DN1as flies can no longer evaluate light as inappropriate during the night. The only other studies of DN1a function in adults found that these neurons produce the neuropeptide CCHamide1 (CCHa1)^29^ (**Extended Data Fig. 10a**), and promote wakefulness in the morning^29, 30^. In our assay, there was no effect of depleting CCHa1 or its receptor (CCHa1R, **Extended Data Fig. 10b**), showing that the function we found for DN1as is distinct from what was previously observed^29^ – these neurons signal that light is inappropriate during nighttime, which is complimentary to the daytime role of LNvs.

Attenuated responsiveness to nighttime light could reflect general locomotor and visual deficits, so we tested whether all genotypes were capable of robust visual and motor function. We reanalyzed data from key experiments (**Fig. 1g**; **Fig. 2c,h**) to see if the animals whose responsiveness to light was attenuated ever reached high levels of locomotor activity. There was no significant difference between experimental animals and controls (**Extended Data Fig. 11a**). The lower population-averaged responsiveness in DN1a-silenced, or clock-disrupted flies, is instead accounted for by infrequency of high activity bouts (**Extended Data Fig. 11b**) which suggests that these animals were less likely to be in a startled state. To further assess locomotor vigor, we tested animals with mechanical stimulation and found that all genotypes reacted with high levels of locomotor activity, easily exceeding levels elicited by light (**Extended Data Fig. 11c**). A visually guided behavior, courtship^31, 32^, was also normal (**Extended Data Fig. 11d**, Methods). Taken together, these data argue that clock neuron manipulations do not simply impair sensory input or motor output, but instead disrupt the ability to contextualize light.

### LNv and DN1a clock neurons are mutually interconnected

LNvs and DN1as appear to have opposite and complimentary functions which suggests that these subpopulations might be somehow coordinated. This theory is supported by the fact that LNvs use PDF to signal onto DN1as during the daytime (**Fig. 2d**). Based on their opponency, we tested whether LNv-to-DN1a signaling is inhibitory, by expressing membrane-tethered PDF^33^ in DN1as. Since tethered PDF is anchored to the membrane, it has short-range, cell-autonomous effects on cells that natively express PDF receptor^34^. Ectopic nighttime PDFR activation attenuated the nighttime startle to light (**Fig. 2i**), similar to the DN1a silencing phenotype (**Fig. 2h**). This result confirms that DN1as express PDFR^35, 36^ and suggests that PDF normally inhibit DN1as during the daytime. Taken together, results presented thus far suggest that LNvs and DN1as have opposite roles in contextualizing light during daytime vs nighttime, and that the peptide PDF is a crucial organizational signal between these subpopulations (**Fig. 2j).**

The interdigitated arrangement of LNv and DN1a projections **(Fig. 3a**, **Extended Data Fig. 12a** and **Extended Data Video 2**) is suggestive of reciprocal communication. Genetically encoded markers of pre-and post-synaptic sites ^37, 38^ showed that LNv axons terminate onto DN1a dendrites within the superior lateral protocerebrum, while DN1a axons terminate onto LNv dendrites within the accessory medulla (**Fig. 3b**, **Extended Data Videos 3,4**). The trans-synaptic tracing tool *trans*-Tango^39^ indeed reported LNvs and DN1as as mutual synaptic targets (**Fig. 3b** and **Extended Data Fig. 12b,c**). To test whether these putative connections are functional, we activated each population at times when their activity is predicted to be low, and looked at the response of the other population *ex vivo.* When LNvs were stimulated via the ATP-gated cation channel P2X2^40, 41^, the calcium sensor GCaMP6s^42^ reported transient inhibition in DN1as (**Fig. 3d**; **Extended Data Fig. 13a,b**). Conversely, stimulating DN1as led to transient LNv inhibition (**Fig. 3e**; **Extended Data Fig. 13c,d**). The weaker effects of DN1a stimulation could be due to biological reasons, or because R23E05-LexA is a weaker driver than PDF-LexA (**Extended Data Fig. 14a**). These results suggest that a reciprocal inhibition motif within the *Drosophila* circadian circuit contributes to opponent predictions about light (**Fig. 3f**).

### Presynaptic structural plasticity regulates behavioral state transitions

LNv and DN1a neurons are required to contextualize light at different times of day and are mutually connected. How does the LNv-DN1a circuit alternate between activity states? It was known that s-LNv axons undergo daily structural remodeling, spreading out in the morning and bundling up at night^43–46^, but the function of this change has remained mysterious^47–51^. We discovered that DN1a axons are also remodeled daily, on a schedule that is anti-phase to LNvs (**Fig. 4a**) - their axons are extended at night and retracted during the day. The fluorescently tagged presynaptic protein Bruchpilot (Brp) showed that changes in presynaptic area correspond with changes in synapse number (**Fig. 4b**, **Extended Data Fig. 15a,b)**. The rhythmicity of axonal remodeling is set by the circadian clock, as it was absent in *period* mutants (**Fig. 4c)**. These results raise the possibility that daily changes in connectivity within a mutually inhibitory LNv- DN1a microcircuit underlie transitions between light-predictive states.

The LNv-DN1a circuit appears in distinct physical configurations during the day (more LNv output sites) vs night (more DN1a output sites). For each population, the time of day when their axons occupy the most space correlates with the time when that population is necessary. To test the idea that axonal structural remodeling contributes to the light-predictive internal model, we looked for manipulations that can affect remodeling in the two populations. We found that DN1as might utilize similar cellular programs as LNVs^46^ - manipulating the GTPase Rho1 levels bidirectionally affected remodeling in both populations. For both LNvs and DN1as, Rho1 overexpression (OE) decreased axonal area, while RNAi-mediated Rho1 depletion increased axonal area (**Fig. 4d,e**).

In agreement with the idea that remodeling supports transitions between opponent predictive states, Rho1 overexpression in LNvs caused increased locomotion in response to daytime light, while overexpression in DN1as attenuated the startling effect of nighttime light (**Fig. 4f**). That is, preventing presynaptic area from increasing phenotypically resembles silencing (**Fig. 2c,h**).

Rho1 overexpression did not appear to overtly damage the LNv neurons, as animals had relatively intact locomotor activity rhythms (**Extended Data Fig. 16a** and **Extended Data Table 2**) and did not have accelerated evening locomotor activity onset, which occurs when LNvs are ablated or constitutively silenced^18, 52^ (**Extended Data Fig. 16a**). The daytime phenotype of Rho1 overexpression in LNvs, and nighttime phenotype of Rho1 overexpression in DN1as, together match the light response phenotypes seen when circadian clocks are disabled (**Fig. 1g**). These data fit a model in which structural plasticity biases the outcome of LNv-DN1a reciprocal inhibition, leading to a flexible internal model of what the light conditions should be at any moment (**Fig. 4g**).

## Discussion

Alterations of neuronal activity, rather than morphology, are usually considered the cause of cognitive flexibility^53^. The mechanism that we describe relies on physical cellular restructuring. What are the advantages of a system like this? While near-instantaneous electrical activity is the basic language of neurons, many behaviors and internal states occur on much longer timescales. Morphological remodeling is a slower process, aligning with functions that change over the course of several hours. In support of this view, changes in neuronal morphology have been found to underlie appetite^54^, sexual experience^55^, and foraging history^56^. Though seemingly wasteful, physical remodeling may be particularly useful for encoding relatively stable states, due to presumably high energetic barrier.

Understanding the mechanisms of circuit state transitions may help clarify the etiology of mood disorders like depression of bipolar disorder, which are characterized by the lack, or excess, of transitions between extreme states. Disorders like these may reflect a collapse of organizational principles that normally permit flexible circuit function. An unsolved question is how behavioral states can be stable across long timescales, but also undergo flexible transitions. Motifs from the LNv-DN1a circuit illustrate one solution to this apparent contradiction. LNvs and DN1as are arranged in a mutually inhibitory system, which may help ensure consistency and accuracy over long timescales. In the absence of external influence, reciprocal inhibition can stabilize a winner-take-all steady state^57^. Structural plasticity is a potential way to overcome this inflexibility, by providing a molecular mechanism to overcome electrical inhibition. We show that *Drosophila* make remarkably accurate estimates of daytime and nighttime, which may be enabled by flexible transitions between stable circuit configurations.

A predictive nervous system enables continual evaluation of reality relative to context. One result of this is that a fixed stimulus can evoke a multitude of behaviors depending on an animal’s history, needs, and external context. We show how the circadian system creates a dynamic internal reference of what environmental conditions *should* be. The paradigm that we developed offers opportunities to understand the interface between internal models and sensory evidence. Circadian neurons are sensitive to environmental inputs – can they autonomously compute prediction error? Clock neurons communicate with downstream dopaminergic populations^58, 59^: are those analogous to mammalian midbrain dopaminergic neurons whose activities reflect prediction error? We propose that flies assign valence to experienced environmental conditions, a computation that utilizes an internal model generated through circuit remodeling.

## Acknowledgments

We thank our lab, the Crickmore lab, Charles Weitz, Michael Do, Matt Pecot, Gordon Fishell, and Michael Rosbash for advice and comments on the manuscript. Tara Kane assisted with experiments. Stephen Zhang helped with coding and curated a collection of sparse driver lines from FlyLight. Stephen Thornquist helped with optogenetics. We thank Rachel Wilson and her lab for saline and for calcium imaging advice; and Corey Harwell for microscope access. For fly stocks, we thank David Anderson, Andreas Bergmann, Justin Blau, Adam Claridge-Chang, Michael Crickmore, Barry Dickson, Paul Hardin, Robert Kittel, Michael Nitabach, Matt Pecot, Jeff Price, Orie Shafer, Amita Sehgal, Paul Taghert and Rachel Wilson. We are especially grateful to Dr. Gerry Rubin and the Howard Hughes Medical Institute’s Janelia Research Campus for access to JRC_SS00681-Gal4 and JRC_SS00645-Gal4 lines prior to publication. These strains will be described further in Dionne, Rubin, and Nern (manuscript in preparation). For antibodies, we thank Paul Hardin and Amita Sehgal.

## Author contributions

B.S. and D.R. designed the study. B.S., S.S. and D.R. performed experiments. B.S. and D.R. analyzed the data and wrote the paper.

## EXPERIMENTAL PROCEDURES

### Drosophila melanogaster stocks

All *Drosophila* stocks used in this study are listed in the key resource table. Flies were grown on cornmeal-agar medium at 25°C under 12 hour light : 12 hour dark conditions in a room with ∼70 lux white light. UAS-myr::GFP, UAS-mCD8::GFP, UAS-tethered PDF, and UAS-Dicer2 lines were outcrossed six or seven times into the control *iso31* background. No differences in light responses were observed between outcrossed and non-outcrossed lines. One experiment (**Fig. 1f,g (**2pm and 2am only) had UAS-mCD8::GFP controls in the GFP condition, whereas the genotype is UAS-myr::GFP elsewhere in the paper. No behavioral differences were seen between UAS-mCD8::GFP and UAS-myr::GFP. RNAis were co-expressed with Dicer2 (Dcr) to increase efficiency ^62^. “LNv-Gal4” used in this study has two copies of Pdf-Gal4, on the second and third chromosomes. Similar results for silencing and neuropeptide knockdown were found using a single copy of Pdf-Gal4 on the second chromosome, although effect sizes were smaller (data not shown). Wild-type *w^+^iso31* wild-type strains were created by backcrossing *Canton S* six times into the *iso31* background. Detailed genotypes and samples sizes for each experiment are provided in **Extended Data Table 1**. Origin of each fly stock is shown in **Extended Data Table 4**. Stocks are available upon request.

### Generation of R23E05-LexA and R23E05-Gal80

Standard Gateway cloning protocols (Thermo Fisher Scientific, 11791020 and 11789020) were followed to derive constructs in which either LexA or Gal80 are driven by the R23E05 enhancer. gtcccgatttcgtcgaaggattcaa forward and gctaaccggatgacggtaccaggag reverse primers were used to PCR-amplify a 644kb enhancer fragment from R23E05-Gal4 flies. This product was subcloned into pBPLexa:P65UW (Addgene plasmid # 26231, ^63^ or pBPGal80uw-6 (Addgene plasmid # 26236, ^63^), which were gifts from Gerry Rubin. Resulting constructs were inserted into the attP2 landing site by embryo injection (Rainbow Transgenics).

### Locomotor activity measurements

Male flies, 1-9 days old, were collected and individually housed in 65 mm glass tubes with approximately 20 mm of cornmeal-agar media. To avoid cumulative effects of repeated exposure to light, separate cohorts were tested for each trial. They were given at least 48 hours to acclimate before experiments began. Female flies were collected as virgins (<1 day old) upon eclosion, and group housed for at least 3 days before testing. Activity and sleep were measured using the Trikinetics *Drosophila* activity Monitor system, which counts infrared beam crosses through the midline of the glass tube. Experiments were conducted in DigiTherm CircKinetics incubators (Tritech Research, DT2-CIRC-TK) at 25°C. Aside from optogenetic experiments, light pulses were delivered using white incubator lights (∼260 lux white fluorescent bulbs, ∼102 μW/mm^2^ in the 470 nm range). Most experiments in **Fig. 2d** and all experiments in **Extended Data Fig. 11c** and **Extended Data Video 1** were conducted in larger incubators (Percival, DR- 41VL) to accommodate video cameras, the mechanical stimulation apparatus, or the large number of locomotor activity monitors required to screen for LNv-Downstream neurons (Fig 2d).

### Optogenetics

For 48-96 hours before the experiment, control and experimental flies were fed 50 mM all-trans-retinal (Sigma Aldrich R2500) that was diluted in ethanol (Koptec, V1001) and coated onto rehydrated potato food (Carolina Bio Supply Formula 4-24 Instant *Drosophila* Medium, Blue). For GtACR1 experiments, six 530 nm green LEDs (Luxeon Rebel, LXML-PM01-0100) were driven by a 700 mA constant current driver (LuxDrive BuckPuck, 03021-D-E-700), and pulse-width modulated signal to an averaged intensity of ∼202 μW/mm^2^. LEDs were placed along the wall of the incubator and controlled with an Arduino Uno Rev3 (Arduino, A000066) microcontroller using a custom script. Between replicates, genotypes were positionally counterbalanced within the incubator to control for nonuniform illumination from the light source (LEDs or white fluorescent bulb). Silencing motor neurons with VGlut-Gal4 allowed us to confirm that all flies received enough illumination to access neurons expressing GtACR1^19, 64^, regardless of position within the incubator. Light intensity measurements were recorded using a power meter (Thorlabs, PM100D). Lux measurements were recorded using a light meter (Extech, LT300). The spectra of ambient white light in the laboratory and in experimental incubators were measured with a spectrometer (Thorlabs, CCS200). All reported measurements were taken with devices facing the light source. Power measurements for white light were taken at the 470 nm range.

### Mechanical stimulation

Flies were shaken using a multi-tube vortexer (Trikinetics TVOR-120) modified to house *Drosophila* Activity Monitors (Trikinetics). The vortexer was programmed to deliver medium intensity vibrations continuously for an hour.

### Courtship assay

Courtship assays were conducted as previously described^65^. Briefly, male flies were isolated at least five days before the assay, to allow recovery of mating drive^65, 66^. On the day of the assay, one male was aspirated into a cylindrical chamber (10 mm diameter and 3 mm height) with one virgin *w^+^iso31* female. Flies were videotaped from above using a handheld camera (Canon, Vixia HRF800) and videos were manually scored for courtship behaviors. Courtship indices were calculated from the percentage of time spent in mating behaviors during a five minute window following courtship initiation (as indicated by unilateral wing extension). If flies did not engage in courtship throughout the entire 15 minute assay, they were given a courtship index value of 0. Flies were illuminated from below (∼3.7 μW/mm^2^ in the 475 nm range) using a light pad (Artograph, LightPad 930), in addition to aforementioned overhead white room lighting. Because optogenetic LEDs interfered with the ability to visualize and record flies, we relied on the white light from the light pad as an optogenetic effector. While the lightpad was substantially dimmer than the LEDs used in our optogenetic silencing experiments, our control experiments with other drivers showed that this light could in principle penetrate the cuticle to effect GtACR1 in the nervous system. For positive controls, we verified the lightpad’s efficacy in inducing paralysis or extending mating duration in flies where GtACR1 was expressed by VGLUT-Gal4 or Corazonin-Gal4 respectively (data not shown, ^19, 67^).

### Immunohistochemistry

Flies were anesthetized under CO2. Brains were then dissected in cold Schneider’s medium (Gibco, 21720-001) and immediately fixed in 4% paraformaldehyde (PFA, Electron Microscopy Sciences, 15710). After a 20 minute fixation at room temperature, brains were washed three times with PBS containing 0.3% Triton X-100 (Amresco, M143-1L), 20 minutes per wash, and blocked overnight with 10% donkey serum (Jackson ImmunoResearch, 017-000-121) at 4°C. Primary and secondary antibodies were diluted in donkey serum and incubated with brains for 48 hours each. For Brp (nc82) stainings, the primary antibody incubation was conducted for 72 hours due to the large number of Brp epitopes in the brain. Three 20 minute washes were done after primary and secondary antibody incubations.

Primary antibodies used: Guinea pig anti-Clock antibody (Gift from Paul Hardin, 1:2000 dilution), Chicken anti-GFP antibody (Aves, GFP-1020, 1:1000 dilution), Mouse anti-Brp antibody (Developmental Studies Hybridoma Bank (DSHB), NC82, 1:7 dilution), Mouse anti-PDF antibody (DSHB, PDF C7, 1:100 dilution), Rabbit anti-DsRed antibody (Clontech, 632496, 1:100 dilution), Guinea pig anti-Period antibody (Gift from Amita Sehgal, 1:50 dilution), Rabbit anti-CCHa1 (Our lab raised antibodies against the peptide QIDADNENYSGYELT ^68^, Genscript, 1:50 dilution). Secondary antibodies used: Donkey anti-Mouse 488 (Thermo Fisher Scientific, A-21202, 1:1000 dilution), Donkey anti-Rabbit 568 (Thermo Fisher Scientific, A-10042, 1:1000 dilution), Donkey anti-Mouse 647 (Thermo Fisher Scientific, A-31571, 1:1000 dilution), Donkey anti-Guinea Pig 488 (Jackson ImmunoResearch, 703-545-148, 1:100 dilution), Donkey anti-Chicken 488 (Jackson ImmunoResearch, 703-545-155, 1:100 dilution), Donkey anti-Guinea pig Cy3 (Jackson ImmunoResearch, 706-165-148, 1:100 dilution), Donkey anti-Guinea pig 647 (Jackson ImmunoResearch, 706-605-148, 1:100 dilution).

Tissues were whole-mounted in Prolong Gold Antifade reagent (Invitrogen, 1942345) on glass slides with coverslips (Electron Microscopy Sciences, 64321-10, 72230-01). Confocal images were obtained using a Leica SP8 confocal microscope at 10x, 2.4 µm intervals, for morphology quantifications; 20x, 1 µm intervals for expression patterns, and 63x, 0.3 µm intervals for imaging of pre- and postsynaptic sites. Maximum projection images and quantifications were obtained using FIJI. Levels of brightness and contrast were adjusted across the whole image using FIJI or Adobe Photoshop.

For quantifications comparing neurite morphologies between mid-day and mid-night, whole heads were fixed because prolonged exposure to light (required for dissections) can modify the operation of the clock. Heads were fixed for 50 minutes at room temperature with fixative containing 4% PFA and 0.3% Triton X-100. For mid-night samples, heads were fixed with minimal light exposure, using red light that is less disruptive to the light-sensitive clock protein Cryptochrome^69–72^. Acquisitions were conducted at 10x due to the large number of samples; as the drivers we used are expressed sparsely, this resolution was sufficient. Because the z-axis of the slide/coverslip chamber is slightly shorter than the height of the brain, all of the brains were pressed slightly, and in similar orientation.

For *trans*-Tango experiments, flies were raised for 5 weeks at 18°C, which permits stronger expression than 25°C^39^. We often noticed aberrant morphology in cells expressing the *trans*-Tango construct, likely due to overexpression of neurexin and the cell adhesion molecule ICAM1 at presynaptic sites^39^.

### Calcium imaging

Experiments were conducted in a 6-hour window centered around periods of putative peak activity (mid-day for DN1a->LNv and mid-night for LNv->DN1a), alternating between control and experimental samples. *Ex vivo* whole mount brains were explanted in Nunclon cell culture dishes (Thermo Scientific, 150318) which contained 3 mL of chilled *Drosophila* saline (Gift from Rachel Wilson, 103 mM NaCl, 3 mM KCl, 5 mM N-tris (hydroxymethyl) methyl-2-aminoethane-sulfonic acid, 8 mM trehalose, 10 mM glucose, 26 mM NaHCO3, 1 mM NaH2PO4, 1.5 mM CaCl2, and 4 mM MgCl2 (osmolarity adjusted to 270–275 mOsm). Saline was bubbled with 95/5% carboxygen prior to the experiment. Brains were dissected in the same media used to conduct the experiment. Brains in which GCaMP was expressed in LNvs were allowed to rest for 2.5 minutes prior to the experiments under the blue imaging light.

Brains in which GCaMP was expressed in DN1as were allowed to rest for 5 minutes prior to the experiments under the blue imaging light. During this baseline period, we noticed increased large calcium transients (**Extended Data Fig. 13a,c**), likely due to control of clock neuron activity by the light-sensitive protein Cryptochrome^73, 74^. During pilot experiments we chose baseline intervals that were usually sufficient to allow activity to stabilize. Two trials were excluded (one experimental and one control) because baseline activities were not stable. These trials are shown as red traces in **Extended Data Fig. 13a,b**. For P2X2 experiments, 20 μL of 150 mM ATP (Sigma, A2383), diluted in *Drosophila* saline, was pipetted gently down the side of the dish, to a final concentration of 1 mM. Positive control experiments in which both P2X2 and GCaMP were expressed in LNvs showed that ATP delivered this way could start inducing small changes within a few frames of delivery. Acquisition occurred at 1 frame/second.

### Quantifying locomotor activity

Sleep and activity data were analyzed using custom Matlab software (available on github at https://github.com/CrickmoreRoguljaLabs) and plotted in Graphpad Prism 8 for Macintosh.

Activity counts were collected at 1 minute intervals. A sleep episode was defined as inactivity lasting at least five minutes^75, 76^.

Circadian analysis was conducted using the Cycle-P function in FaasX, using 30-minute bins^77^. Our experiments occurred during the second day of darkness, but rhythmicity, tau and power of locomotor rhythms (**Extended Data Table 2**) were calculated during 4 days in darkness.

Additional days of analysis allowed us to acquire more accurate measurements^77, 78^. This analysis is conservative, because circadian deficits grow stronger with more time spent in darkness^18^.

### Quantifying morphological imaging data

Measurements were conducted blind using the segmented line tool in FIJI on the maximum intensity projection of whole brain z-stacks. No obvious daytime-nighttime differences were observed in the z-axis for the DN1a ventrolateral tract. For Brp quantifications, acquisitions were done at 63x. The sparsity of synaptic sites along the ventrolateral DN1a tract allows for visualization and counting of individual punctum. Brp counts were conducted blind. Each hemisphere was computed as an independent sample because of variability between hemispheres.

### Quantifying immunostaining intensity

For quantifications of fluorescence intensities used to validate the efficiency of RNAi, the experimenter was not blinded during quantification, which we deemed acceptable due to the large and consistent effect sizes. Regions of interest were selected using the freehand selection tool in FIJI, on summed intensity projections of whole brain z-stacks. For measurements of Clock- and Period- RNAi efficacy, intensity measurements were taken within the most visible s-LNv cell body per brain because all 5 s-LNvs could not all be easily identified in the knockdown conditions. For measurements of GFP intensity, when comparing the strength of LexA drivers, there was substantial variability between hemispheres.

Thus measurements were taken for both hemispheres and subsequently averaged.

### Quantifying *ex vivo* calcium imaging

Data were analyzed with ImageJ, using the freehand selection tool to choose a ∼100μm ROIs from s-LNv dorsal terminals. Only the brighter hemisphere was used for analysis. LNv axons and DN1a dendrites were chosen for quantification because these regions were consistently identifiable, whereas DN1a axons and LNv dendrites were not usually visible with GCaMP6s. For figures, 2-3 frames (two seconds) of data was removed from each sample because of motion artifacts from pipetting. Unaltered trials are in reported in **Extended Data Fig. 13a,c**. Baseline fluorescence was calculated from the average of ten frames prior to ATP delivery, excluding the first frame before ATP. Minimum and maximum fluorescence was calculated using standard Excel functions from all frames after ATP delivery except for the first five frames, to exclude potential residual motion artifacts. 1-3 samples per condition showed drift after pipetting, thus we used an ImageJ registration plugin (TurboReg) to create a new series corrected against a time-series averaged reference. Two nonrepresentative trials (one experimental trial and one control) were excluded from averaged results shown in **Fig. 3e**. These excluded trials are shown in **Extended Data Fig. 13b** and were excluded due to unusually large and early depolarizations that were putatively due to the effects of blue light stimulation.

### Statistical analysis

All statistical tests were conducted with Prism 8 for Macintosh (GraphPad). All data are presented as mean ± S.E.M. For significance indicators (asterisks) referring to multiple post hoc tests, we conservatively report the largest (least significant) p-value from each of the tests.

Exact p values are in Extended Data Table 1. Only significant comparisons are indicated. For light probe experiments, group means were compared using a Two-way ANOVA with Tukey’s post-hoc comparisons against all possible conditions. For all behavioral panels using two-way ANOVA, multiple comparisons between time points are not reported, except for **Extended Data Fig. 1a**. Significant differences between control genotypes are not indicated in figures. We report them here: in **Fig 2i** the two parental controls significantly differed during the day (p = 0.0111), the two parental controls in **Extended Data Fig. 6b**. significantly differed from each other at night (p < 0.0001), and in **>Extended Data Fig. 10c**, DN1a>GFP was significantly different from both other conditions during the day, p = 0.0384 vs. DN1a > CCHa1 RNAi and p = 0.0005 vs UAS parental control). Behavioral experiments in main figures each had at least three replicates of approximately sixteen flies each. In cases where genotypes were only tested at one time point, we used a One-way ANOVA followed by Tukey’s post-hoc test. For imaging experiments where we quantified LNv or DN1a morphology, we treated each hemisphere as a single sample because we noticed substantial variability between hemispheres. In **Figure 4**, comparisons between VL and VM DN1a tracts are not reported. Comparisons between Rho1 overexpression and Rho1-RNAi are also not reported. Power analyses to predetermine sample size were not conducted. Experimenters were not blind to conditions except during quantifications of morphology. Sample sizes are shown in **Extended Data Table 1**.

### DATA AND SOFTWARE AVAILABILITY

All data and materials are available upon request.

## EXTENDED DATA

**Extended Data Figure 1.**
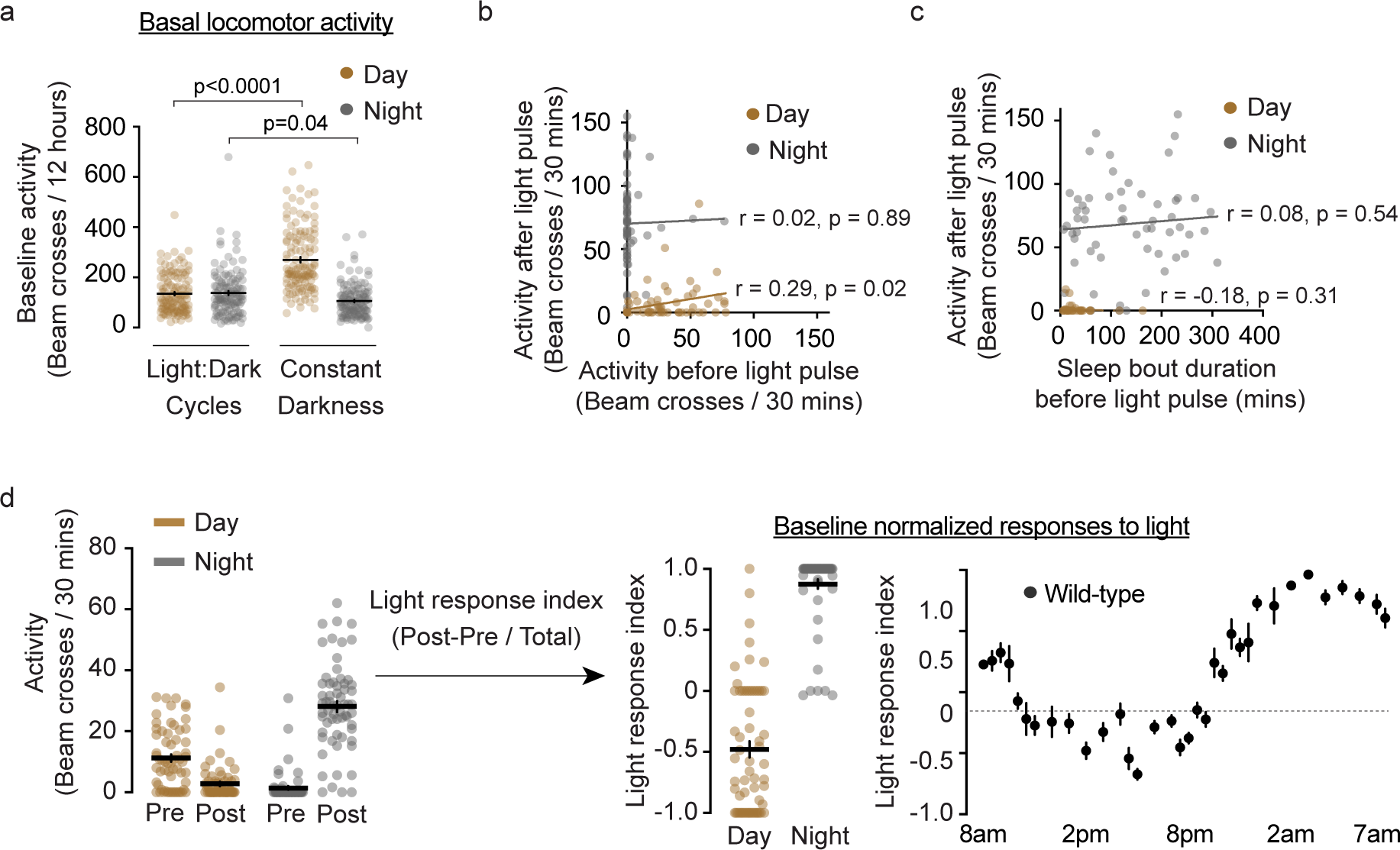
Light responsiveness is independent of circadian fluctuations in baseline locomotion. **(a)** Basal locomotor activity is higher in constant darkness than in light- dark cycles, but only during daytime. **(b)** Pre-pulse locomotor activity weakly correlates with activity during the pulse given mid-day (p=0.02). Pre-pulse locomotor activity does not correlate with activity during the pulse given mid-night (p=0.89). **(c)** Sleep bout duration prior to the light pulse does not correlate with behavioral response during the mid-day (p=0.31) or mid-night (p=0.54). (a-c) show the same flies as Fig.1, a-c. **(d)** Activity normalized to baseline still shows two distinct states. Same flies as Fig. 1c.

**Extended Data Figure 2.**
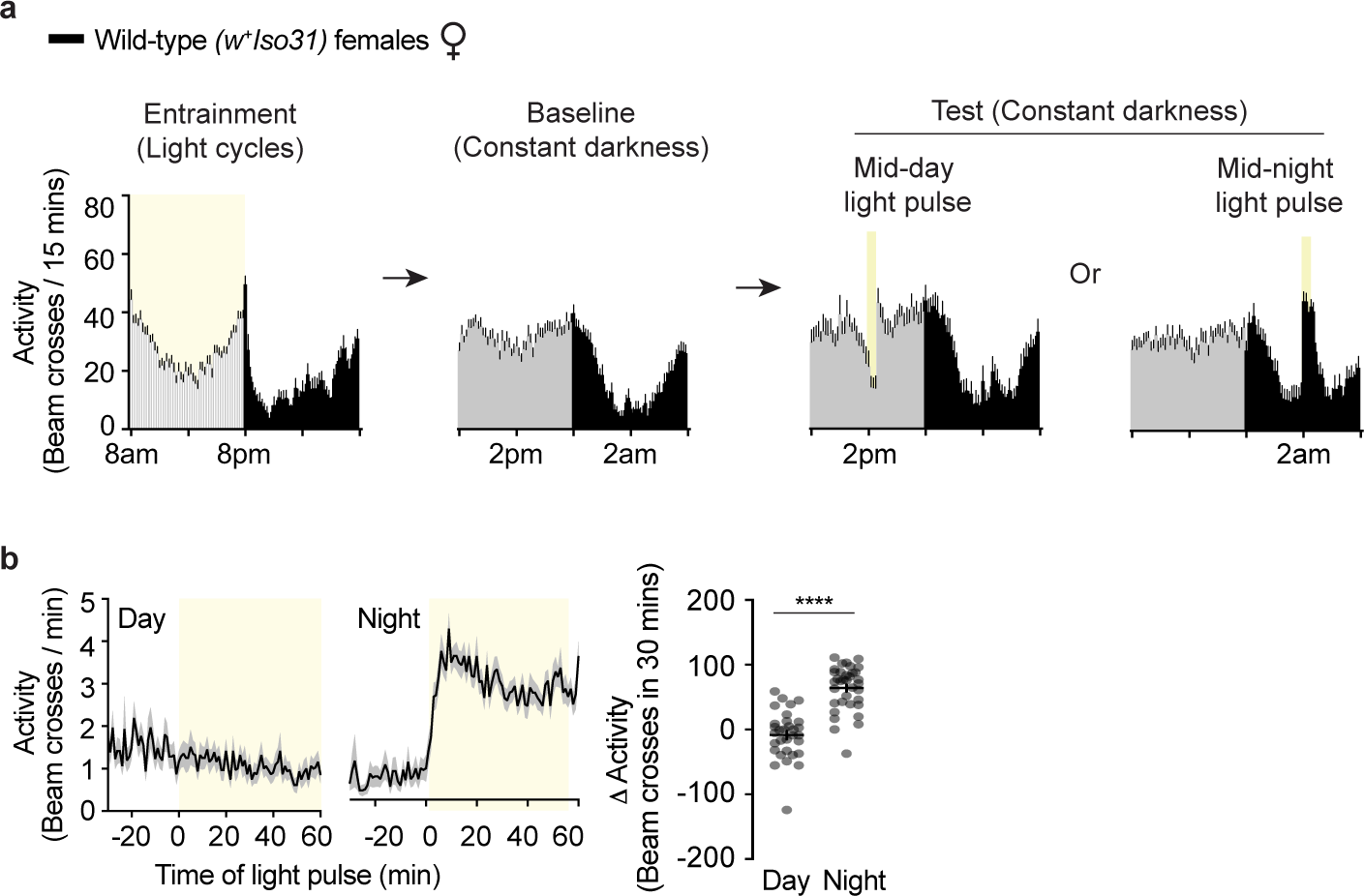
Female flies also respond to light differently during the daytime vs nighttime. **(a)** Experimental setup. Virgin female *w^+^Iso31* flies underwent the same treatment shown in Fig. 1a. For entrainment and baseline, 24 hours of activity are shown. **(b)** Left, Averaged activity 30 minutes prior to, and during, the 60 minute light probe. Right, quantification of light-evoked change in activity.

**Extended Data Figure 3.**
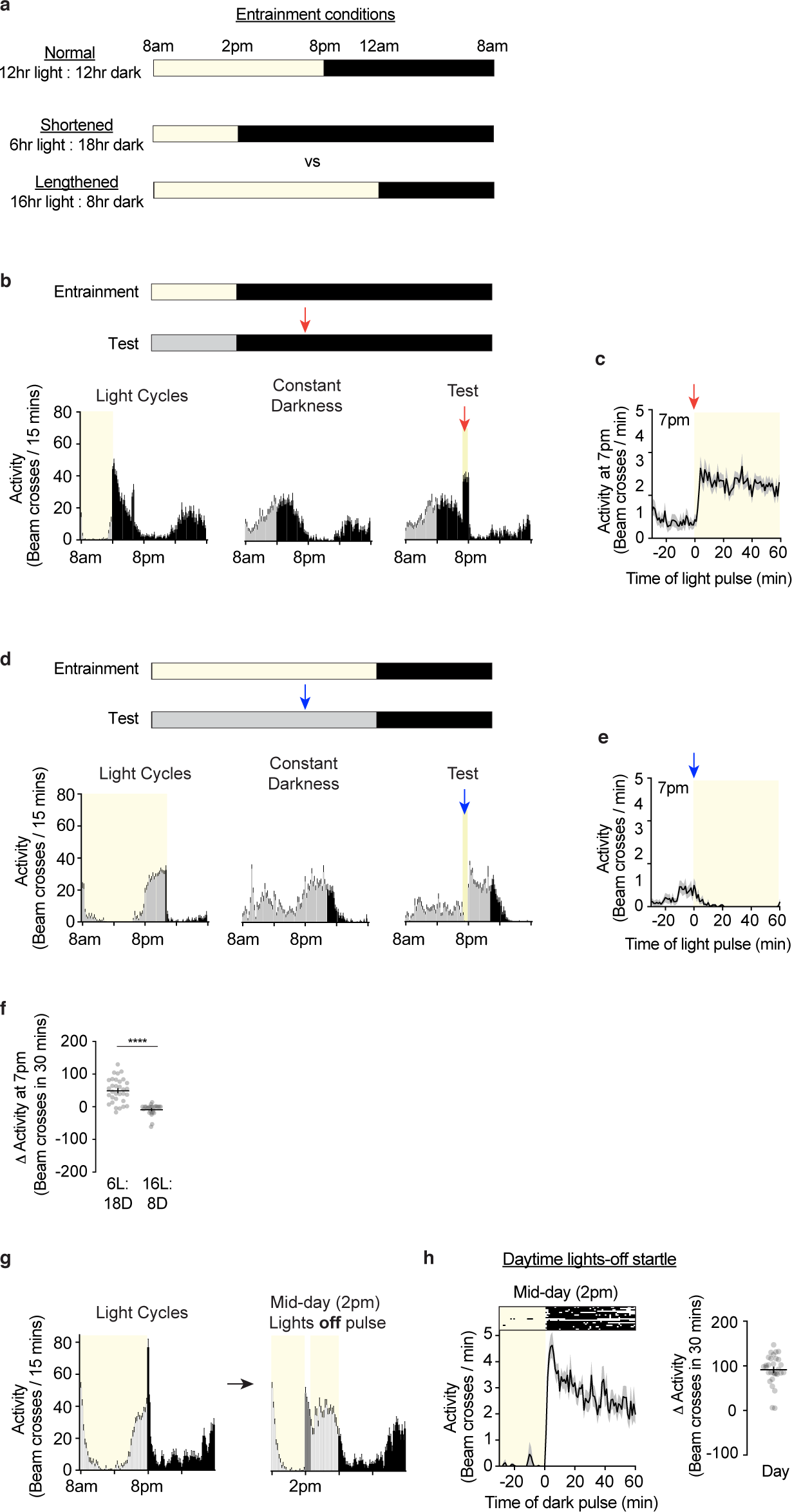
Locomotor responses to light are instructed by prior entrainment experience. Darkness during daytime acutely evokes locomotor activity. **(a)** The top rows show the standard entrainment protocol used in all other experiments. The bottom two rows show alternative light-dark entrainment cycles. In (a-e), yellow bars indicate when lights are on, black bars indicate when lights are off, and grey bars indicate subjective daytime in constant darkness. **(b)** 6 hour light : 18 hour dark conditions were used for entrainment and flies were subsequently tested in constant darkness with a one-hour light pulse. Arrows indicate time of light onset. **(c)** A light pulse at 7pm evokes locomotor activity for flies entrained in 6:18 light:dark cycles. **(d)** Flies were entrained to 16 hour light : 8 hour dark conditions before testing. **(e)** A light pulse at 7pm suppresses locomotor activity for flies entrained to 16:8 light:dark cycles. **(f)** Quantification of differences between light-evoked responses at 7pm for flies entrained to 6:18 or 16:8 light:dark cycles. **(g)** Experimental protocol for testing the effect of acute darkness during daytime. Vertical bars: 15 minute locomotor activity bins. Yellow indicates light. Unlike for experiments shown in other figures, flies were not in constant darkness but only in light-dark cycles. On the experimental day, lights were turned off for 1 hour between 2pm and 3pm. **(h)** Lights-off during the daytime elicits startle. Top, representative individual raster plots. Bottom, averaged activity prior to and during the dark probe.

**Extended Data Figure 4.**
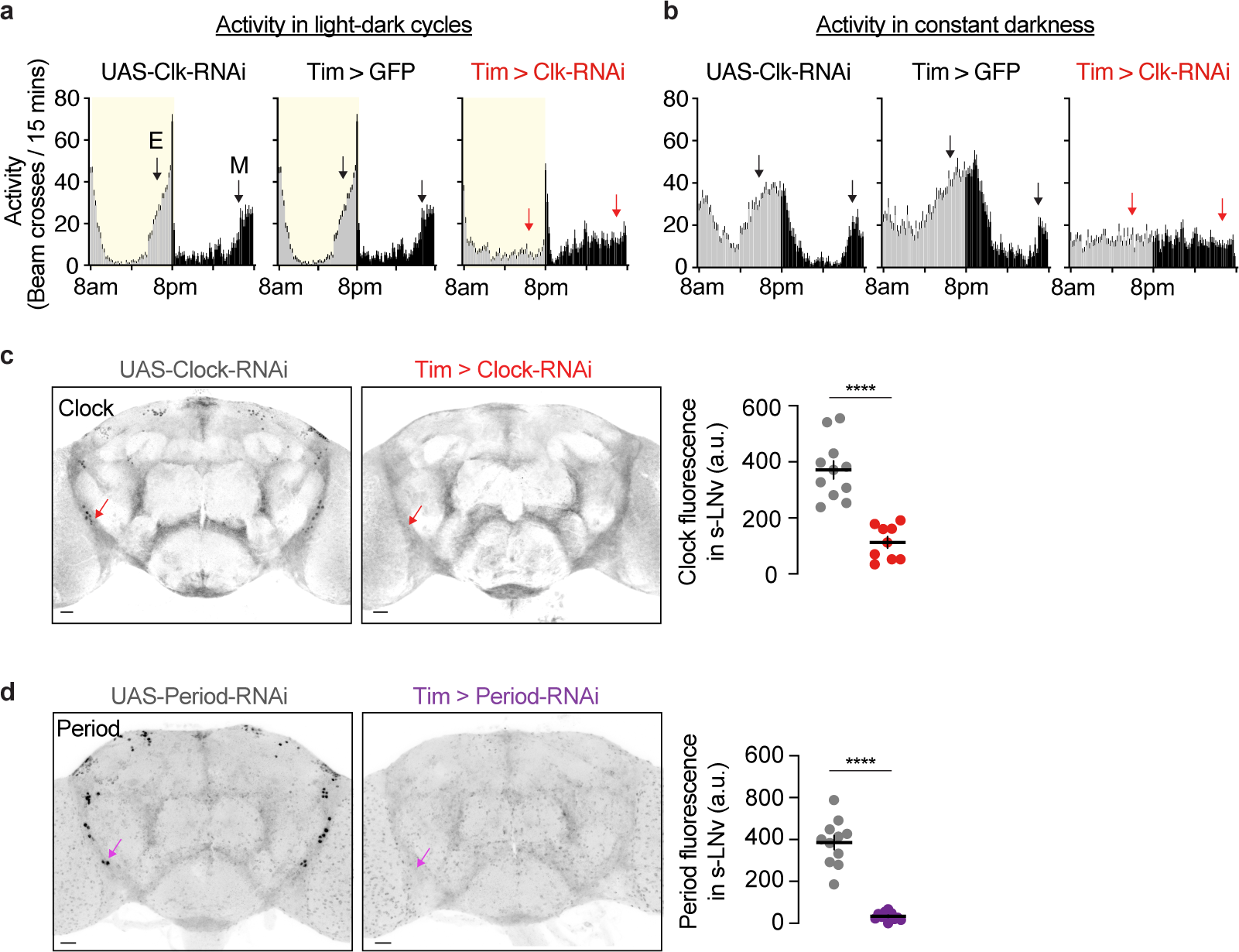
Testing the efficacy of Clock- and Period- RNAi. **(a)** 24-hour activity profiles in light-dark cycles show that targeting the core circadian protein Clock (Clk) by Tim-Gal4- driven RNAi prevents anticipatory locomotor activity (i.e. the ramp-up of activity that occurs prior to lights turning on (morning anticipation, M) or off (evening anticipation, E). Arrows indicate the presence (black) or absence (red) of anticipation. **(b)** 24-hour locomotor activity profiles in constant darkness show arrhythmicity when the clock is disrupted. **(c)** Evidence that Clk-RNAi effectively depletes Clock protein. Clock staining was arbitrarily performed at ZT6 since Clock protein levels are detectable throughout the day^79^. (**d**) Confirmation that Per-RNAi effectively depletes Per protein. Period staining was performed at 9am (Zeitgeber Time 1, ZT1), since Period expression oscillates in controls and is near peak levels during this time^80^. Period staining is broad because of glial expression^81, 82^.

**Extended Data Figure 5.**
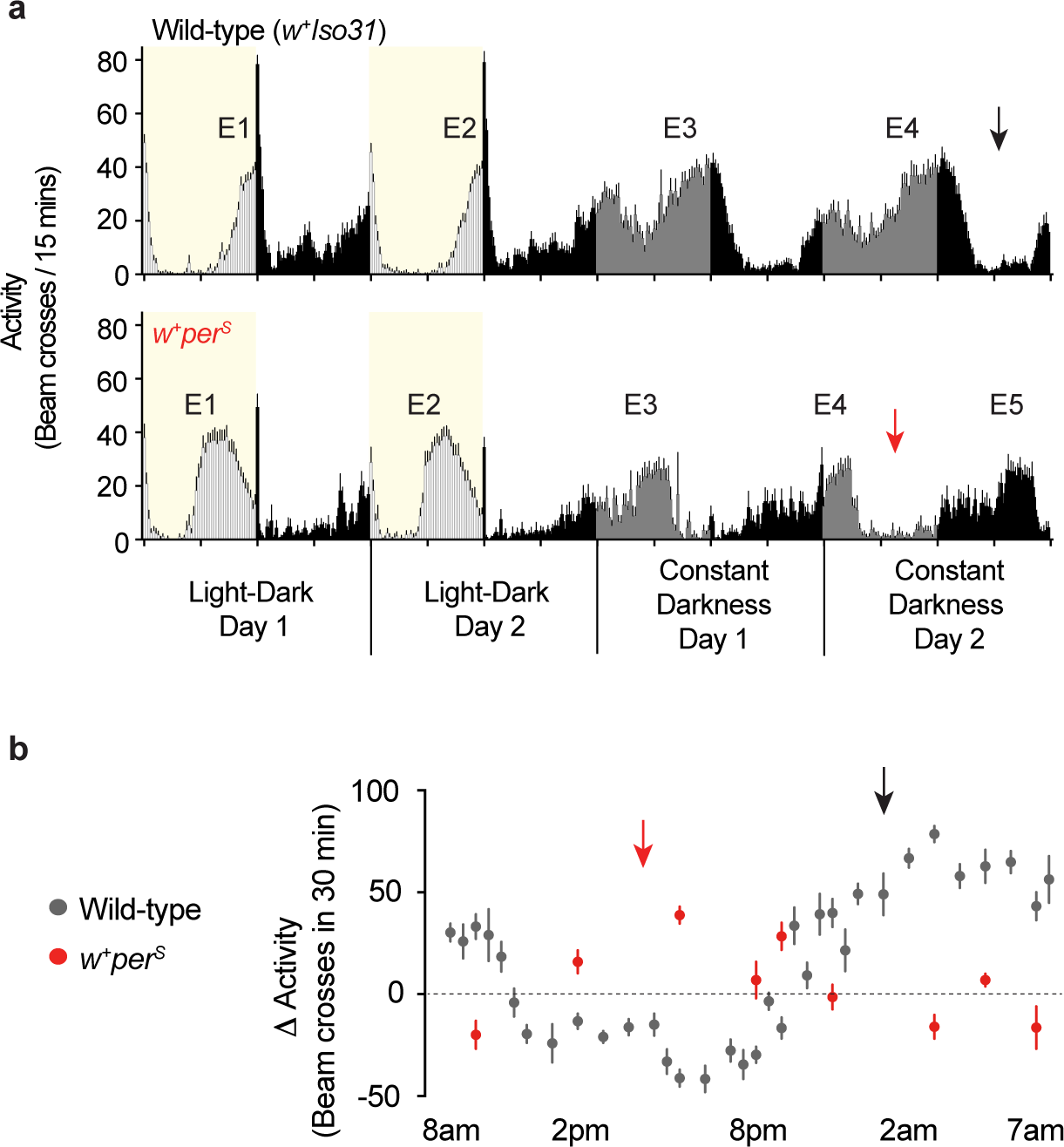
The timing of light responsiveness is controlled by circadian clocks. **(a)** A mutation in *period* (*period*^*Short*^ or *per^S^*) accelerates circadian rhythms. E# indicates corresponding peaks of evening activity between control and mutant flies. Arrows indicate comparable timepoints - subjective nighttime after the fourth peak of evening activity. **(b)** This mutation also accelerates the timecourse of light responsiveness. Same control flies as in Fig. 1c.

**Extended Data Figure 6.**
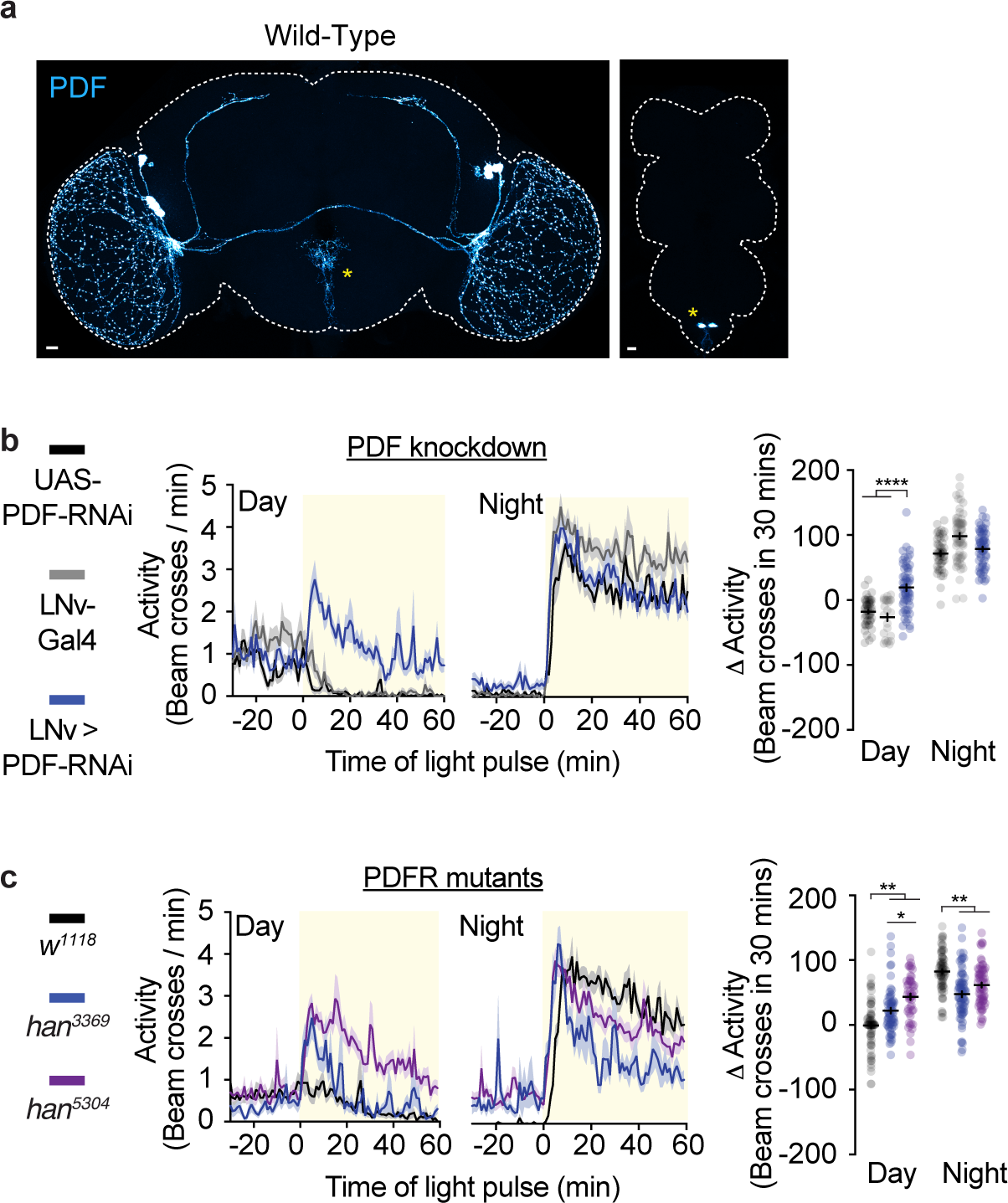
PDF in LNVs is required for normal daytime, but not nighttime, light reactivity. **(a)** Expression pattern of PDF in the brain and VNC. Asterisks indicate variable expression in the midline of the brain and non-circadian neurons^83^ in the VNC. **(b)** Knockdown of PDF in LNvs perturbs only daytime light responsiveness. **(c)** Two hypomorphic mutations of PDF receptor (PDFR) show perturbed responsiveness to daytime light. Though nighttime light responses differ from controls, peak responses to nighttime light are similar to wild-type.

**Extended Data Figure 7.**
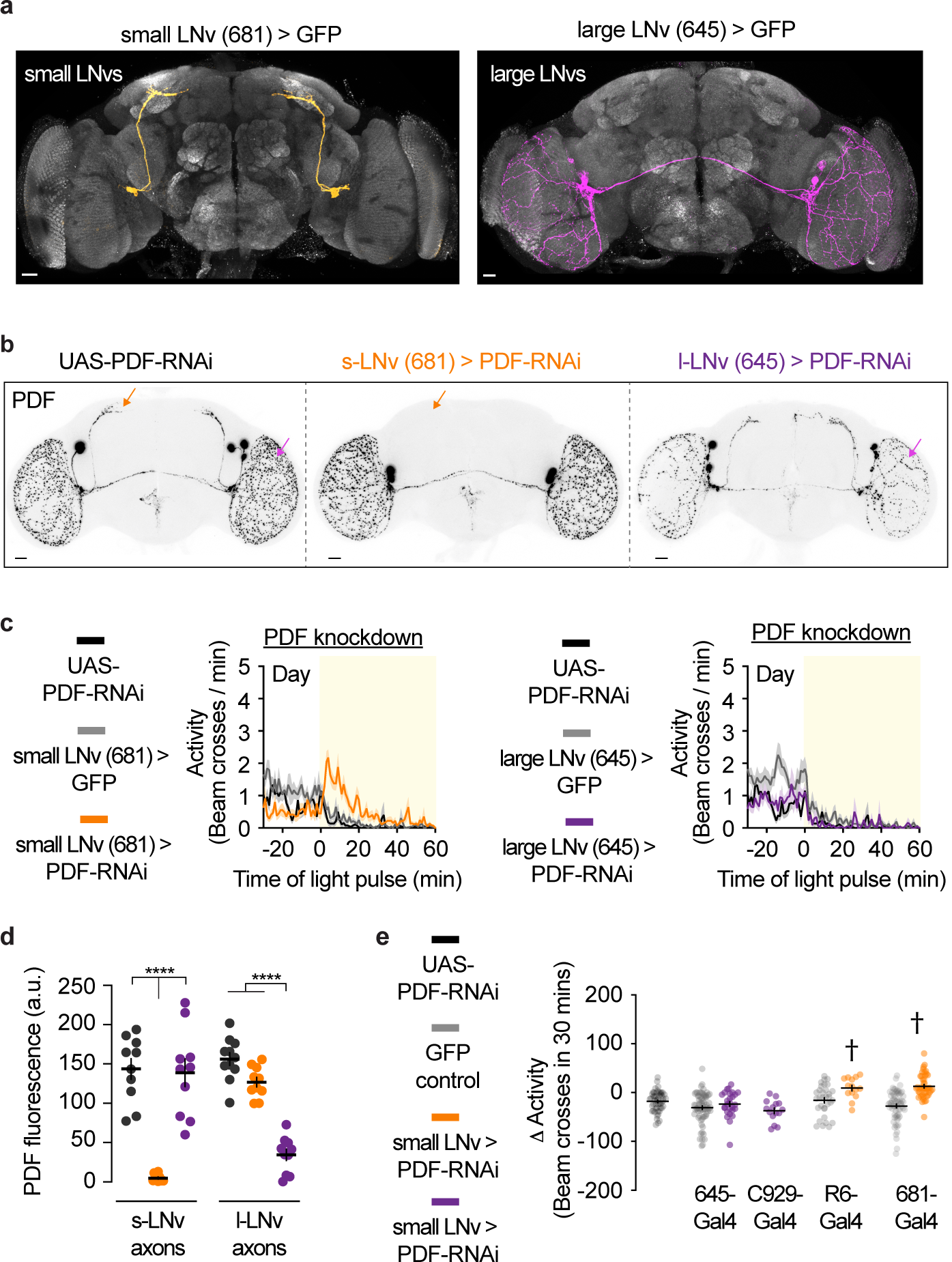
PDF from small LNvs is required for normal daytime light reactivity. **(a)** Expression patterns of Gal4 drivers in small (JRC_SS00681-Gal4) or large (JRC_SS00645-Gal4) LNv subpopulations. **(b)** Representative PDF expression patterns when PDF-RNAi was driven in either small or large LNv subpopulations. Arrows indicate quantified regions in (d). Note that PDF was not completely depleted by l-LNv-Gal4-driven RNAi. **(c)** PDF knockdown shows that small LNvs (681-Gal4) are a necessary source of daytime PDF. UAS- PDF-RNAi controls (black) are duplicated between left and right panels. **(d)** Quantification of PDF immunostaining intensity in s-LNv dorsal axons or l-LNv axons in the optic lobe. **(e)** Light-evoked change in activity using two s-LNv and two l-LNv drivers to express PDF-RNAi. One- way ANOVA of daytime light responses, with Tukey’s post hoc test. Cross symbols within panel indicate significance (p<.05) versus all other conditions, aside from each other.

**Extended Data Figure 8.**
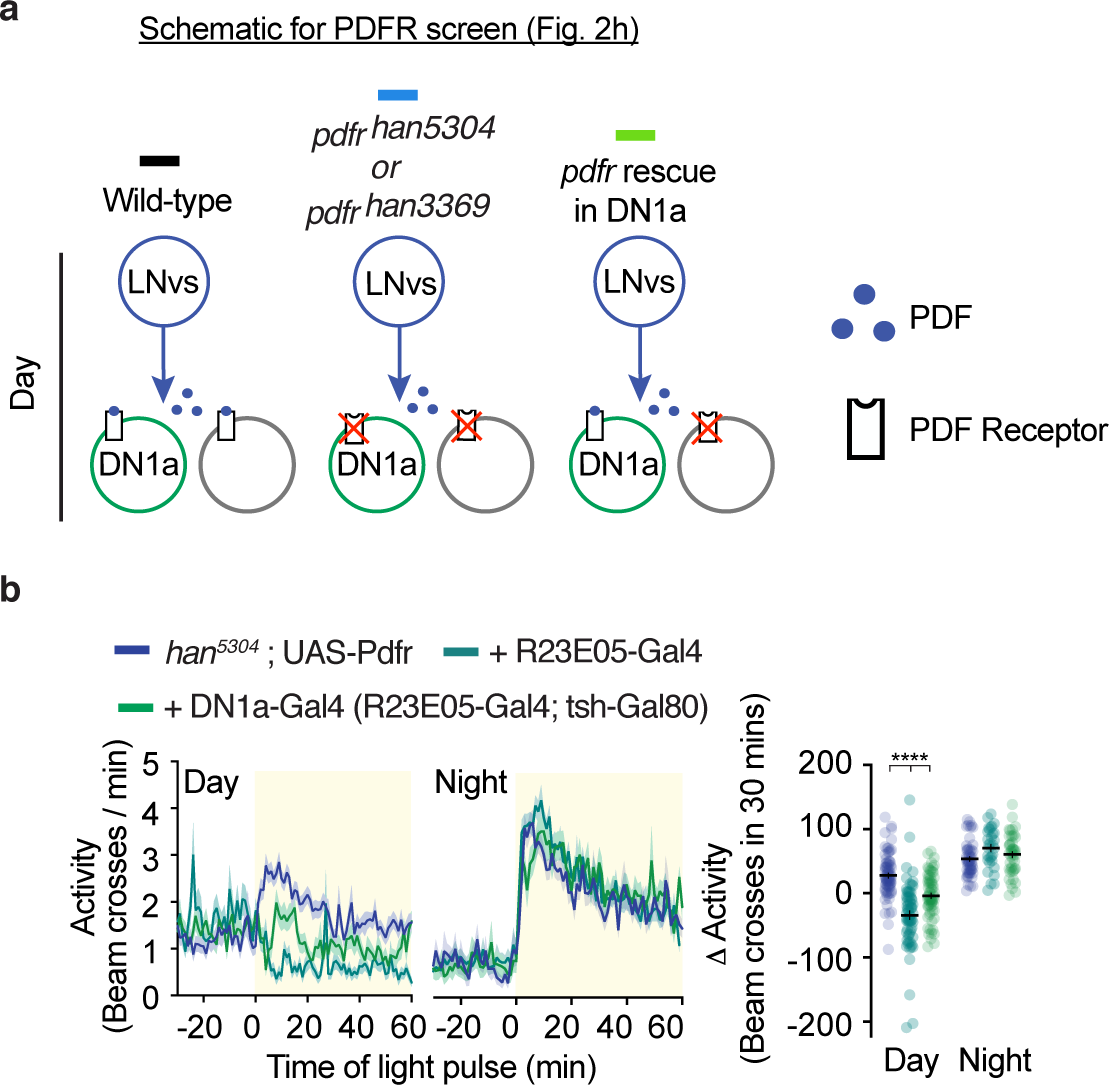
Additional details about PDFR rescue screen. **(a)** Schematic of experiment in Fig. 2d. **(b)** Activity traces for flies in which PDFR was expressed using R23E05- Gal4 (dark green) or DN1a-Gal4 (light green) in the *han*^**5304**^ *pdfr* mutant background. This experiment, done in the *han^3369^ pdfr* mutant background is shown in Fig. 2d.

**Extended Data Figure 9.**
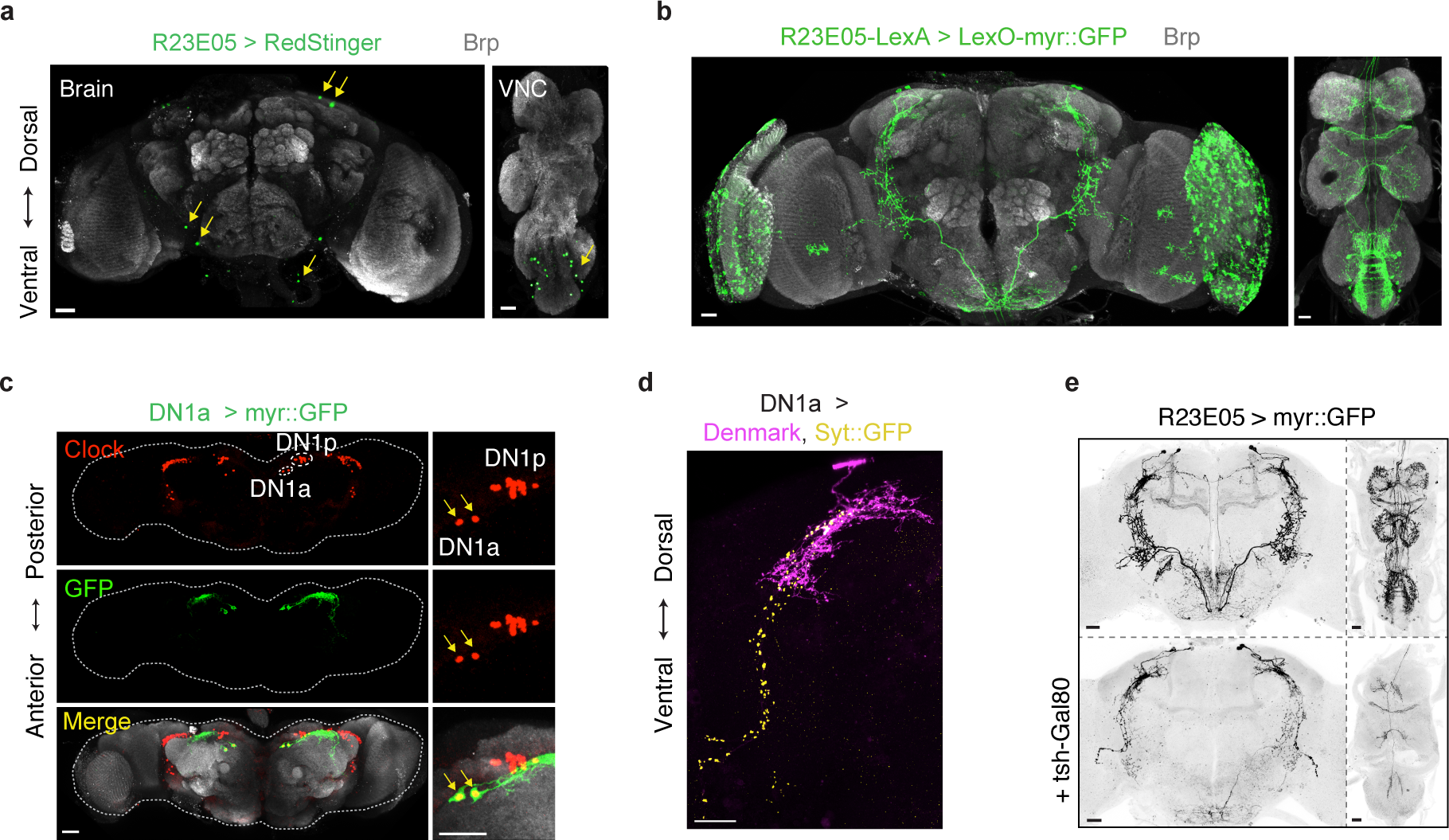
Additional anatomical and behavioral characterization of R23E05-Gal4. **(a)** R23E05-Gal4 driving the nuclear reporter RedStinger. The nervous system expression includes 2 DN1as, 2 neurons in the saddle, 1 cell in the inferior posterior slope, and ∼10 neurons in the ventral nerve cord, per hemisphere. With some reporters, we saw weak expression in the mushroom body. **(b)** Expression pattern of R23E05-LexA. We saw strong expression in the lamina, which was not seen with R23E05-Gal4. **(c)** Dorsal view of DN1a-Gal4 expression pattern confirms expression in anterior, and not posterior, DN1 subpopulations. **(d)** DN1a-Gal4 driving markers of postsynaptic sites (Denmark) and presynaptic sites (Synaptotagmin:::GFP, Syt::GFP). **(e)** Restriction of R23E05 primarily to DN1as using teashirt- Gal80 (tsh-Gal80). Expression was diminished in most VNC neurons and in some central brain neurons.

**Extended Data Figure 10.**
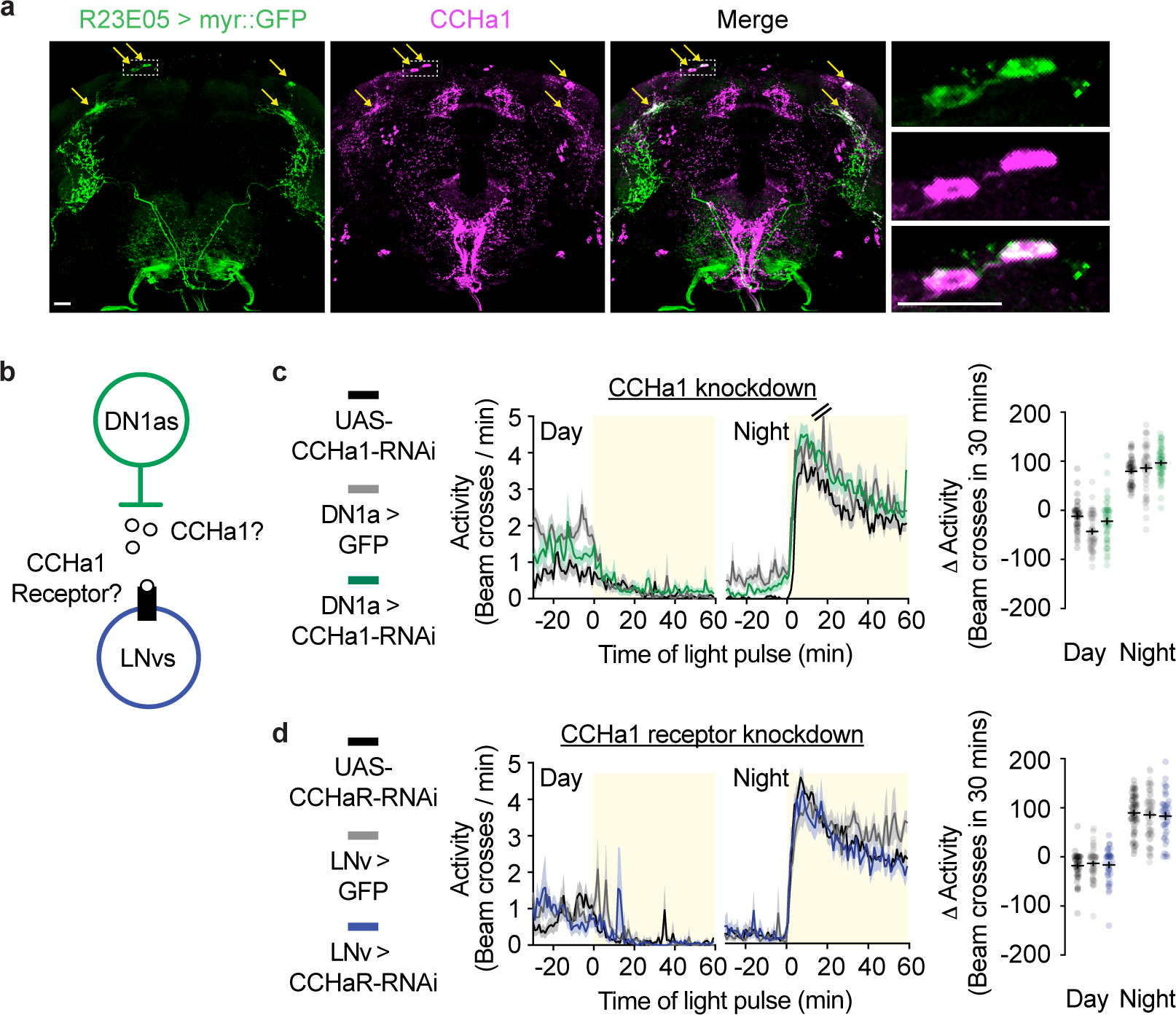
The peptide CCHa1 is not the relevant nighttime DN1a-to-LNv signal. **(a)** Co-staining of CCHa1with GFP in DN1as. **(b)** Schematic of DN1a-to-LNv communication via CCHa1. **(c)** RNAi against CCHa1 in DN1as does not mimic the effects of DN1a silencing. **(d)** RNAi against CCHa1 receptor in LNvs does not mimic the effects of DN1a silencing.

**Extended Data Figure 11.**
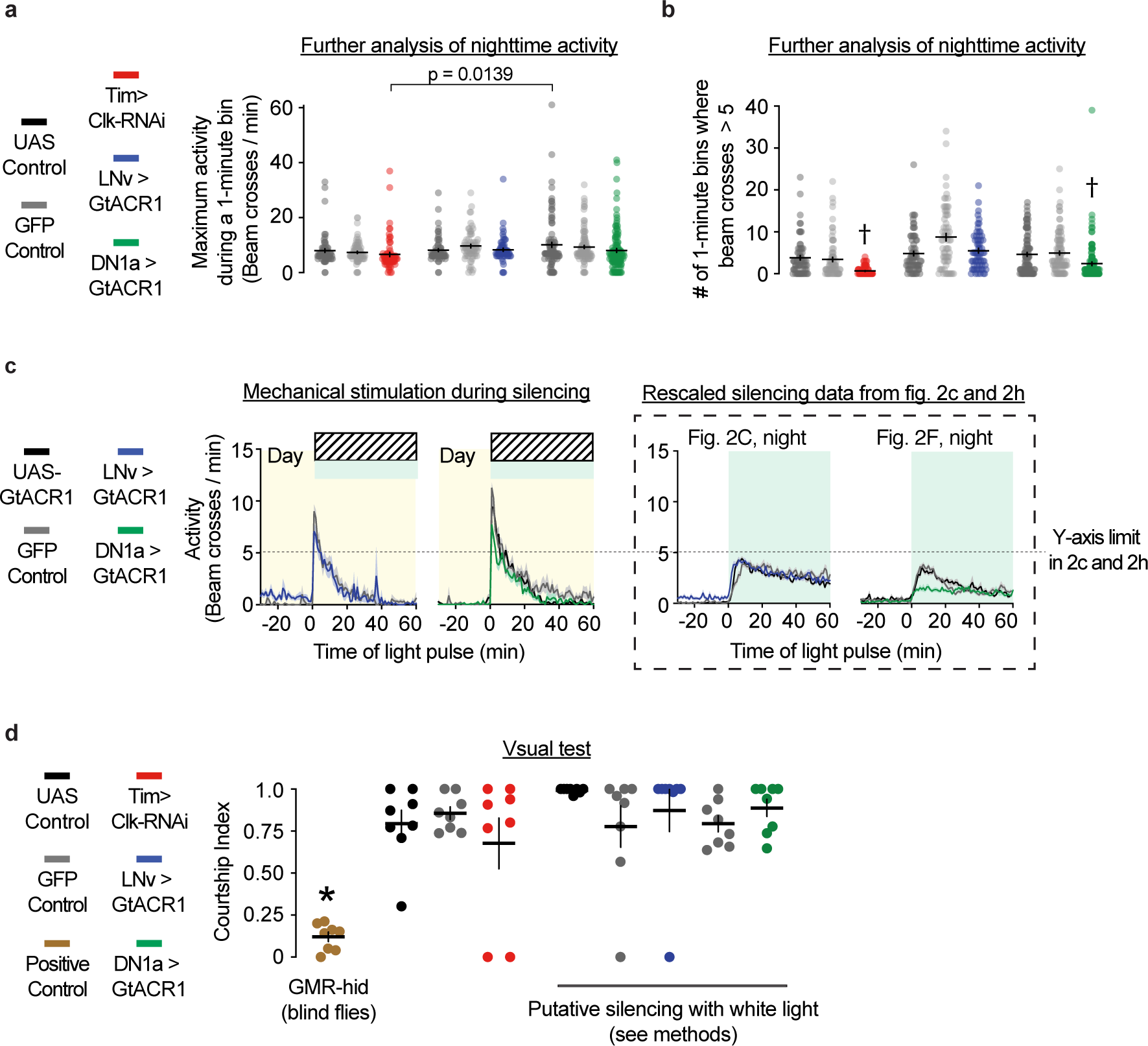
Manipulating LNv or DN1a activity does not cause general visual or locomotor defects. **(a)** Left, peak locomotor activity per 1-minute bin, during nighttime light pulses (same flies as Fig. 1g) or during nighttime optogenetic silencing (same flies as Fig 2c and Fig. 2h). Right, number of high activity bouts (animals cross the middle of the tube more than 5 times per 1-minute bin). Cross symbols within panel indicate significance against all other conditions (except against each other). The LNv > GFP control genotype had significantly more high-activity bouts than all other conditions, which is not indicated in the figure. One-way ANOVA with Tukey’s post hoc test. (**G)** Mechanical stimulation of LNv-silenced and DN1a-silenced flies shows that all genotypes are capable of reaching high levels of locomotion. **(H)** Courtship of clock-disrupted flies and subpopulation-silenced flies, which is compared to GMR-hid flies, a positive control with visual defects. See methods for details about optogenetics during courtship.

**Extended Data Figure 12.**
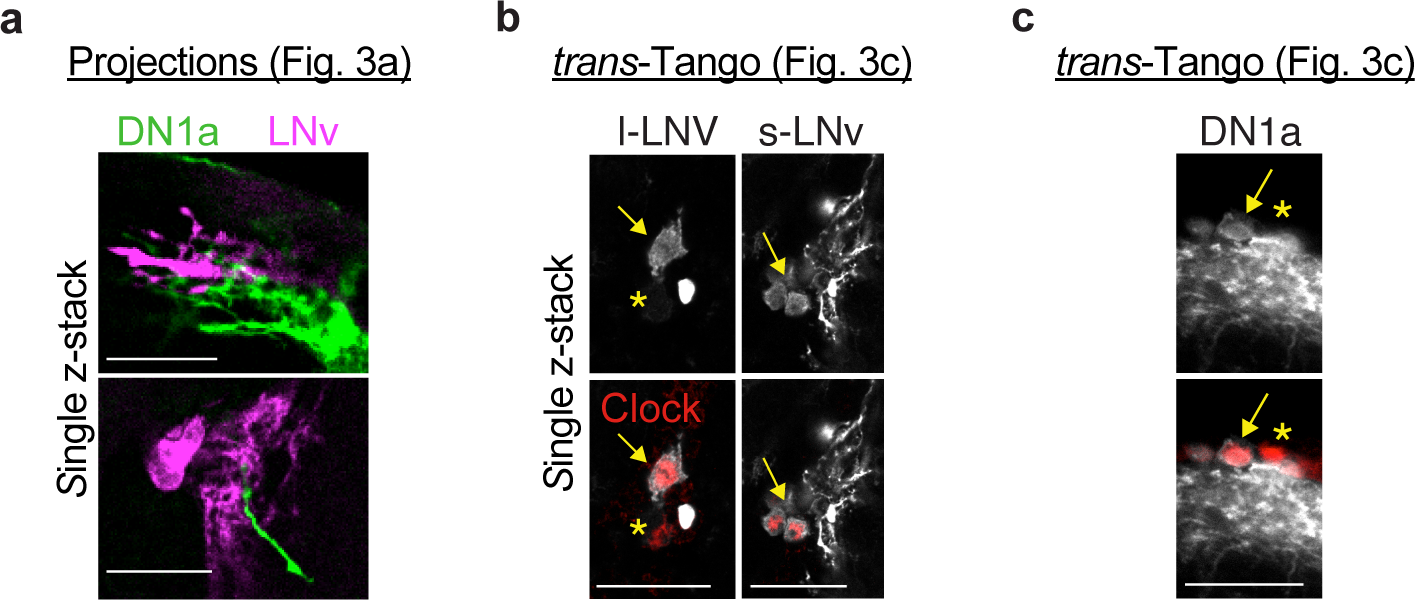
Additional data for mutual connectivity experiments. **(a)** Single confocal z-stacks with imaging depths of 0.3 µm show proximity of LNv and DN1a terminals **(b)** Single confocal z-stacks show specificity of DN1a->LNv *trans*-Tango experiments. **(c)** Single confocal z-stacks show specificity of LNv->DN1a *trans*-Tango experiments.

**Extended Data Figure 13.**
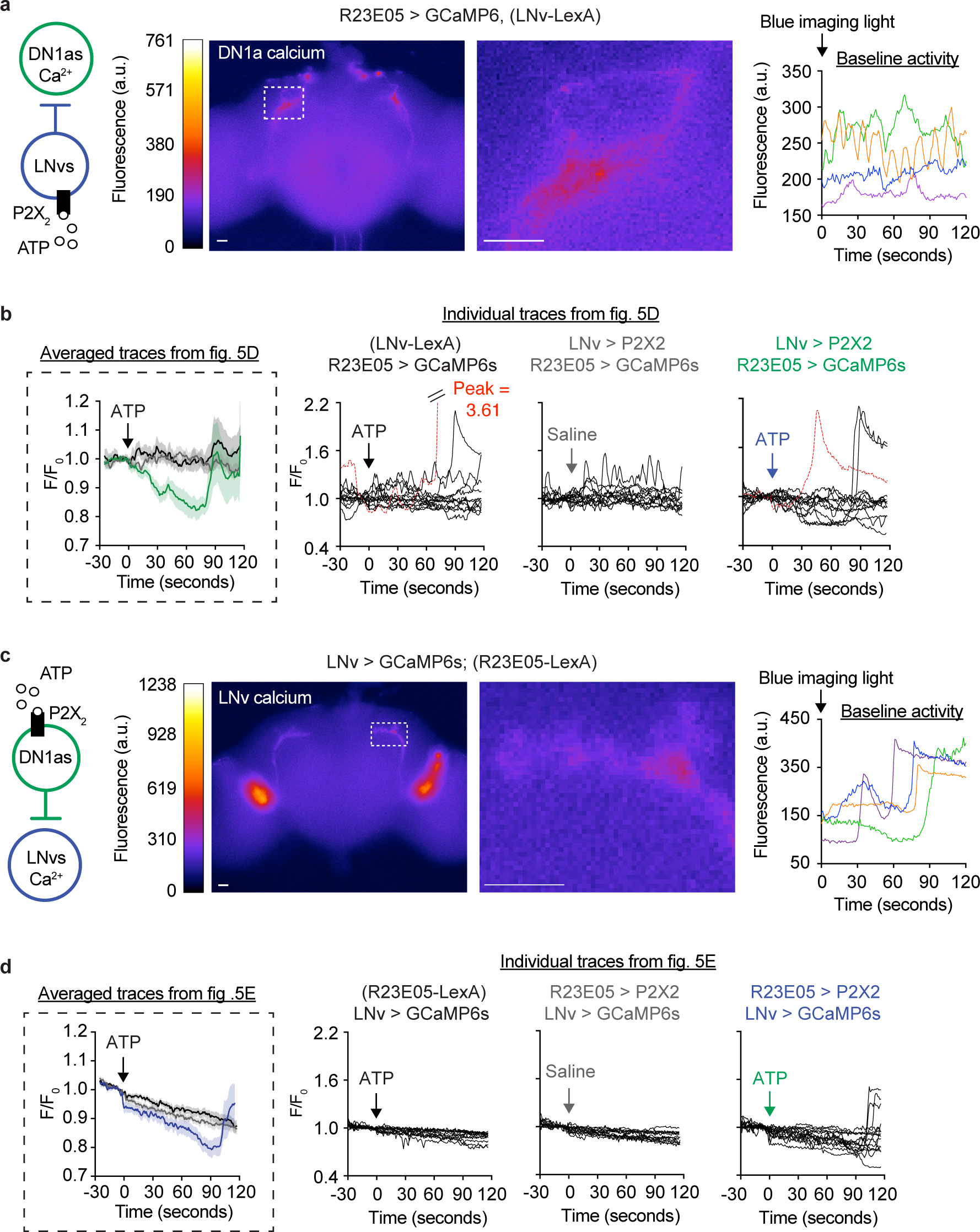
Additional data for functional imaging. **(a)** Left, representative fields of view during LNv-to-DN1a circuit tracing experiment. Dashed box shows magnified DN1a dendritic region selected for analysis. DN1a dendrites and LNv axons were chosen for analysis because these regions were consistently identifiable, whereas DN1a axons and LNv dendrites were not usually visible with GCaMP6s. Left, representative baseline calcium fluctuations in DN1as before ATP stimulation. Right, representative calcium fluctuations in DN1a dendrites during two minutes of baseline imaging before ATP stimulation. Each color shows an independent trial. **(b)** All trials for experiments shown in Fig. 3d. Red traces are one experimental and one control trial that were excluded from analysis because of non- representative depolarizations (see methods for more details). **(c)** Left, representative fields of view during DN1a-to-LNv circuit tracing experiments. Inset represents LNv axonal area selected for analysis. Dashed box shows magnified region of analysis in LNv axons. Right, representative calcium fluctuations in LNv axons during two minutes of baseline imaging before ATP stimulation. Each color shows an independent trial. **(d)** All trials for experiments shown in Fig. 3e.

**Extended Data Figure 14.**
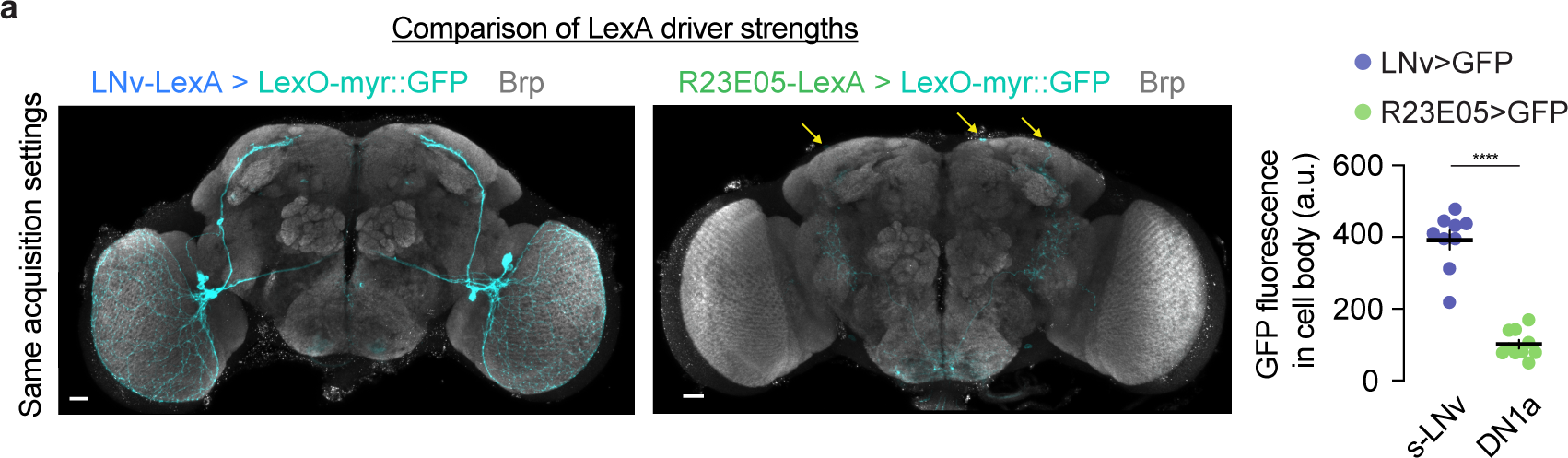
Comparisons of LexA driver strengths used for chemogentic stimulation in Figure 3. **(a)** LNv-LexA and DN1a-LexA driver strengths are compared by crossing to the same reporter (LexO::myrGFP), and imaged under the same acquisition settings GFP intensity was measured in cell bodies for comparison.

**Extended Data Figure 15.**
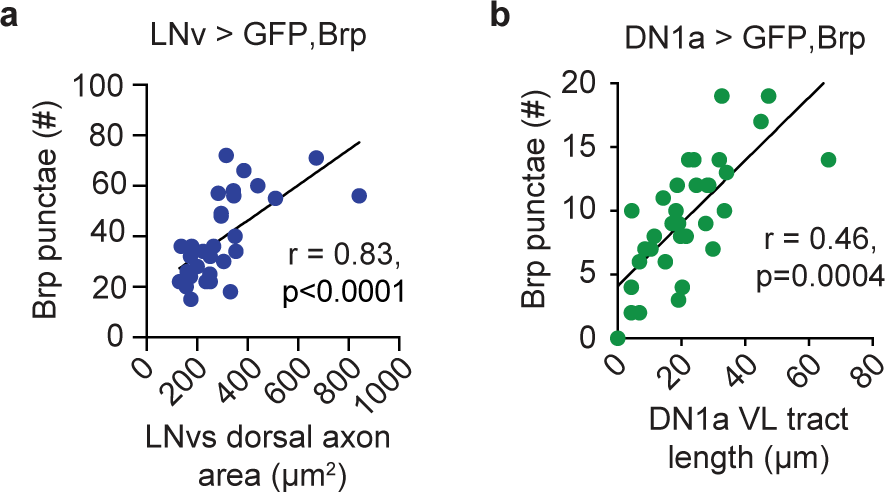
Additional information related to plasticity patterns. **(a)** s-LNv synapse number as a function of s-LNv axon area. **(b)** DN1a synapse number as a function of DN1a axon length. A and B are reanalyses of the same animals shown in Fig. 4b.

**Extended Data Figure 16.**
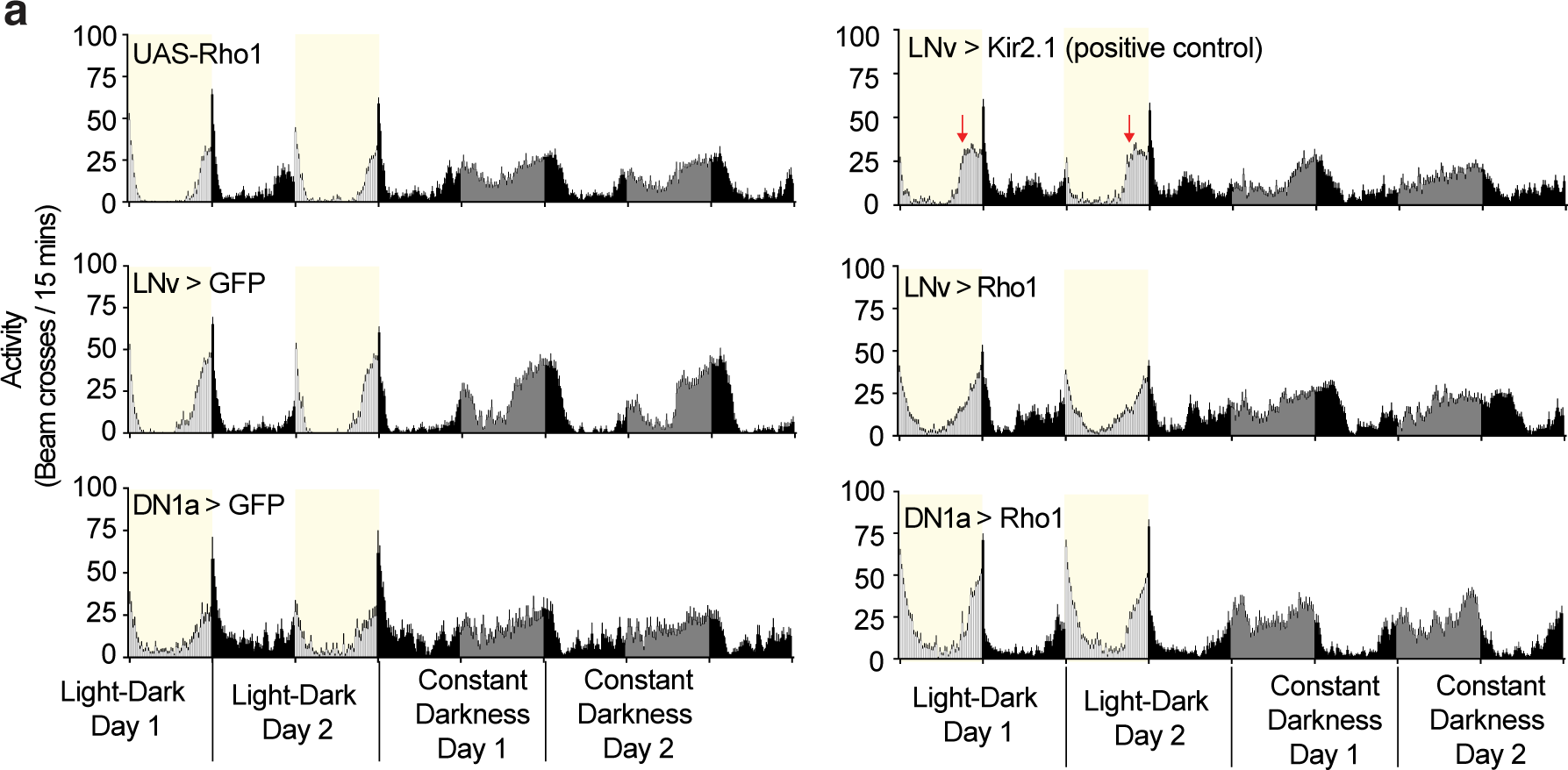
Additional information related to plasticity patterns. **(a)** Rho1 overexpression does not overtly perturb rhythmic locomotor activity. Locomotor activity of flies expressing Rho1 in LNvs or DN1as, across two days in light-dark cycles and two days in continual darkness. Red asterisks indicate advanced evening activity that occurs when LNvs are constitutively silenced using overexpression of inwardly rectifying potassium channel Kir2.1^84^.

**Extended Data Video 1. Circadian light reactivity in wild-type flies.** Light pulse presented to wild-type (*w^+^iso31*) flies. Flies are in glass tubes with food on one end (outer end of the frame), and a cotton plug on the other (inner end of the frame). Lights turn on 13 seconds into the video. Flies in left and right columns are entrained to opposite cycles. On the left side, lights turn on in the middle of subjective day. On the right side, lights turn on in the middle of subjective night. Video is sped up 30x.

**Extended Data Video 2. Overlapping projections of LNvs (blue) and DN1as (green).** 3D reconstruction of confocal stacks using Imaris, shown with and without nc82 (grey). Same image as Fig. 3a.

**Extended Data Video 3. LNv axons (blue) adjacent to DN1a dendrites (green).** 3D reconstruction of confocal stacks using Imaris, shown with and without nc82 (grey). Same image as Fig. 3b, left.

**Extended Data Video 4. DN1a axons (green) adjacent to LNv dendrites (blue).** 3D reconstruction of confocal stacks using Imaris, shown with and without nc82 (grey). Same image as Fig. 3b, right.

**Extended Data Table 1.**
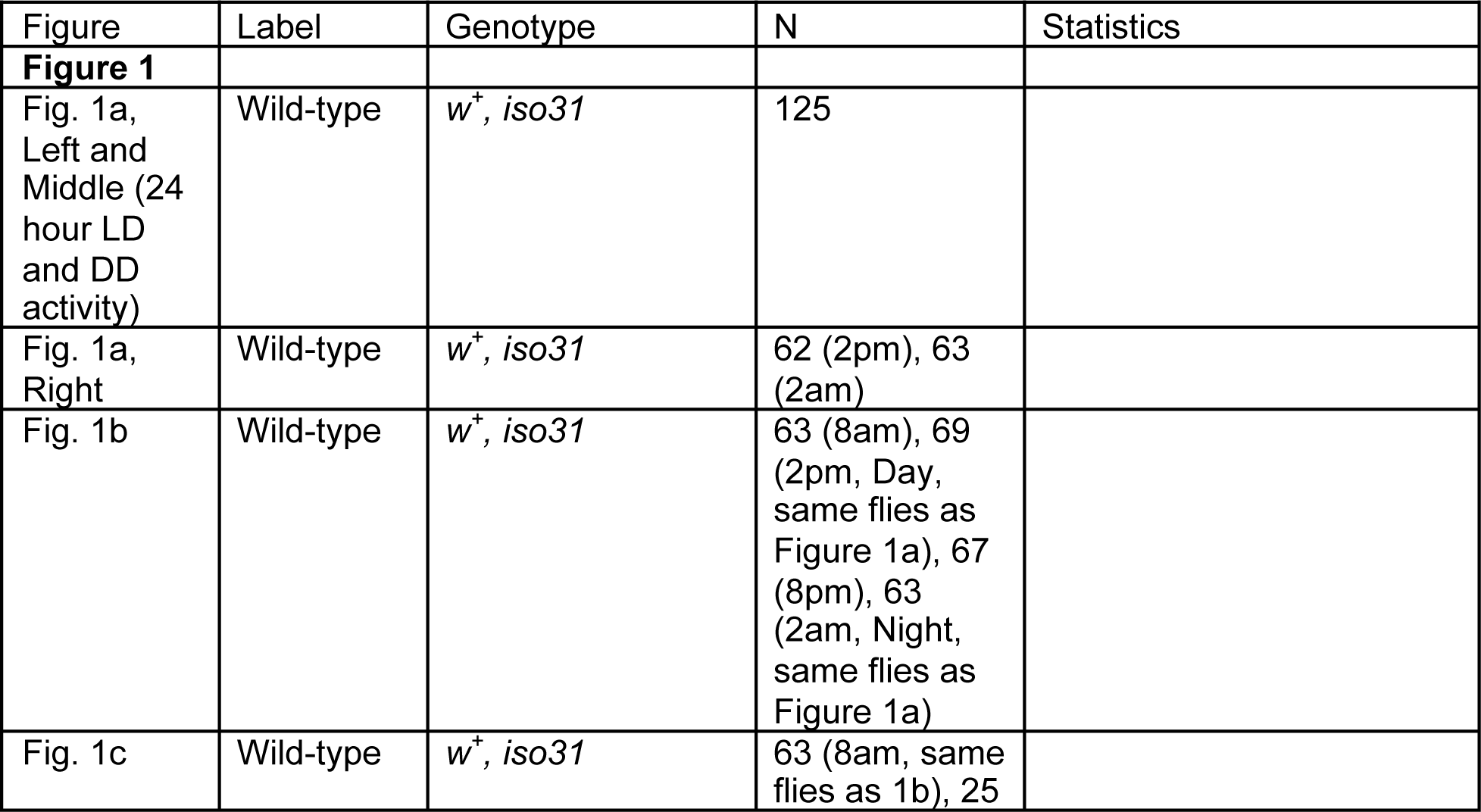
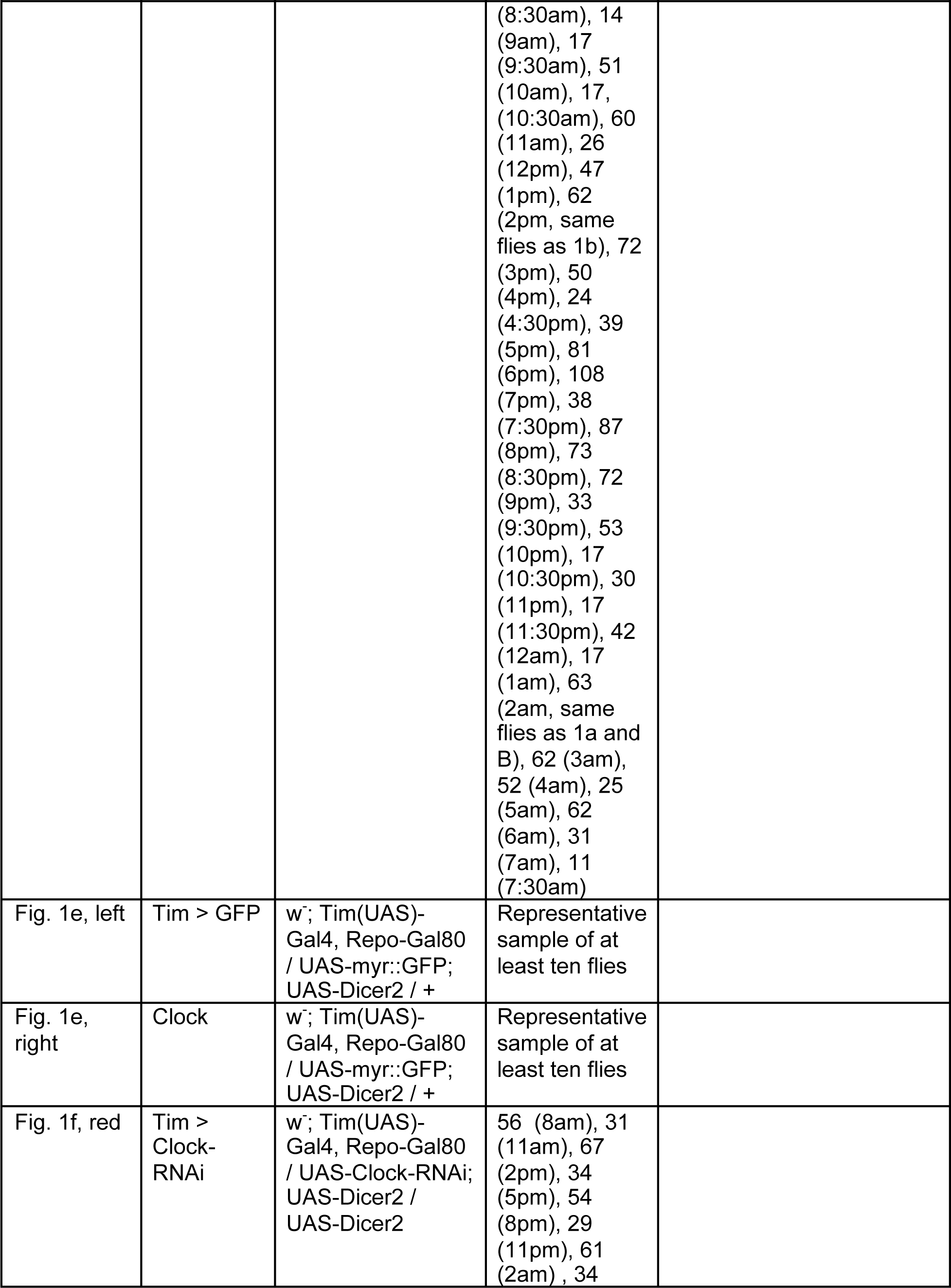
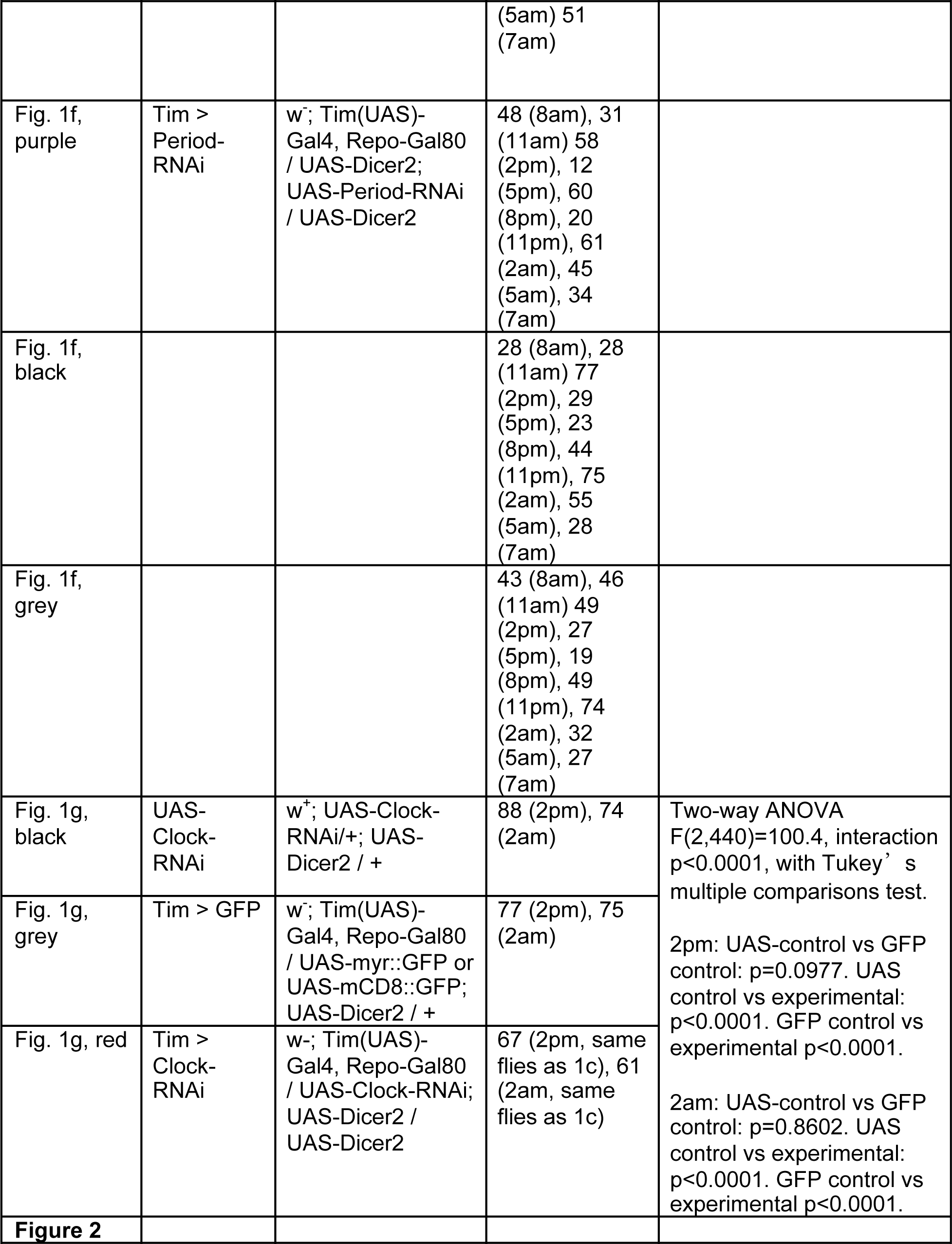
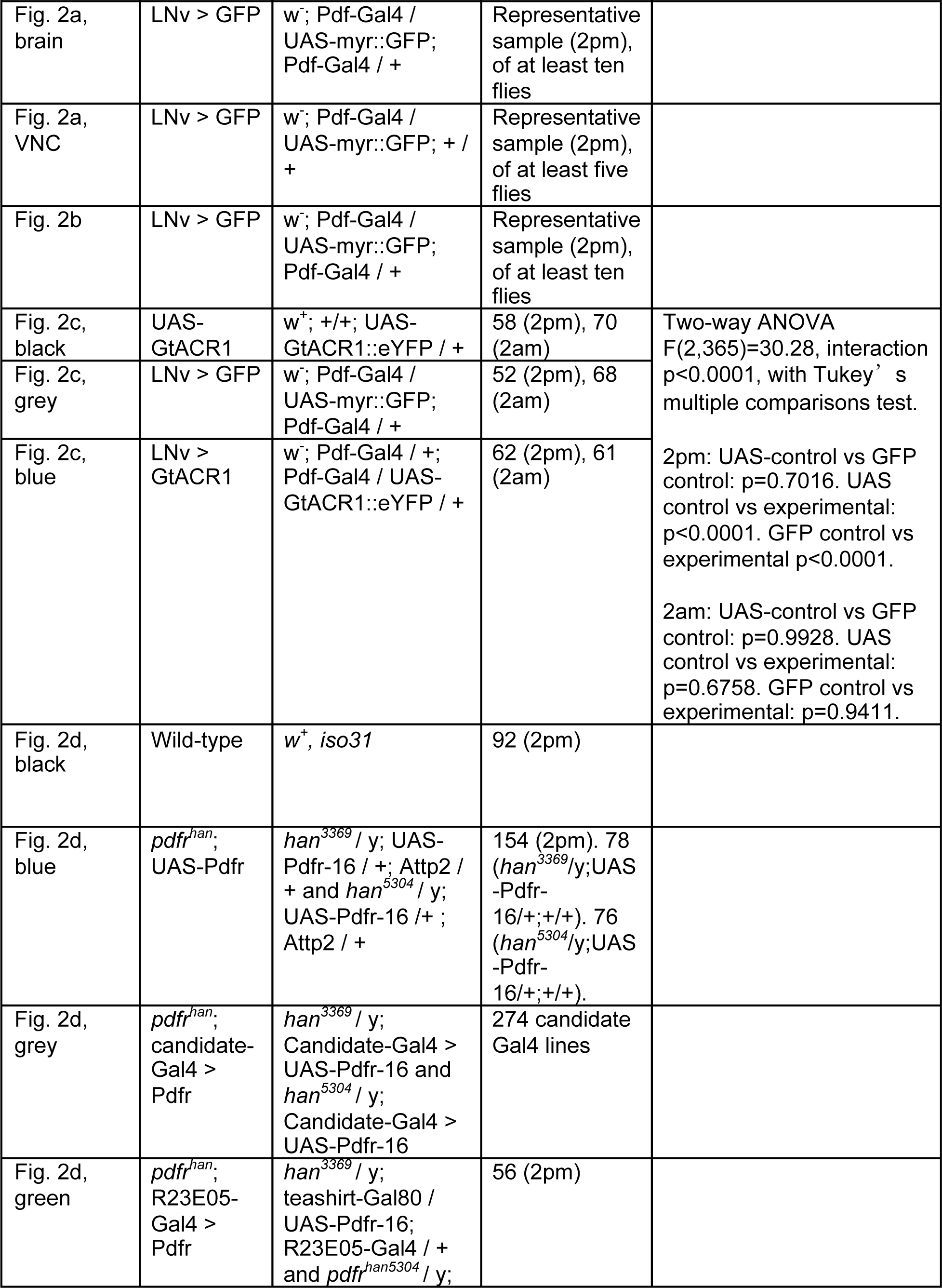
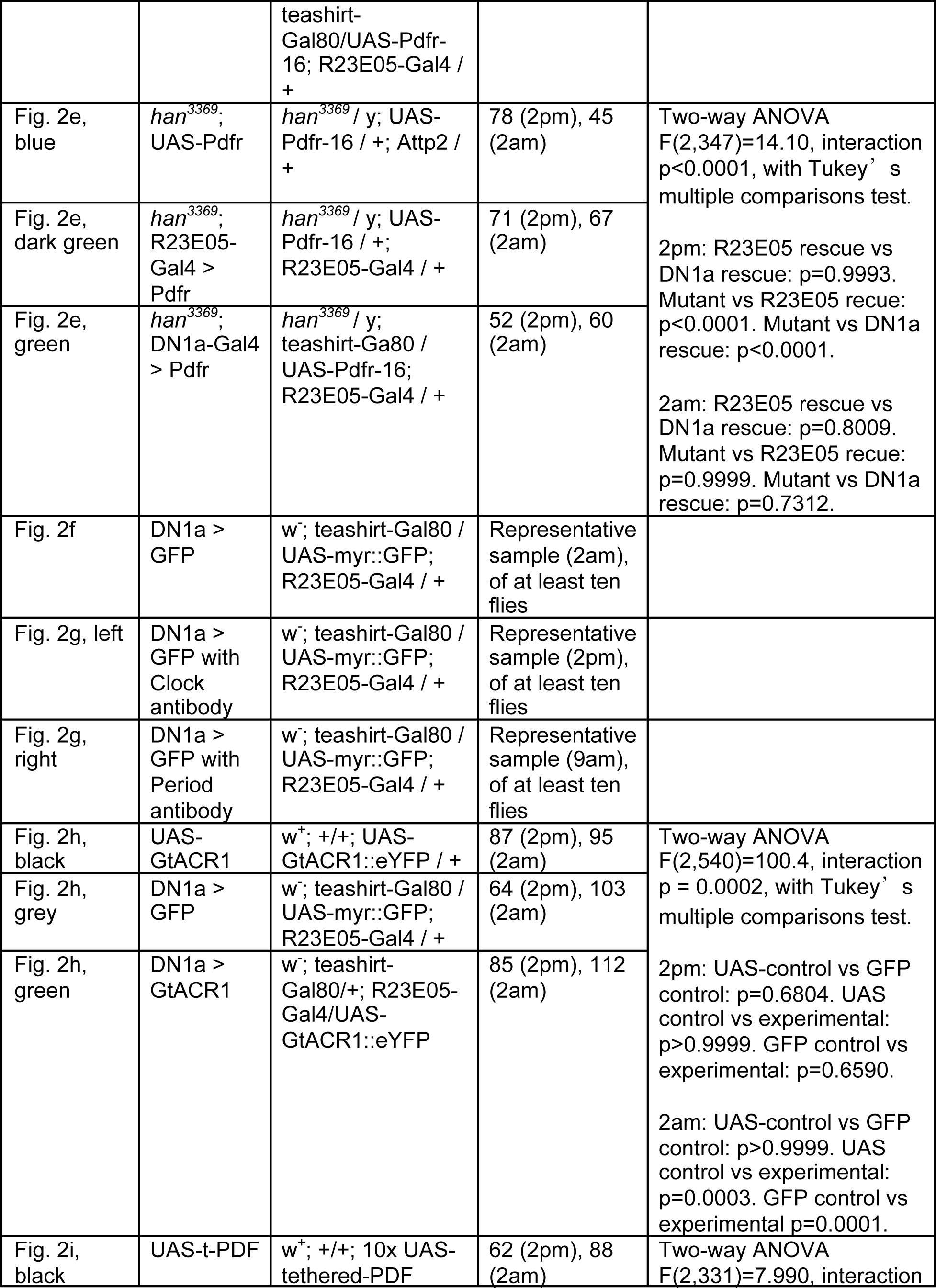
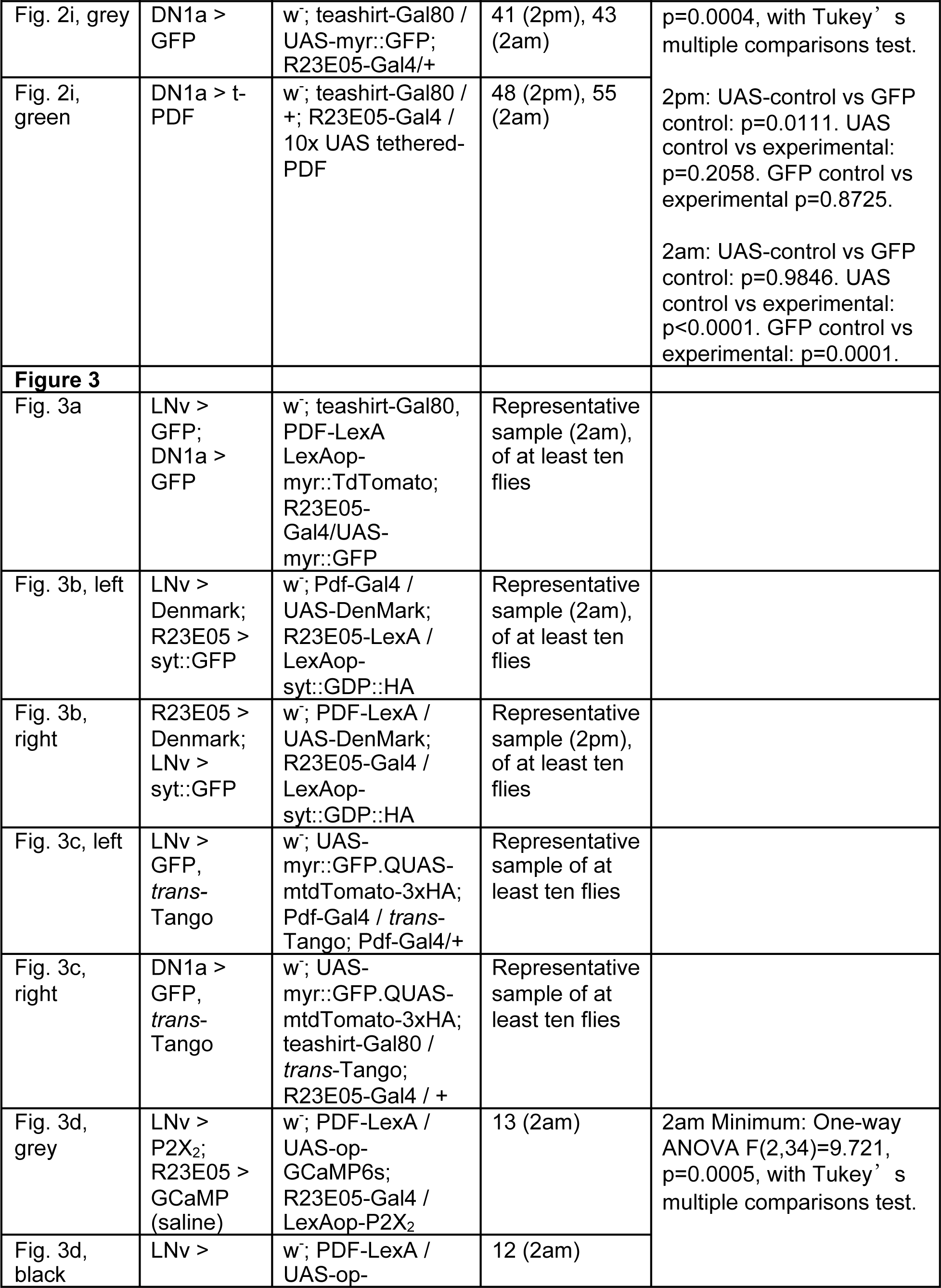
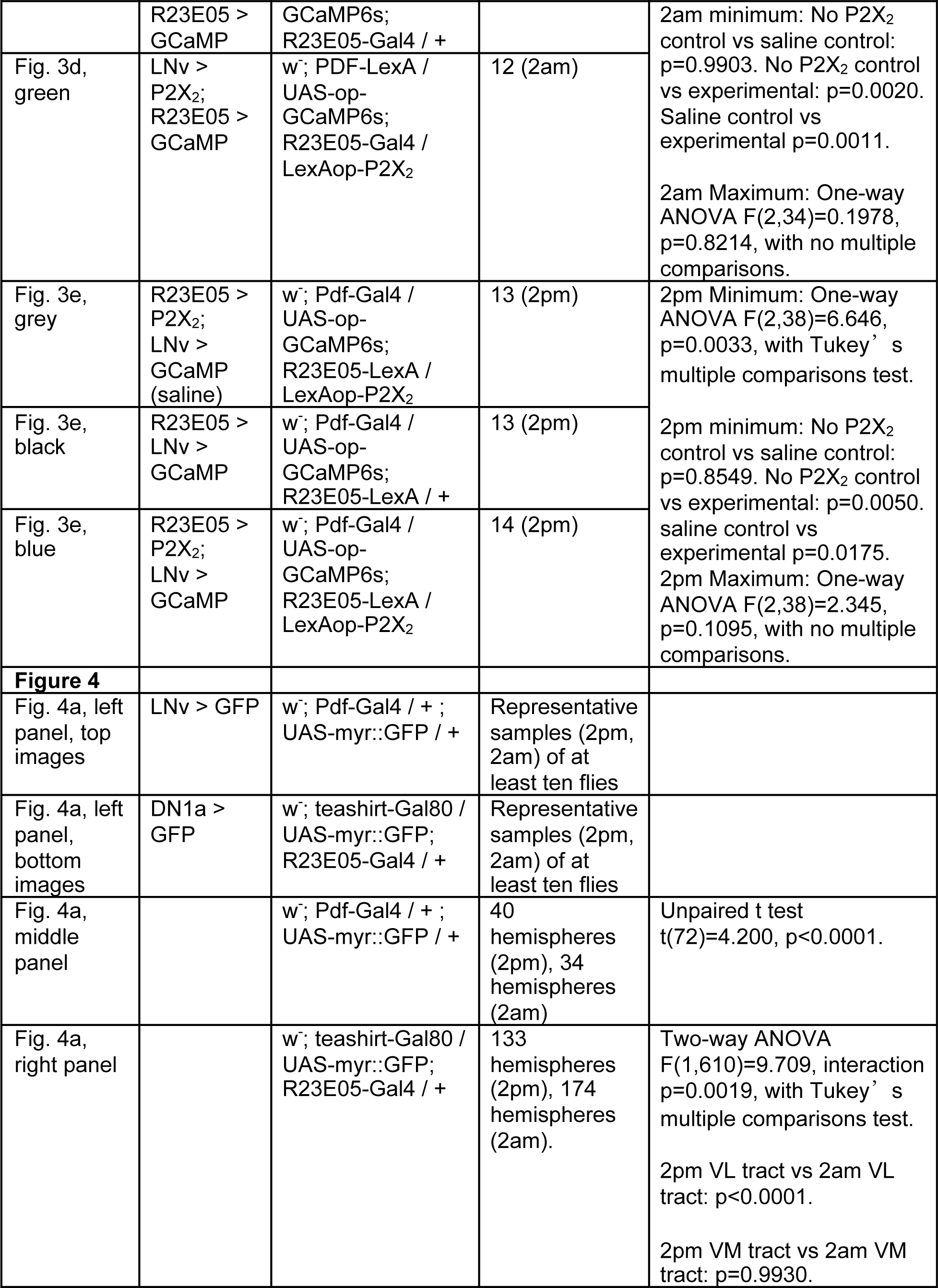
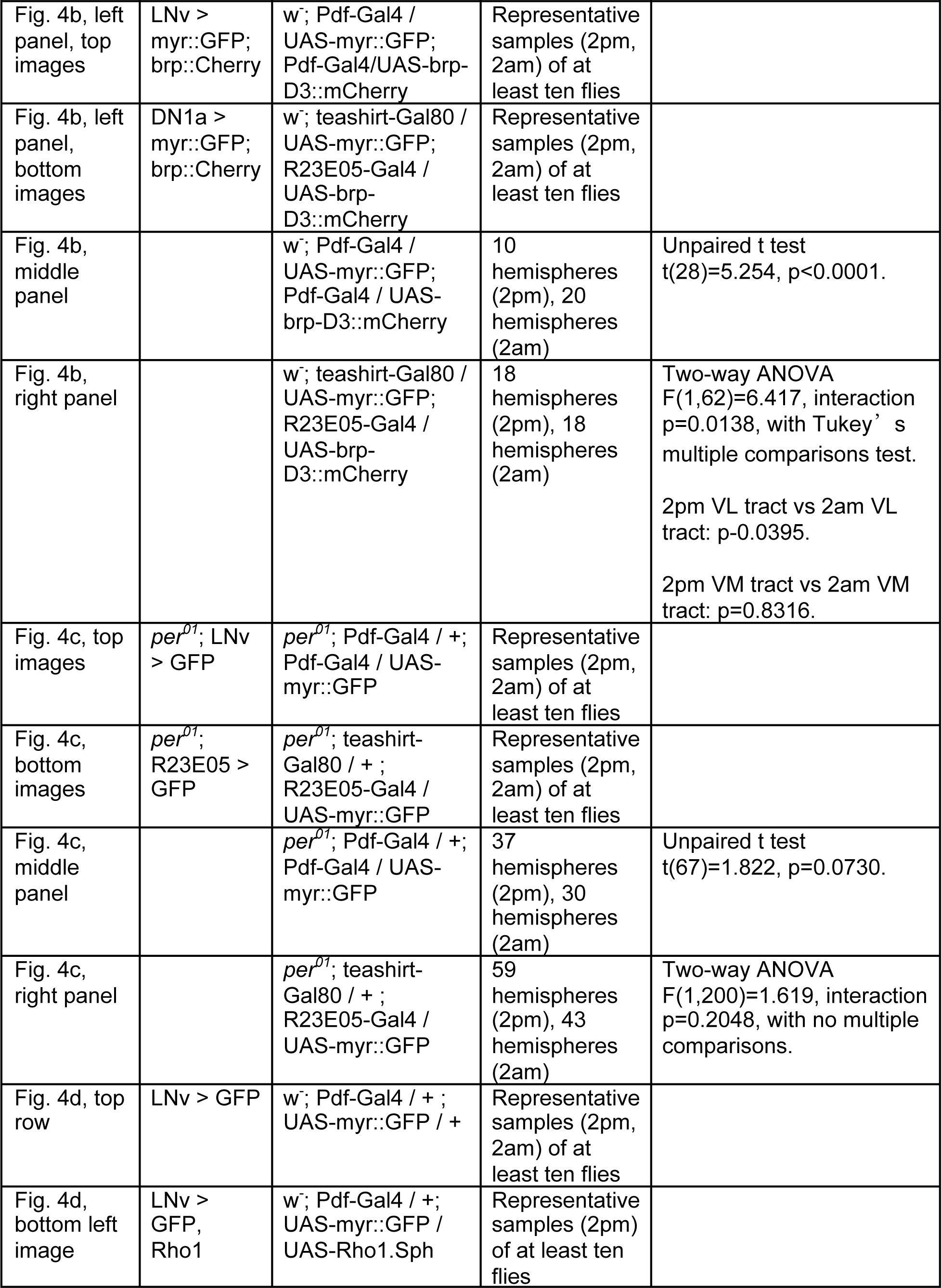
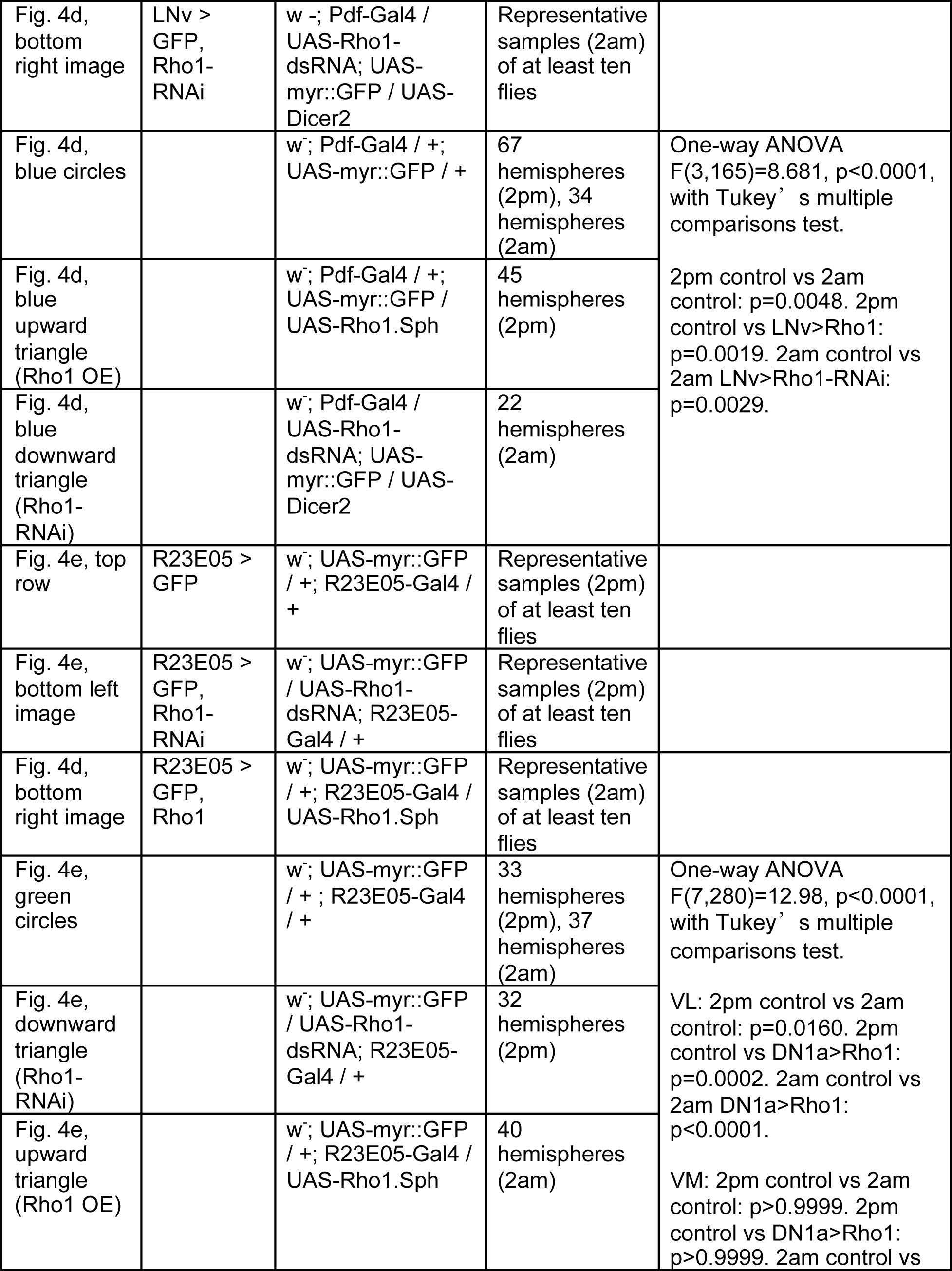
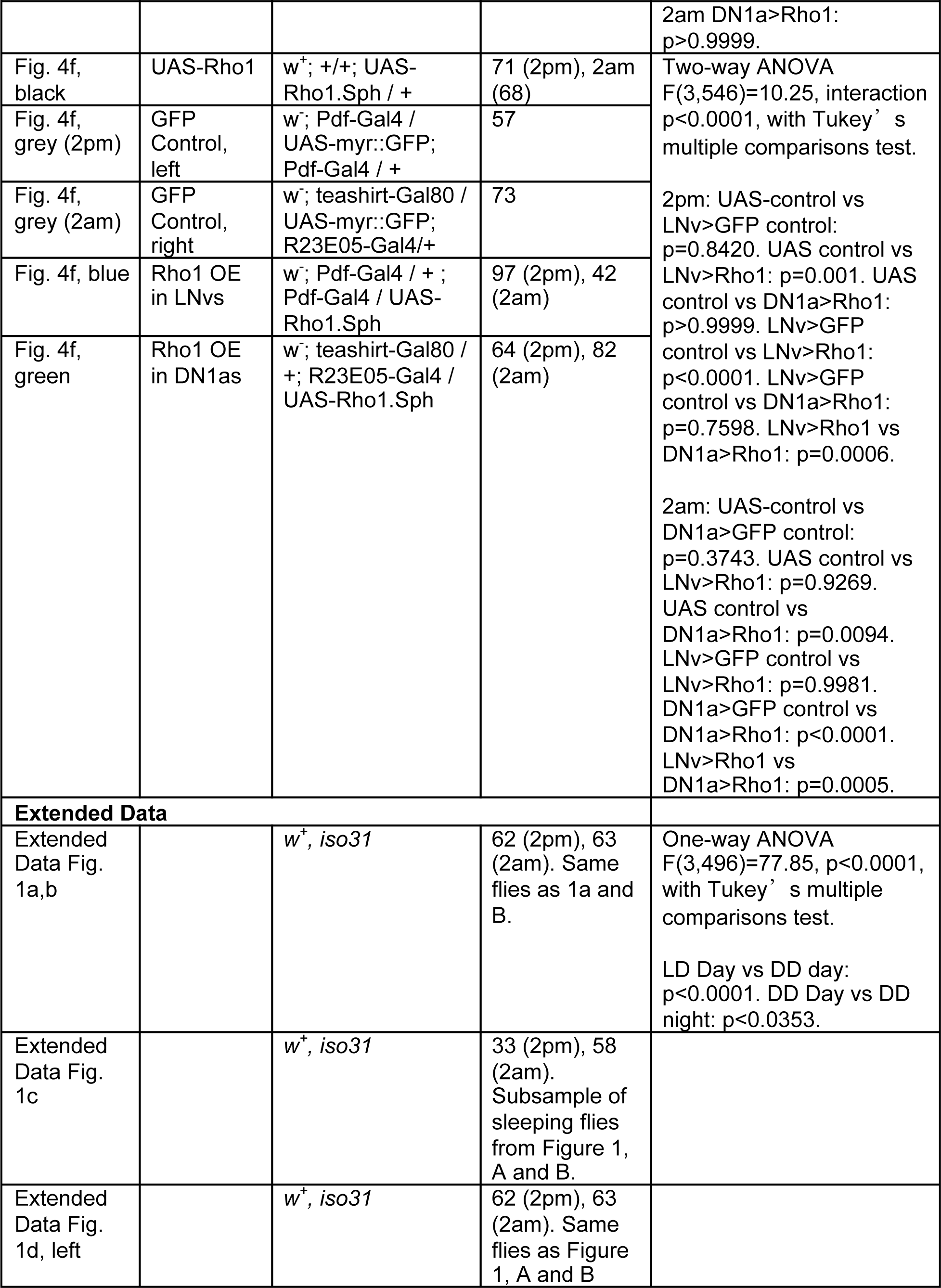
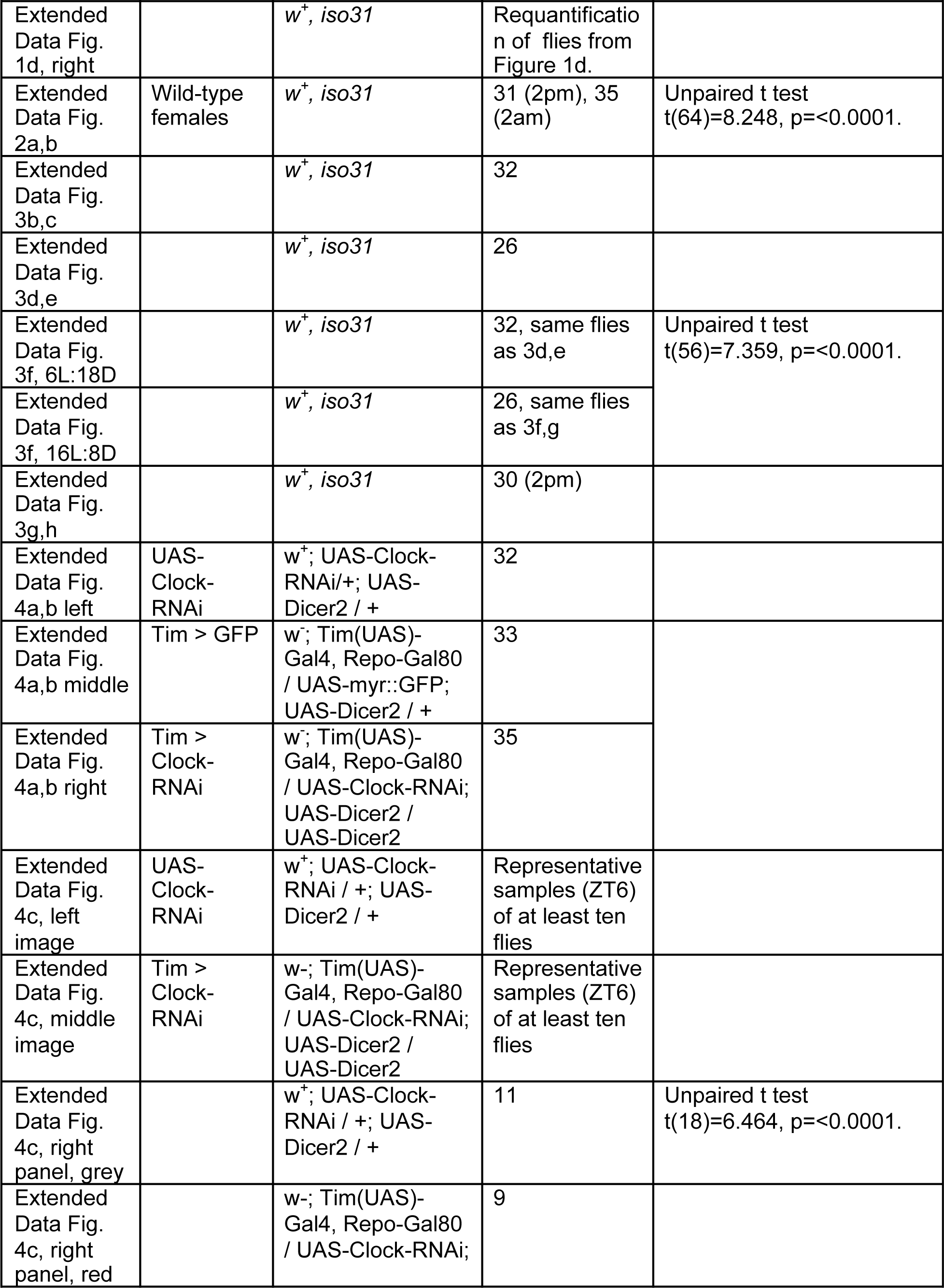
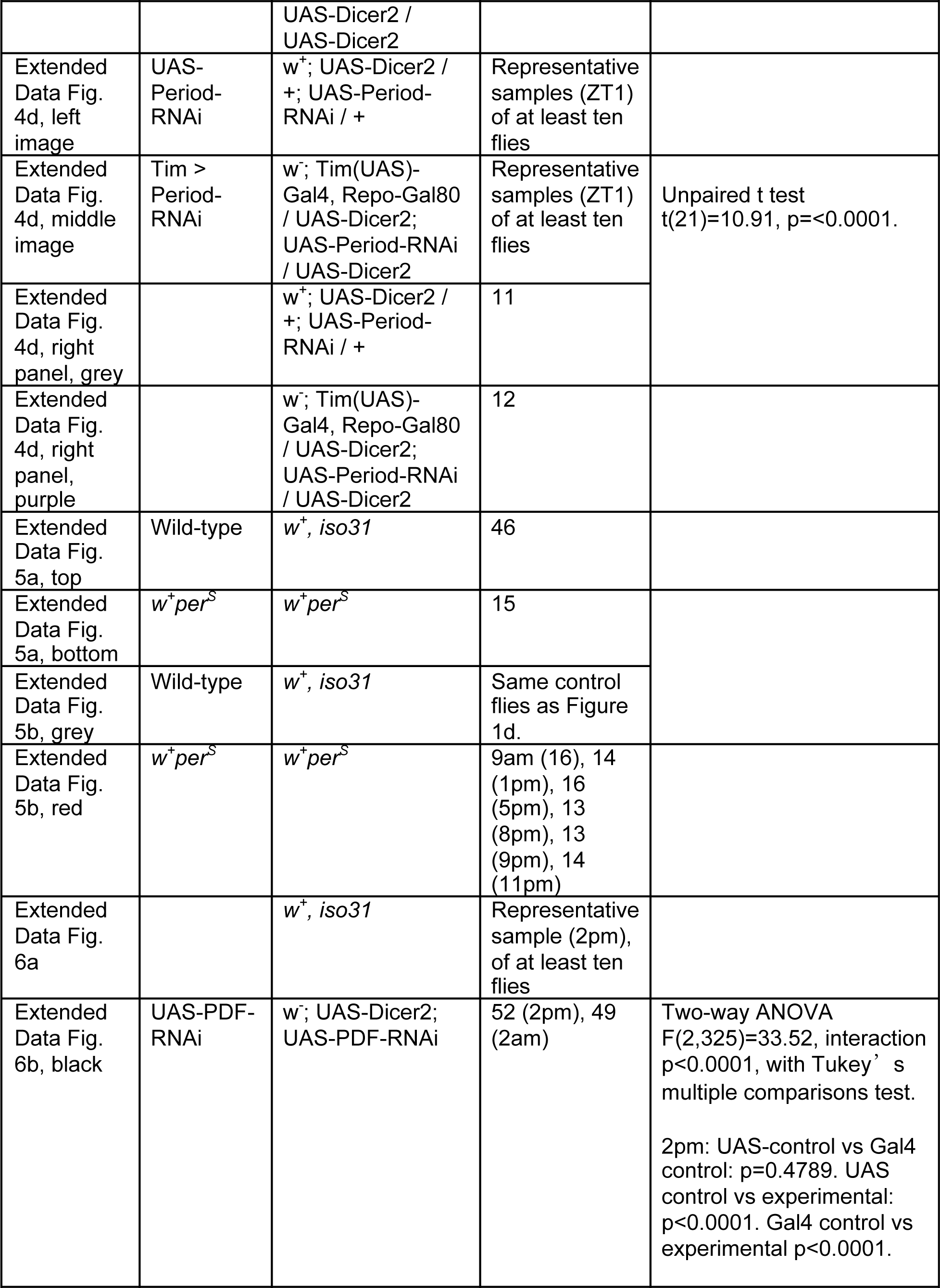
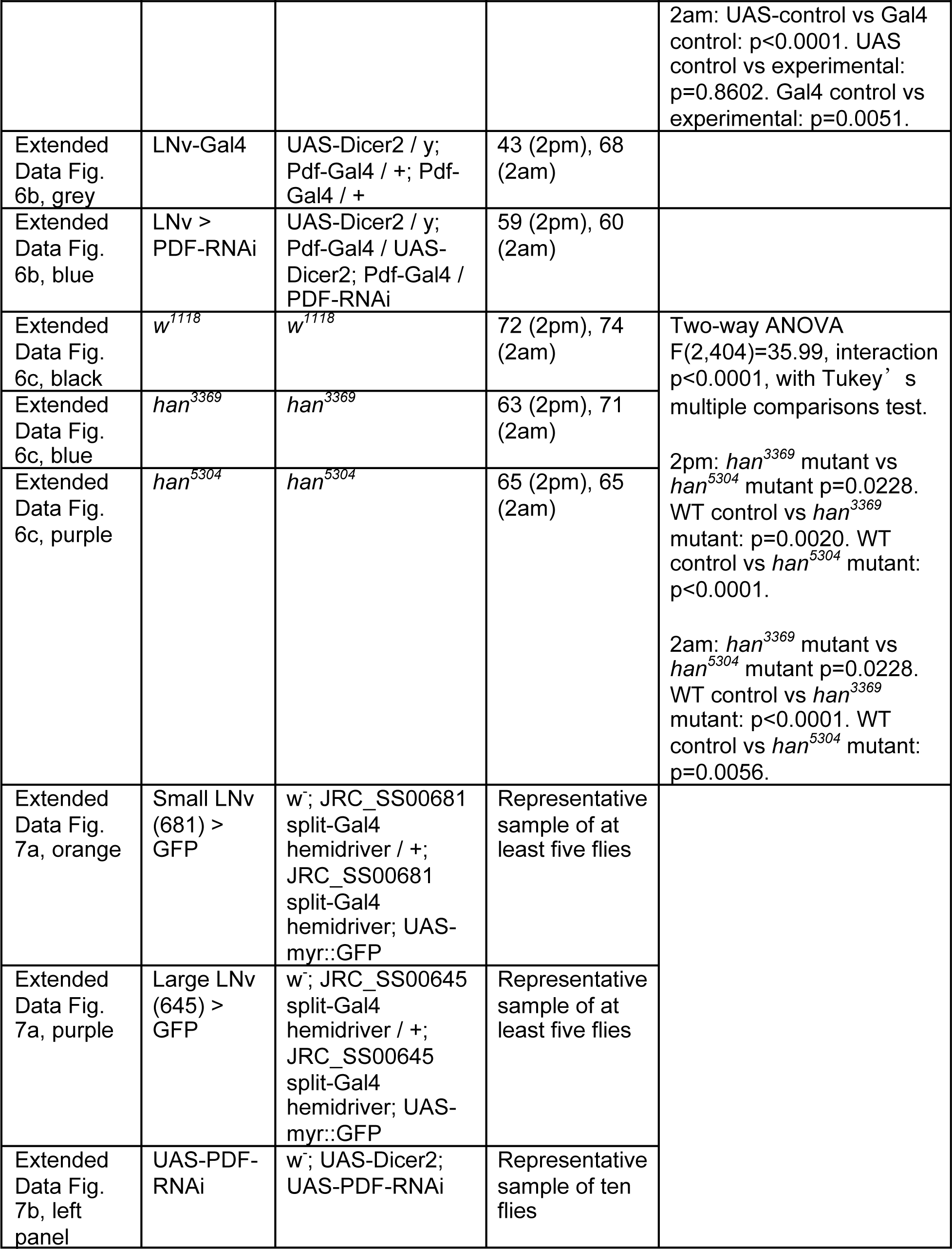
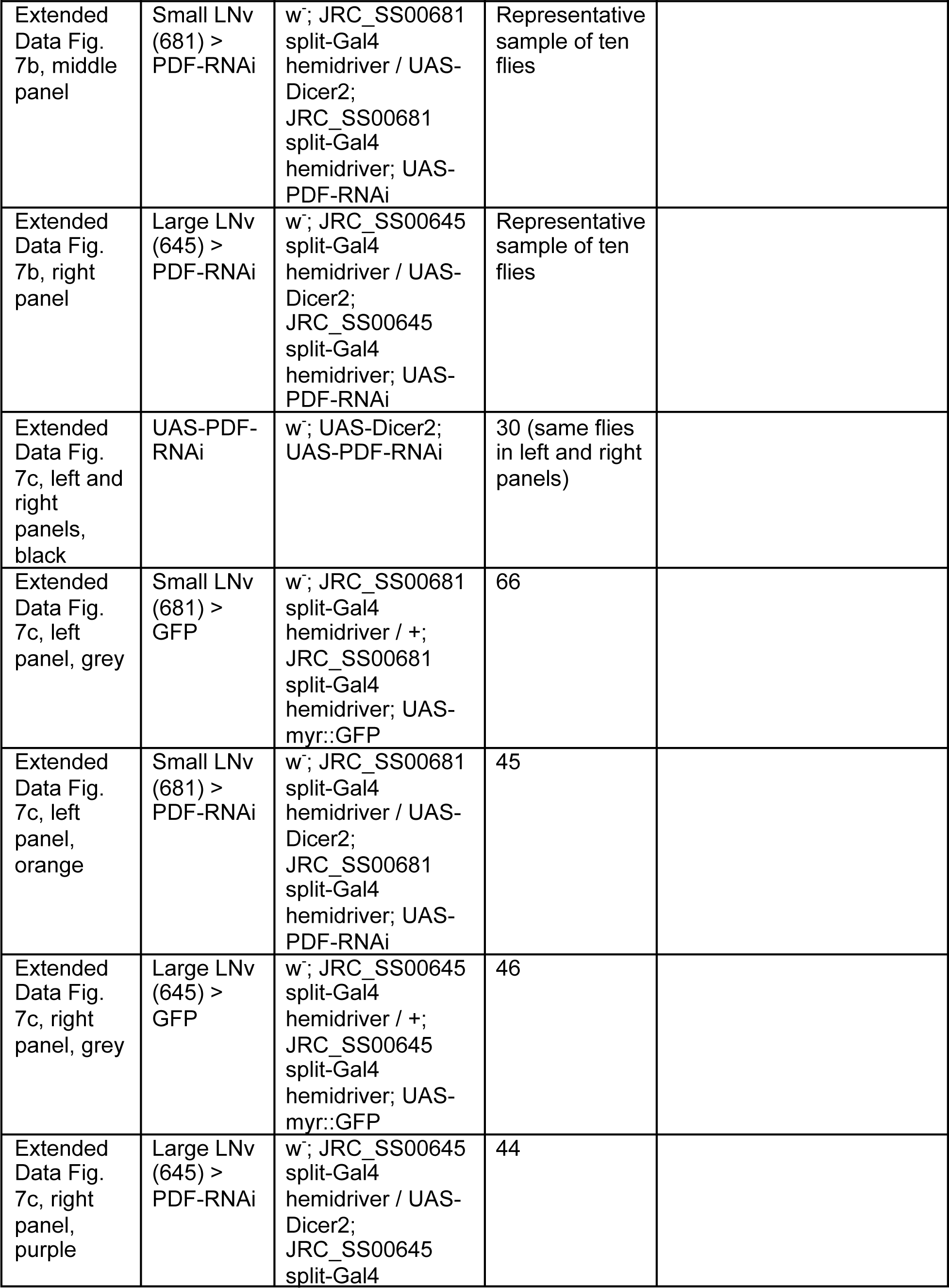
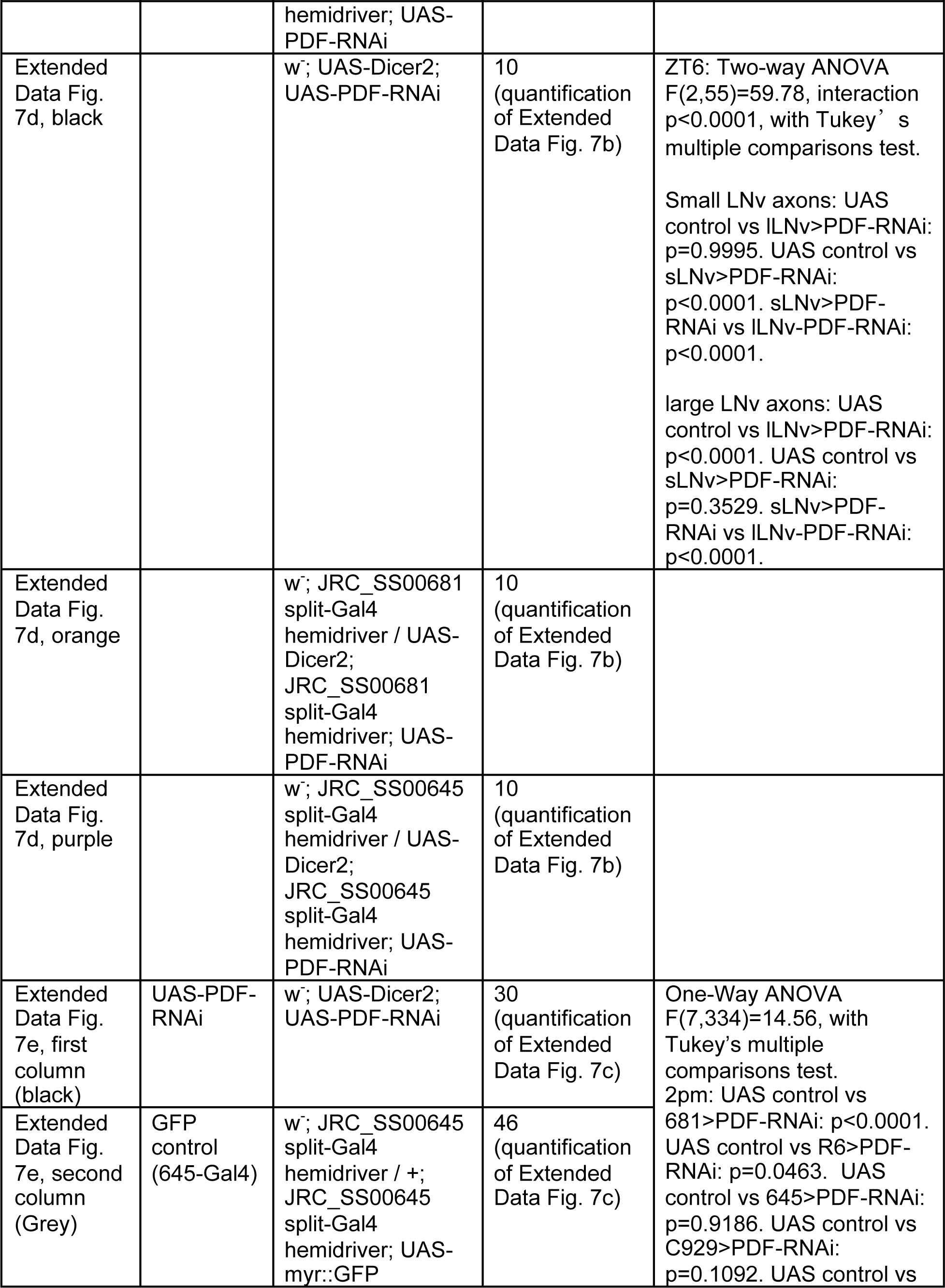
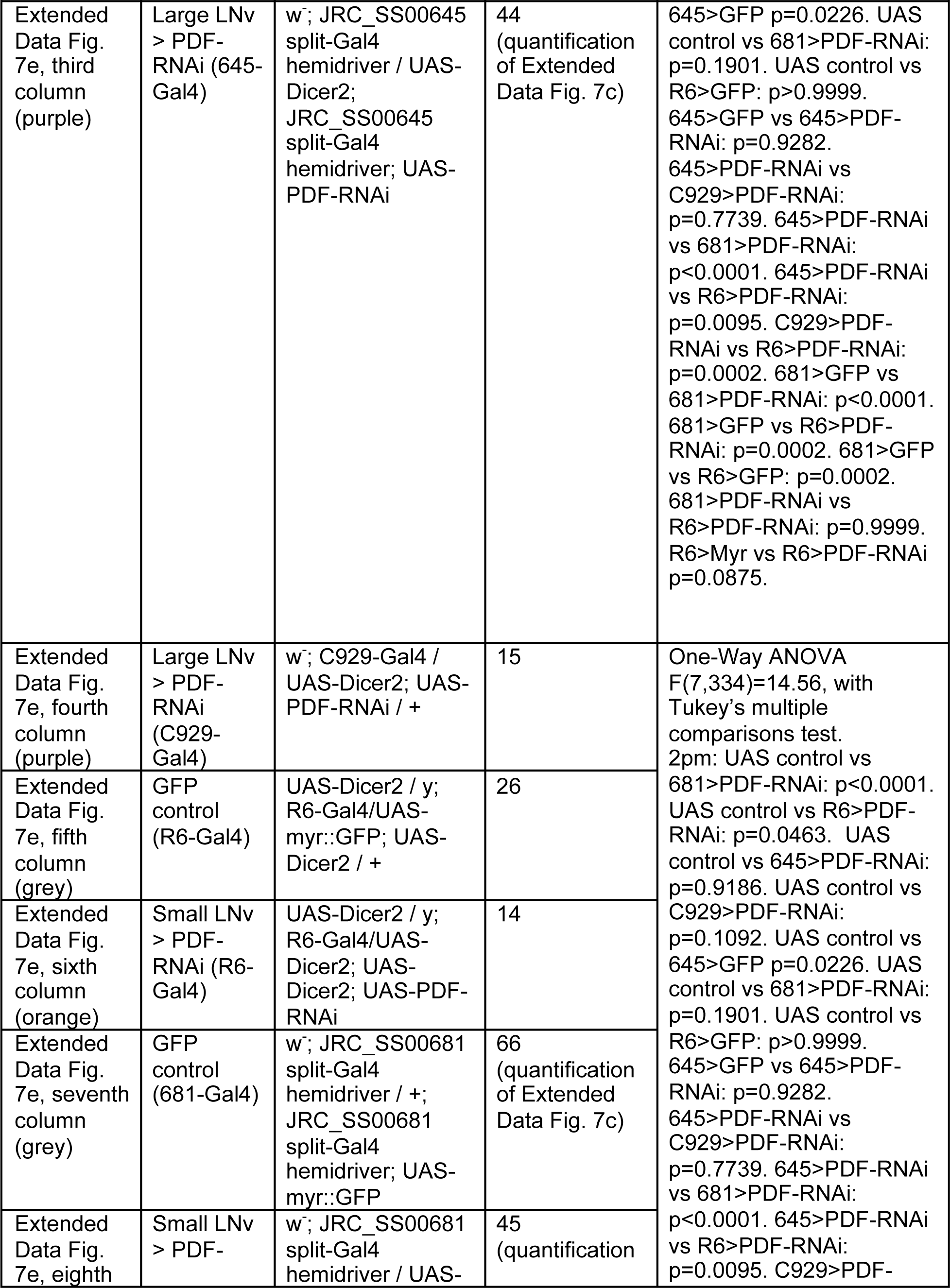
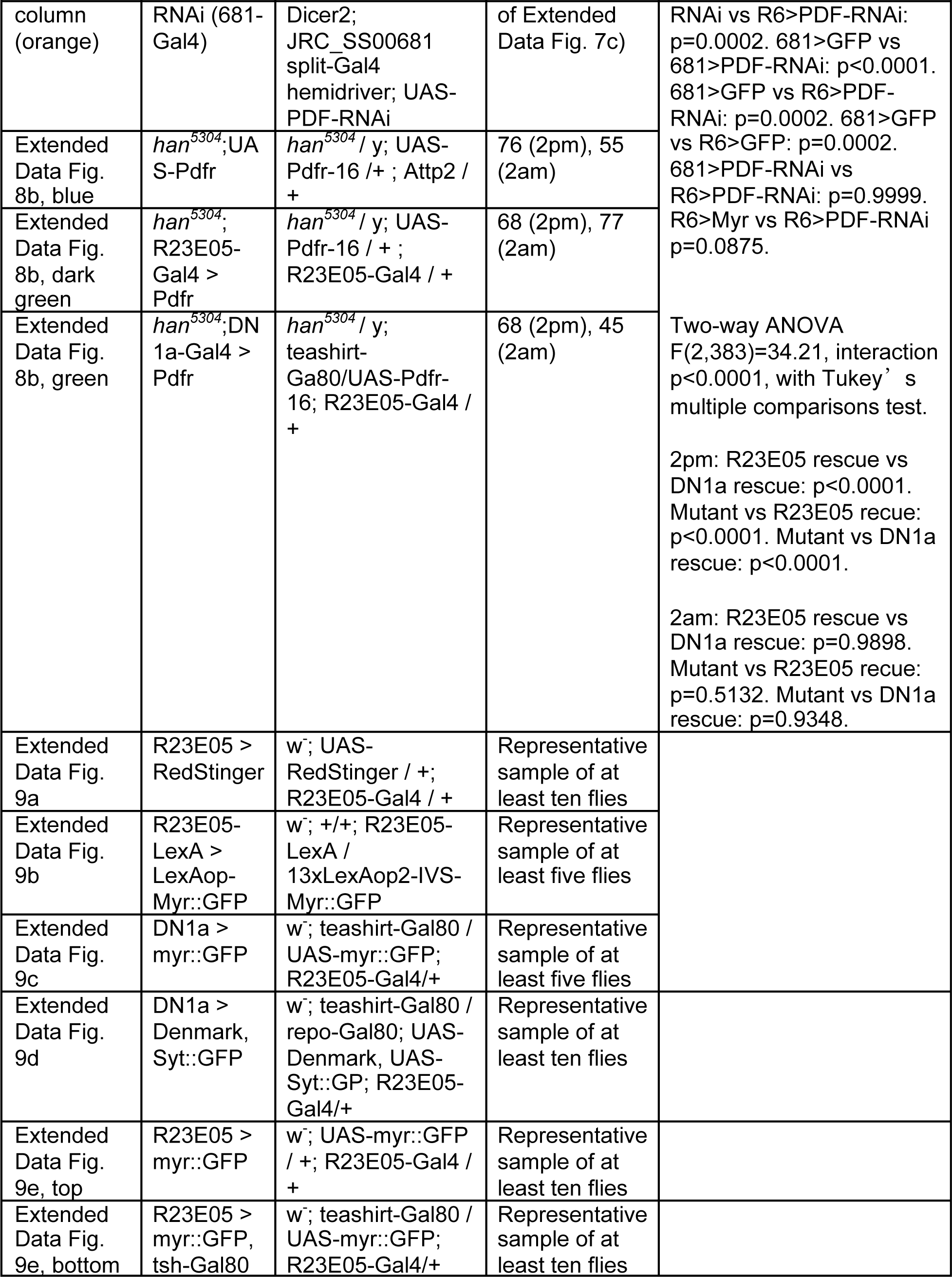
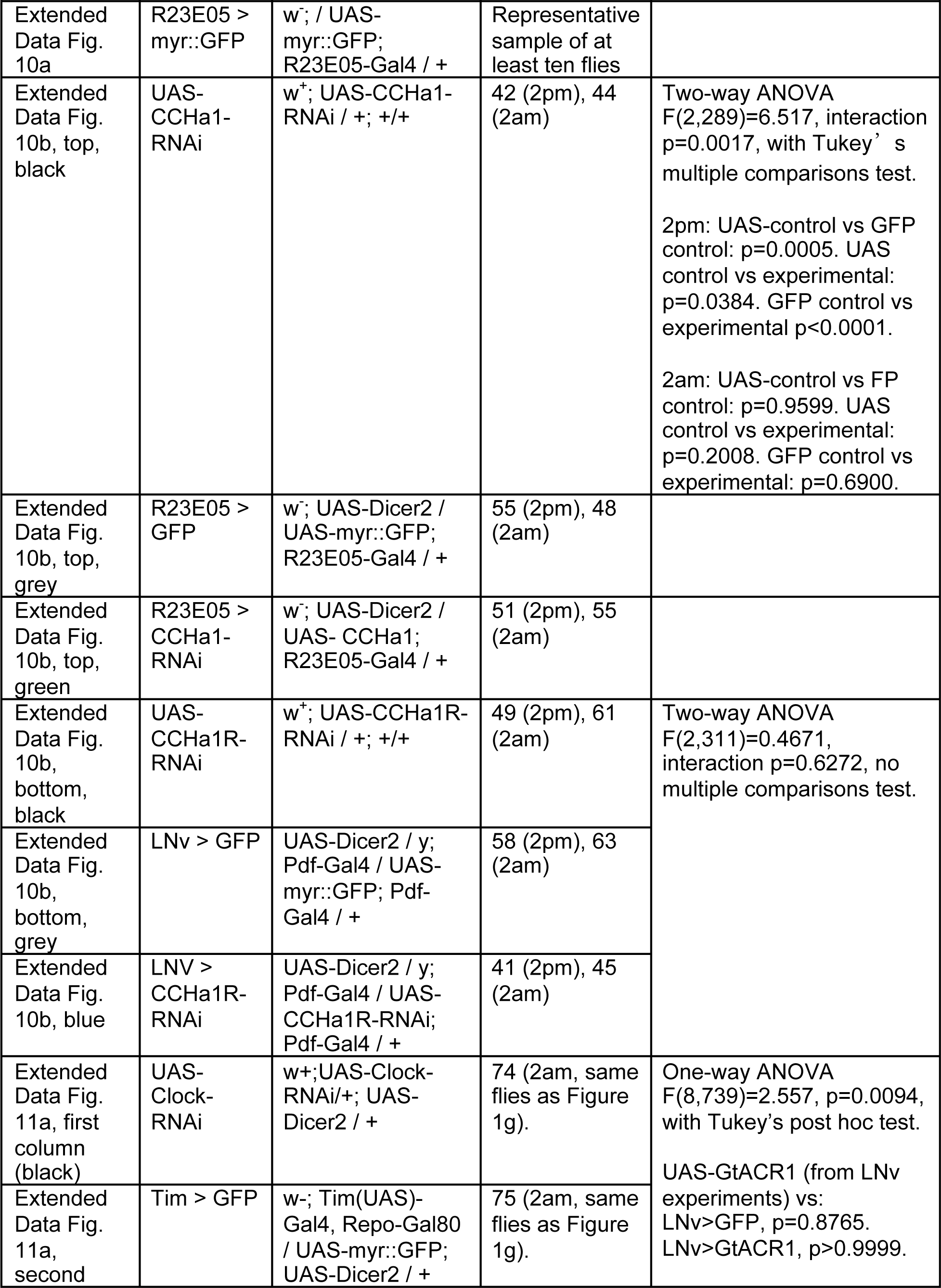
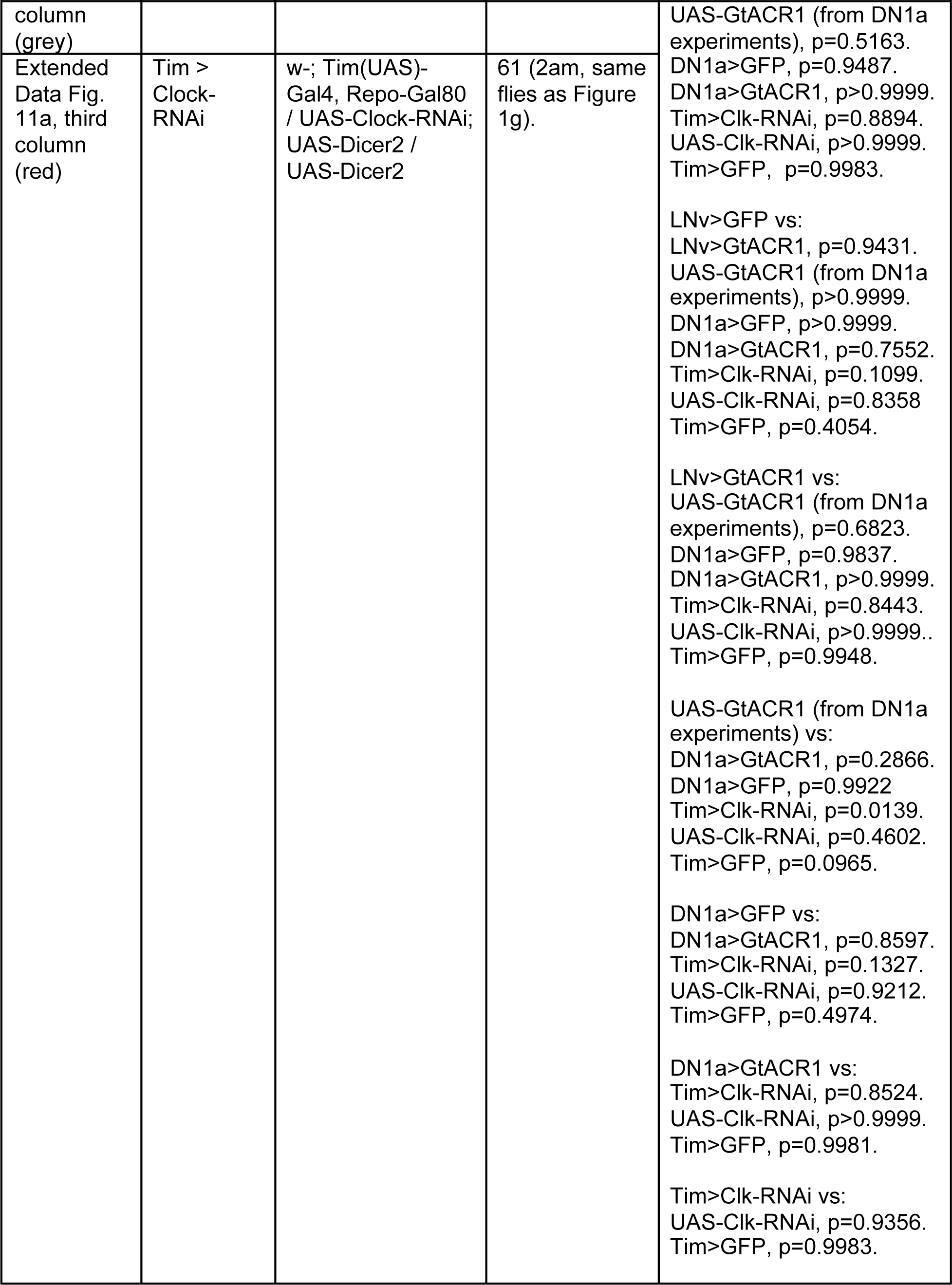
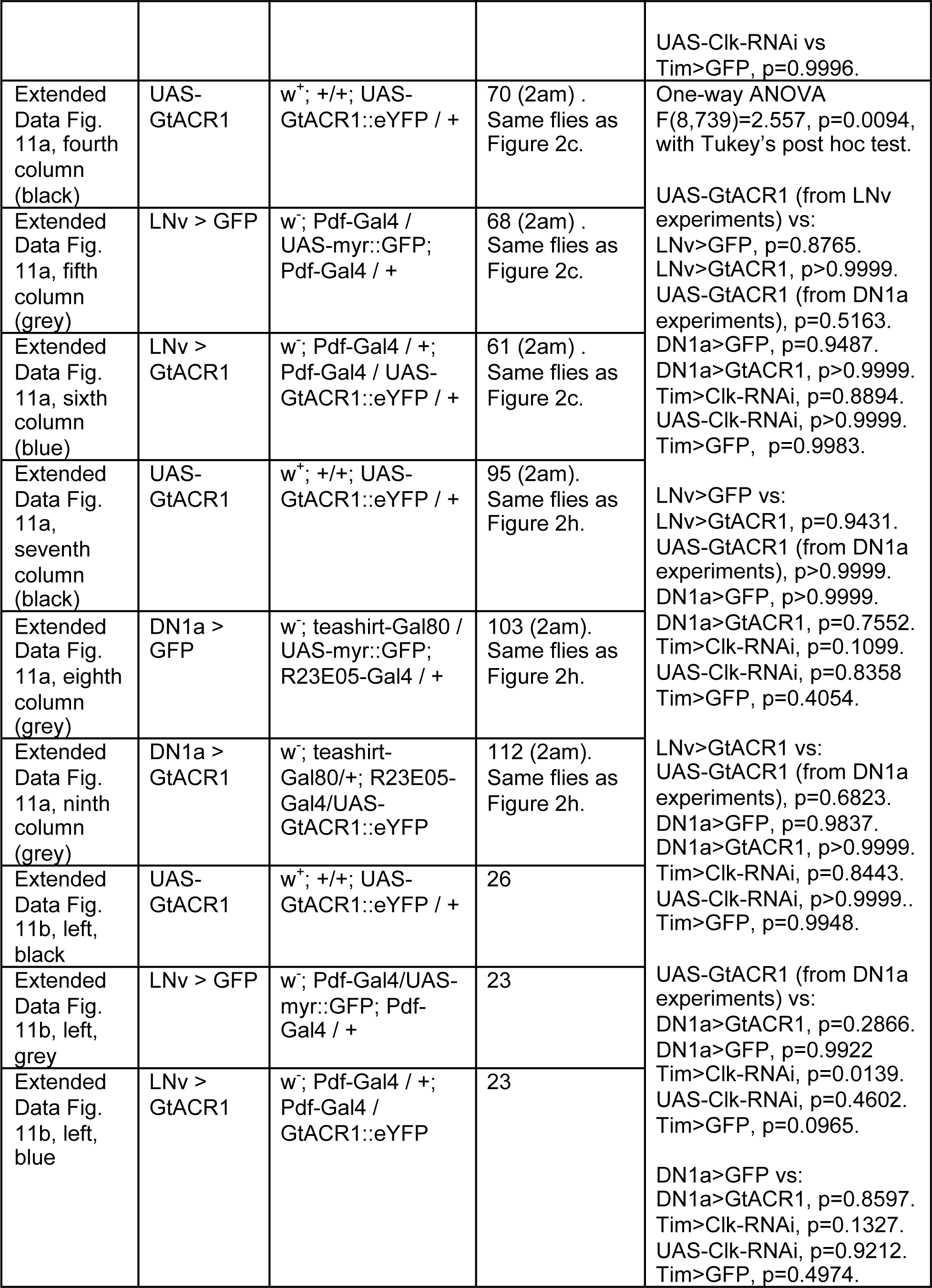
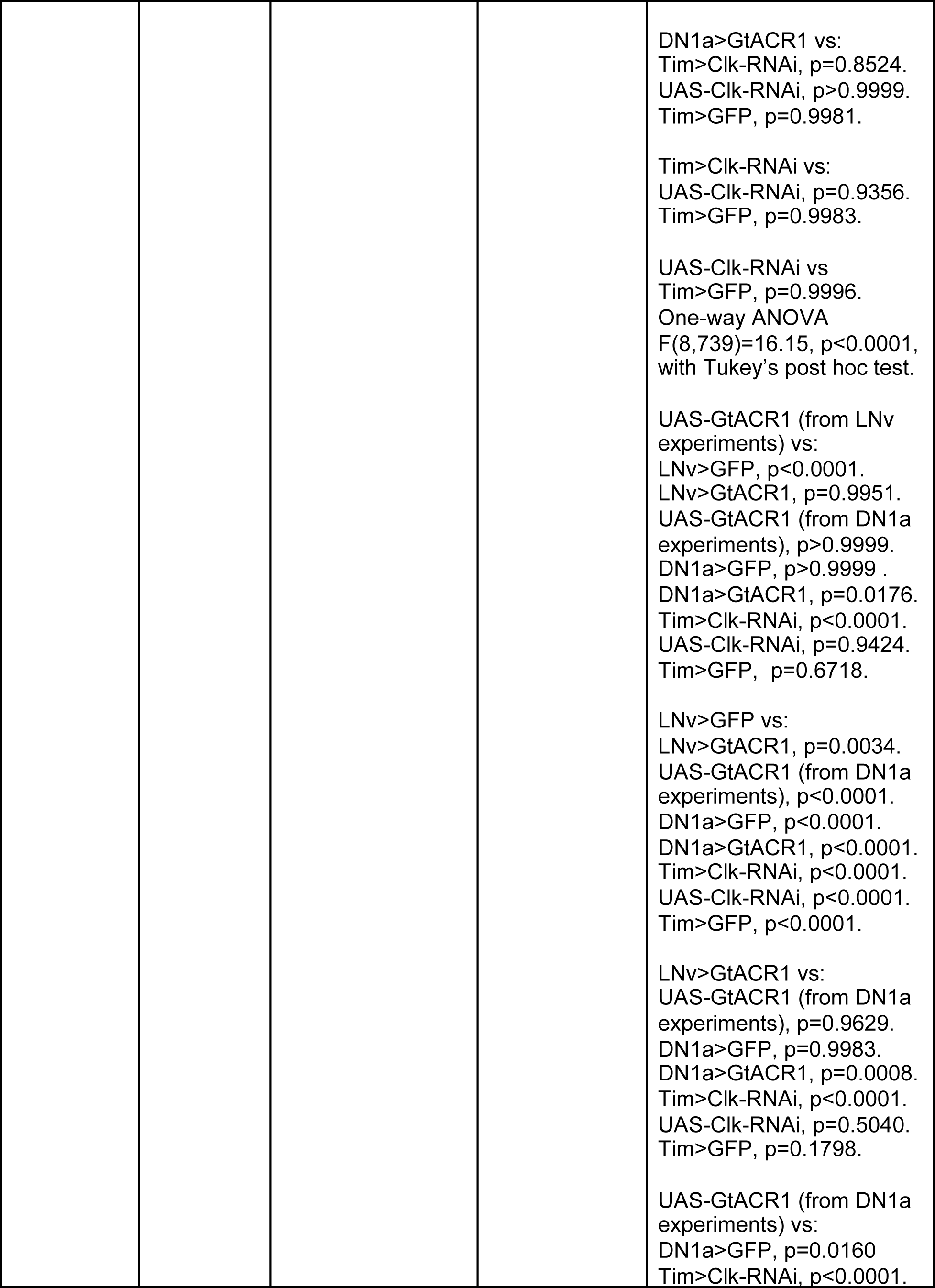
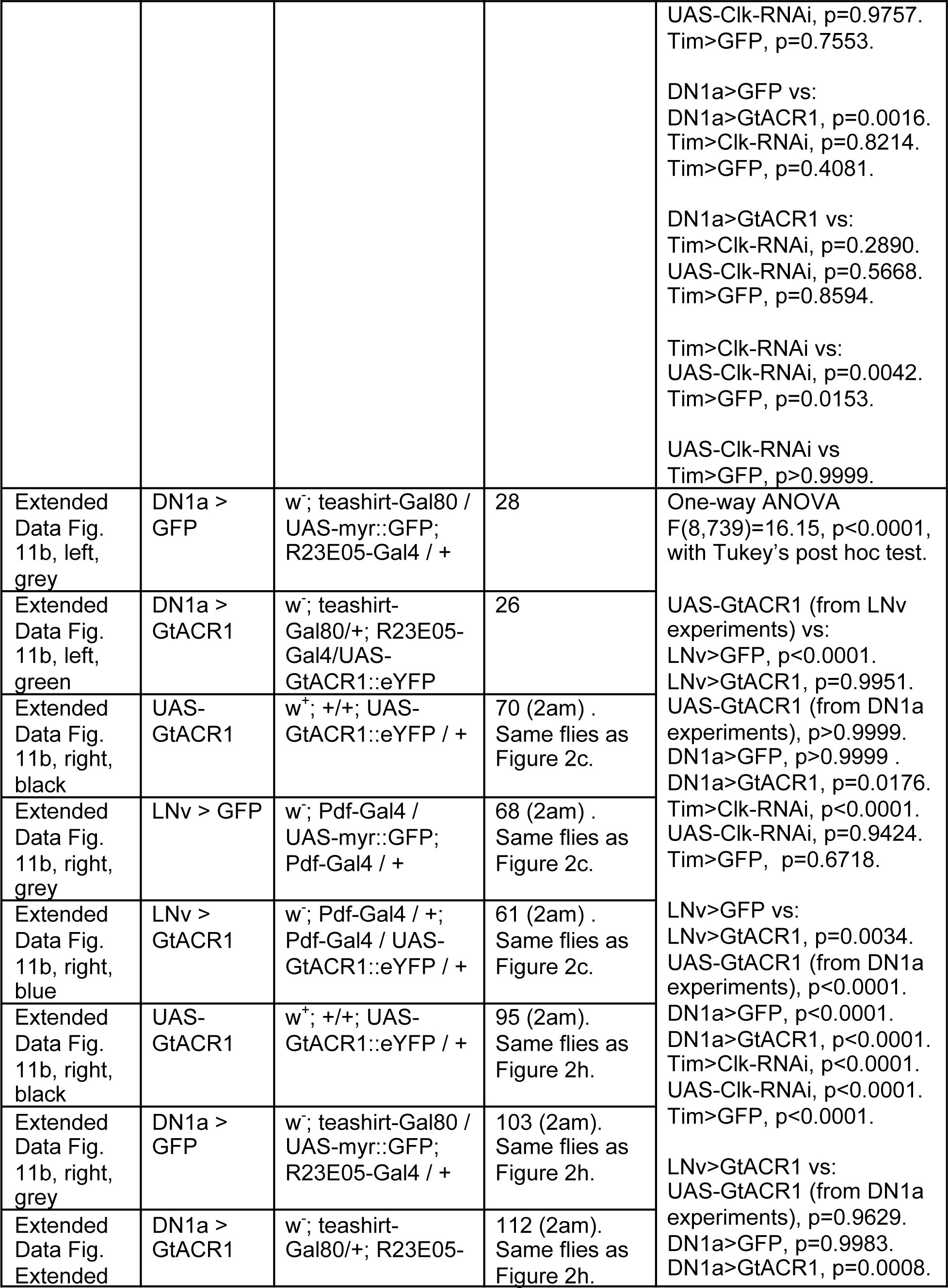
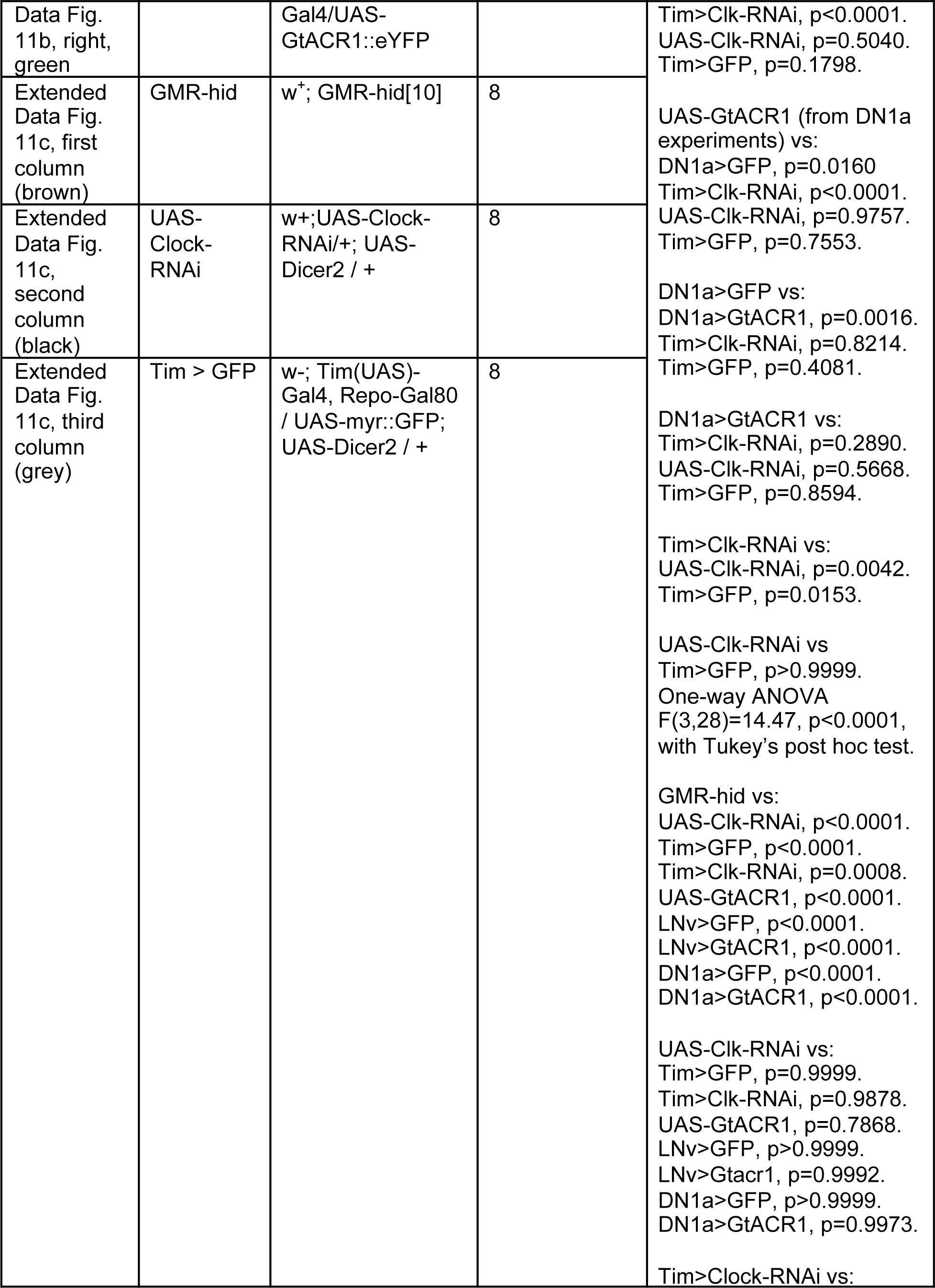
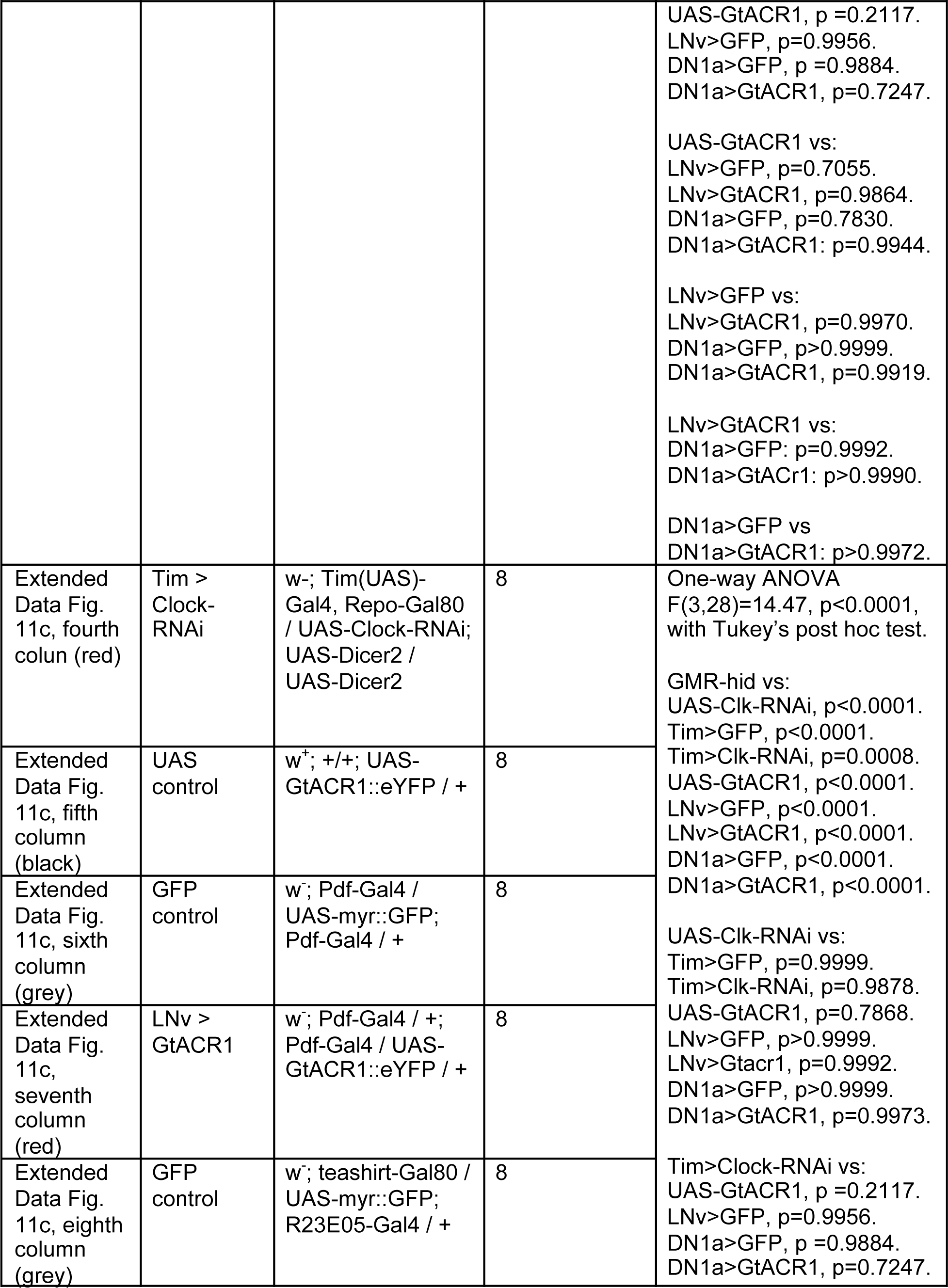
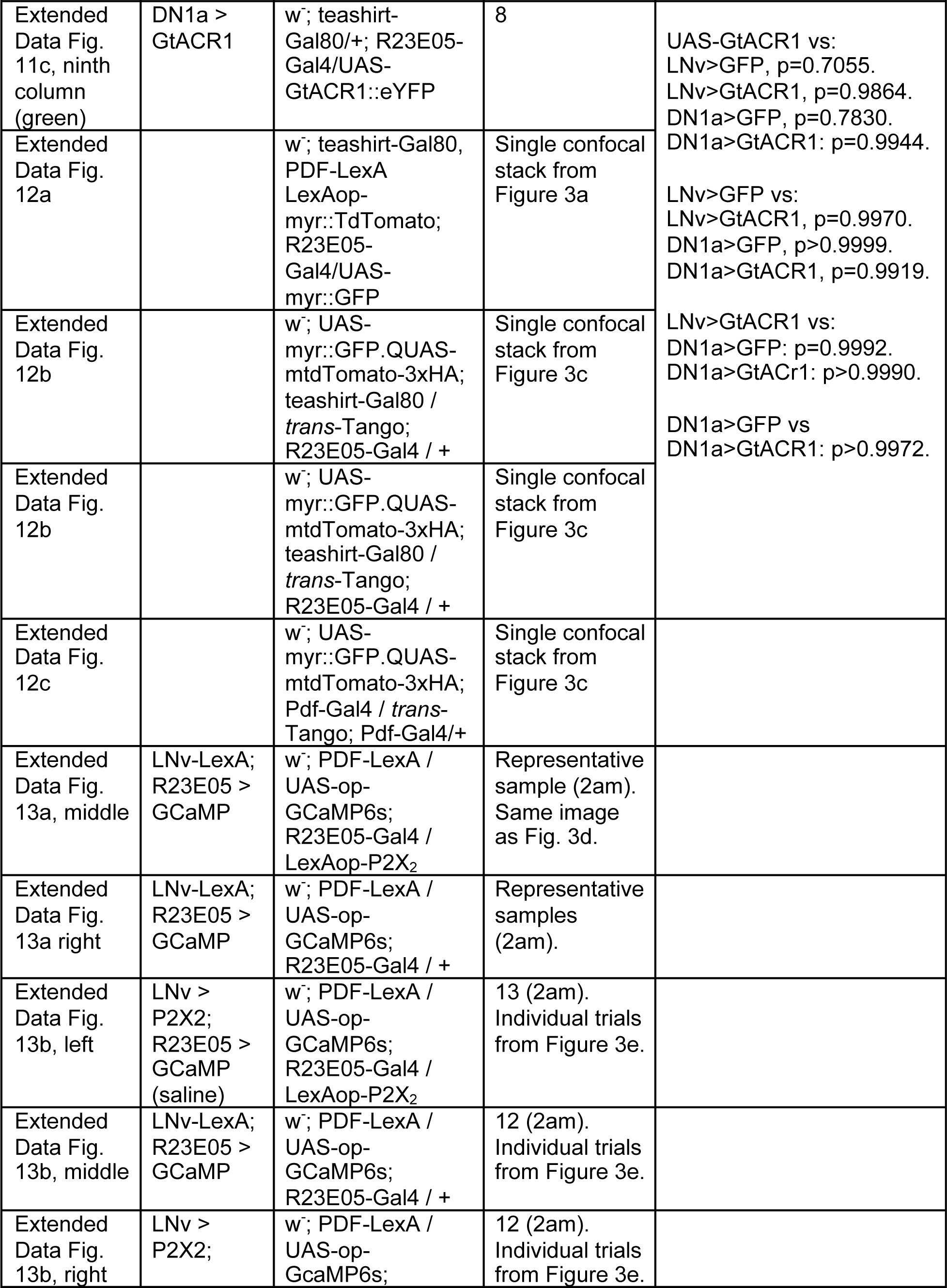
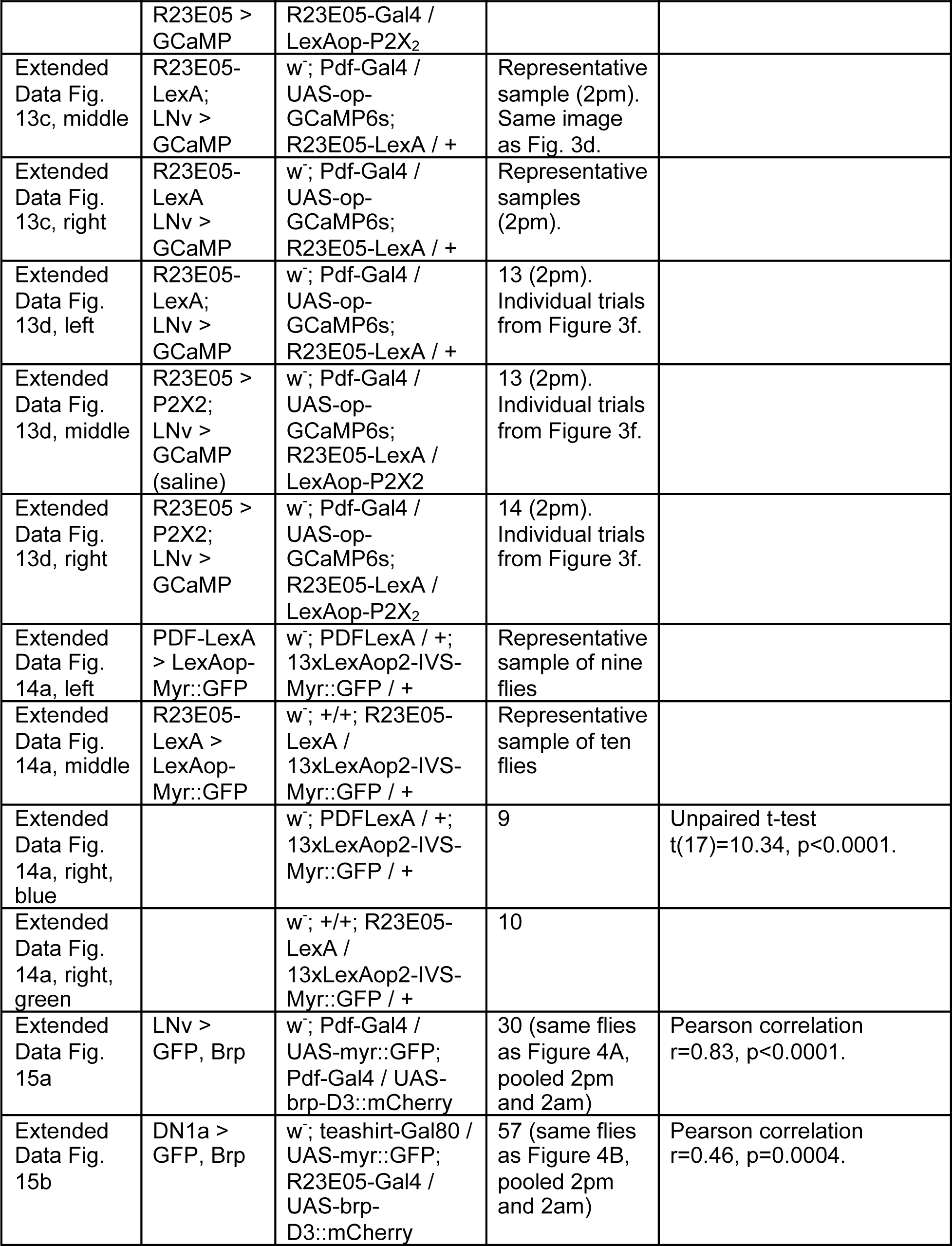
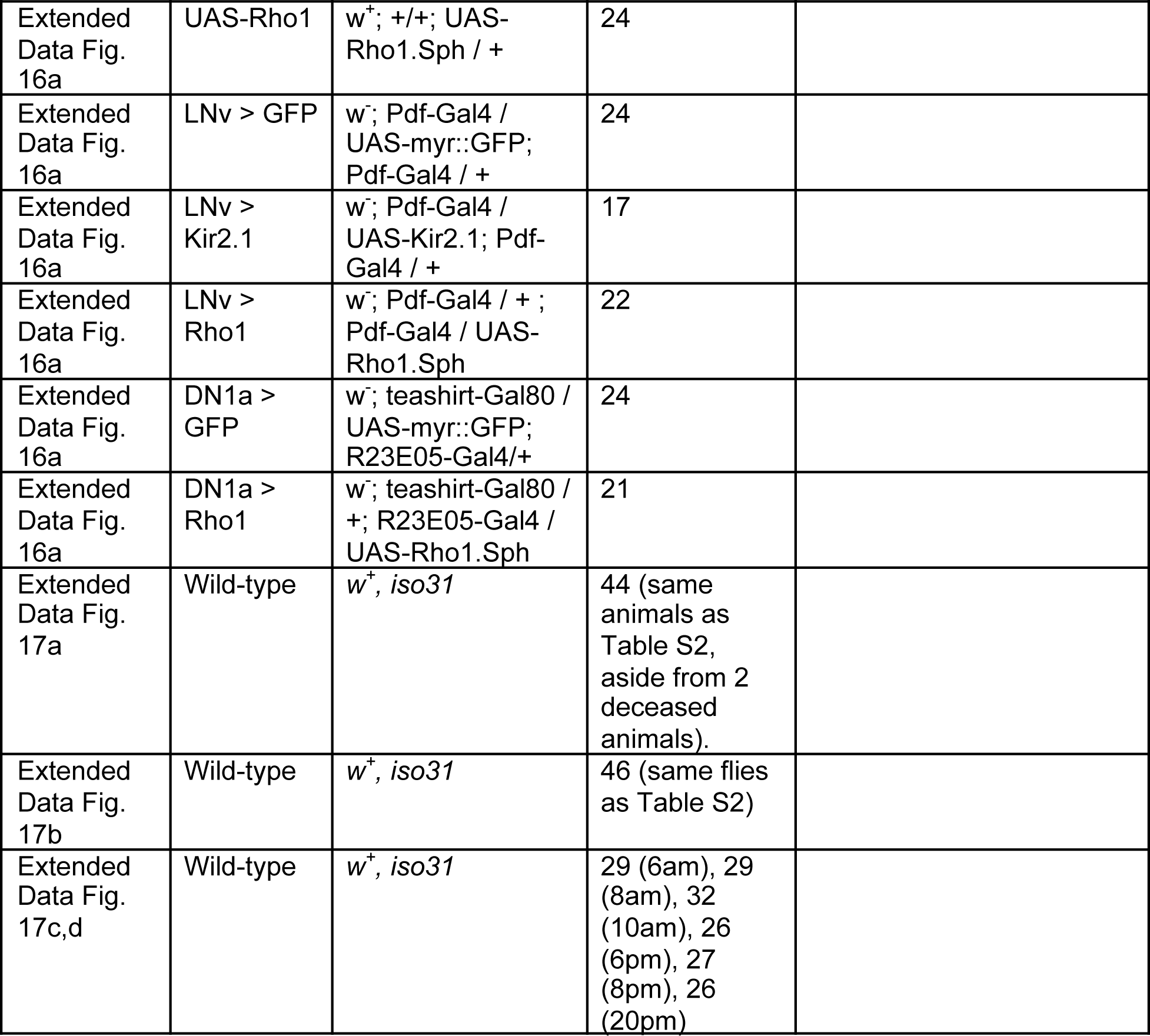

**Extended Data Table 2.** Circadian rhythmicity during the first four days in darkness

| Genotype | N | % Rhythmic after 4 days in darkness | Tau (S.E.M.) | Power (S.E.M.) |
| --- | --- | --- | --- | --- |
| $w^+$, *iso31* | 46 | 100 | 23.9 (0.12) | 71.9 (1.7) |
| UAS-Clock-RNAi | 23 | 100 | 23.8 (0.12) | 74 (2.44) |
| Tim > GFP | 24 | 100 | 23.6 (0.1) | 86.3 (2.33) |
| Tim > Clock-RNAi | 22 | 9.1 | 17.8 (0.75) | 22.7 (2.55) |
| UAS-Period-RNAi | 23 | 100 | 24.7 (0.13) | 71.9 (1.7) |
| Tim > Period-RNAi | 24 | 0 | **** | **** |
| $w^+$ *per<sup>S</sup>* | 19 | 94.7 | 18.9 (0.05) | 69.8 (4.82) |
| UAS-GtACR1 | 20 | 95 | 23.6 (0.06) | 68.2 (4.91) |
| LNv > GFP | 24 | 100 | 24.3 (0.16) | 71 (2.47) |
| LNv > GtACR1 | 21 | 52.4 | 23.8 (0.31) | 53.8 (5.3) |
| UAS-Kir2.1 | 20 | 94.4 | 23.7(0.15) | 74.7 (3.52) |
| LNv > Kir2.1 | 17 | 70.6 | 23 (0.13) | 66.7 (5.71) |
| DN1a > GFP | 25 | 84 | 23.8 (0.11) | 56 (4.59) |
| DN1a > GtACR1 | 19 | 78.9 | 23.9 (0.23) | 37.8 (3.49) |
| *han*<sup>3369</sup>, UAS-PDFR16 | 14 | 57.1 | 23.4 (0.48) | 52.9 (7.09) |
| *han*<sup>5304</sup>, UAS-PDFR16 | 17 | 70.6 | 23.1 (0.21) | 46.2 (5.56) |
| *han*<sup>3369</sup>, DN1a > PDFR16 | 10 | 90 | 23.4 (0.15) | 66 (7.86) |
| *han*<sup>5304</sup>, DN1a > PDFR16 | 17 | 100 | 22.9 (0.12) | 84.1 (3.82) |
| UAS-tethered PDF | 22 | 95.5 | 23.5 (0) | 87.5 (3.79) |
| DN1a > tethered PDF | 21 | 100 | 23.5 (0.02) | 87.8 (3) |
| UAS-Rho1 | 24 | 95.8 | 23.7 (0.09) | 72.2 (3.26) |
| LNv > Rho1 | 22 | 90.9 | 24.8 (0.28) | 52 (2.74) |
| DN1a > Rho1 | 21 | 95.2 | 23.5 (0.07) | 71.5.4 (3.95) |
| UAS-Rho1-RNAi | 24 | 95.8 | 23.9 (0.13) | 73.8 (3.56) |
| LNv > GFP (with Dicer2) | 22 | 100 | 24 (0.16) | 70.1 (2.43) |
| LNv > Rho1-RNAi (with Dicer2) | 15 | 66.7 | 23.6 (0.16) | 55 (6.19) |
| R23E05 > GFP (with Dicer2) | 24 | 100 | 23.5 (0) | 81.8 (4.79) |
| R23E05 > Rho1-RNAi (with Dicer2) | 25 | 72 | 23.9 (0.17) | 60.2 (6.29) |

**Extended Data Table 3.**
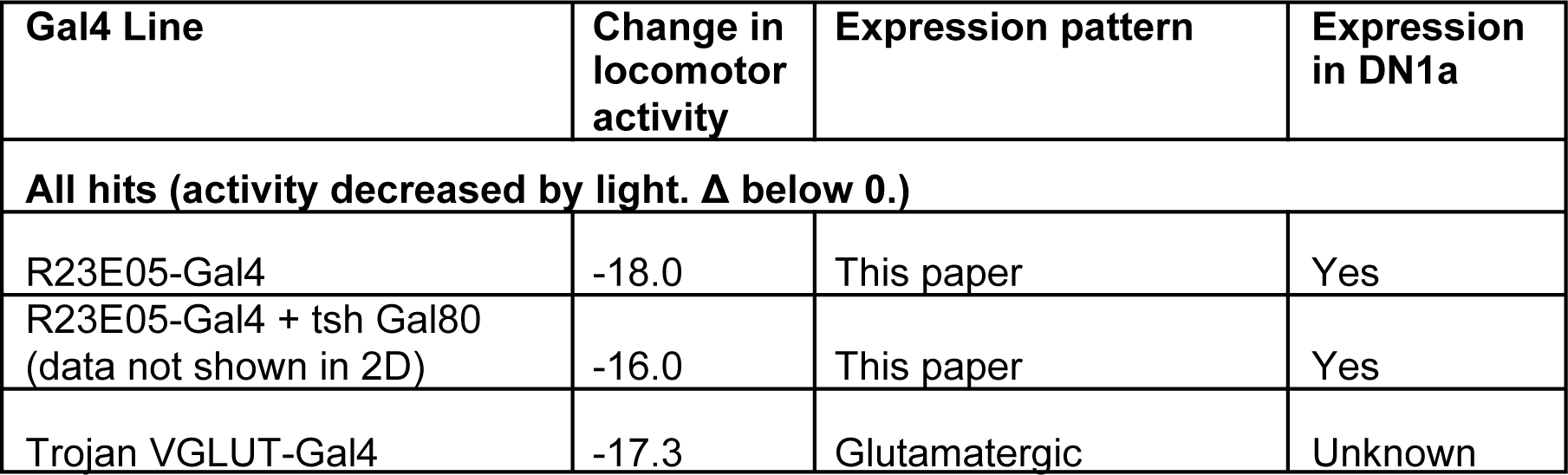
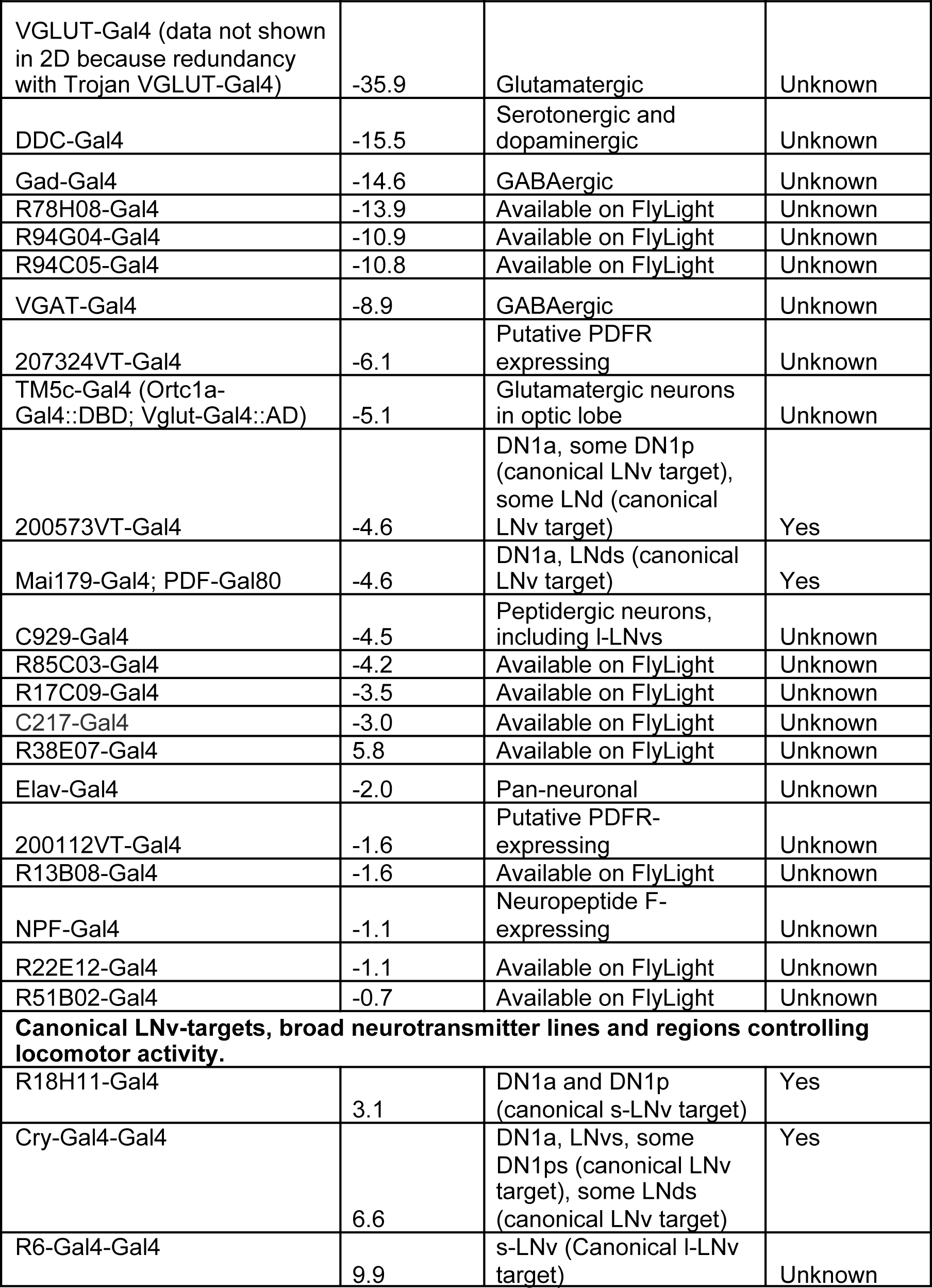
Selected results from PDFR screen

| Gal4 Line | Change in locomotor activity | Expression pattern | Expression in DN1a |
| --- | --- | --- | --- |
| **All hits (activity decreased by light. $\Delta$ below 0.)** |  |  |  |
| R23E05-Gal4 | -18.0 | This paper | Yes |
| R23E05-Gal4 + tsh Gal80 (data not shown in 2D) | -16.0 | This paper | Yes |
| Trojan VGLUT-Gal4 | -17.3 | Glutamatergic | Unknown |
| VGLUT-Gal4 (data not shown in 2D because redundancy with Trojan VGLUT-Gal4) | -35.9 | Glutamatergic | Unknown |
| DDC-Gal4 | -15.5 | Serotonergic and dopaminergic | Unknown |
| Gad-Gal4 | -14.6 | GABAergic | Unknown |
| R78H08-Gal4 | -13.9 | Available on FlyLight | Unknown |
| R94G04-Gal4 | -10.9 | Available on FlyLight | Unknown |
| R94C05-Gal4 | -10.8 | Available on FlyLight | Unknown |
| VGAT-Gal4 | -8.9 | GABAergic | Unknown |
| 207324VT-Gal4 | -6.1 | Putative PDFR expressing | Unknown |
| TM5c-Gal4 (Ortc1a-Gal4::DBD; Vglut-Gal4::AD) | -5.1 | Glutamatergic neurons in optic lobe | Unknown |
| 200573VT-Gal4 | -4.6 | DN1a, some DN1p (canonical LNV target), some LND (canonical LNV target) | Yes |
| Mai179-Gal4; PDF-Gal80 | -4.6 | DN1a, LNDs (canonical LNV target) | Yes |
| C929-Gal4 | -4.5 | Peptidergic neurons, including I-LNVs | Unknown |
| R85C03-Gal4 | -4.2 | Available on FlyLight | Unknown |
| R17C09-Gal4 | -3.5 | Available on FlyLight | Unknown |
| C217-Gal4 | -3.0 | Available on FlyLight | Unknown |
| R38E07-Gal4 | 5.8 | Available on FlyLight | Unknown |
| Elav-Gal4 | -2.0 | Pan-neuronal | Unknown |
| 200112VT-Gal4 | -1.6 | Putative PDFR-expressing | Unknown |
| R13B08-Gal4 | -1.6 | Available on FlyLight | Unknown |
| NPF-Gal4 | -1.1 | Neuropeptide F-expressing | Unknown |
| R22E12-Gal4 | -1.1 | Available on FlyLight | Unknown |
| R51B02-Gal4 | -0.7 | Available on FlyLight | Unknown |
| **Canonical LNV-targets, broad neurotransmitter lines and regions controlling locomotor activity.** |  |  |  |
| R18H11-Gal4 | 3.1 | DN1a and DN1p (canonical s-LNV target) | Yes |
| Cry-Gal4-Gal4 | 6.6 | DN1a, LNVs, some DN1ps (canonical LNV target), some LNDs (canonical LNV target) | Yes |
| R6-Gal4-Gal4 | 9.9 | s-LNV (Canonical I-LNV target) | Unknown |

**Extended Data Table 4.** Sources of fly stocks used in this paper

| *Drosophila melanogaster* | Source | Stock Number |
| --- | --- | --- |
| *iso31* | Ryder et al., Genetics, 2004 |  |
| *Canton S* | Barry Dickson (via Michael Crickmore) |  |
| UAS-myr::GFP (attP2) | BDSC (Bloomington Drosophila Stock Center) 32197 (via Matt Pecot) |  |
| UAS-myr::GFP (attP40) | BDSC 32198 |  |
| Tim(UAS)-Gal4 | Blau and Young, Cell, 1999 (Flybase:FBtp0011839) |  |
| UAS-Dicer2 (X) | BDSC 24646 |  |
| UAS-Dicer2 (2) | BDSC 24650 |  |
| UAS-Dicer2 (3) | BDSC 24651 |  |
| repo-Gal80 | Awasaki et al., J. Neurosci, 2008 (Flybase:FBtp0067904) |  |
| UAS-Clock-RNAi | Vienna Drosophila Resource Center VDRC (VDRC) | 107576KK |
| UAS-Period-RNAi | Fly Stocks of National Institutes of Genetics (via Michael Young) | 2647R-1 |
| *w<sup>+</sup>per<sup>S</sup>* | Jeffrey Price |  |
| yw; PDF-Gal4; PDF-Gal4 (LNv-Gal4) | Justin Blau |  |
| UAS-GtACR1::eYFP | Adam Claridge-Chang (via Michael Crickmore) |  |
| UAS-ChR2-XXM | Robert Kittel (via Michael Crickmore) |  |
| R23E05-Gal4 | BDSC | BDSC_49029 |
| UAS-ChR2-XXL | BDSC | BDSC_58374 |
| *trans*-Tango | BDSC | BDSC_77124 |
| *pdf<sup>Han3369</sup>* | Laboratory of Paul Taghert |  |
| *pdf<sup>Han5304</sup>* | Laboratory of Paul Taghert |  |
| UAS-Pdfr-16 | Laboratory of Paul Taghert |  |
| UAS-Denmark | BDSC | BDSC_33062 |
| PDF-LexA | Shang et al., PNAS, 2008 (Flybase: FBtp0093323) |  |
| LexAop-myr::tdTom | Laboratory of Matt Pecot (Chen et al., 2014) |  |
| LexAop-syt::GDP::HA | BDSC | BDSC_62142 |
| UAS-opGCaMP6s | Laboratory of David Anderson (via Michael Crickmore) |  |
| UAS-tdTomato | BDSC 36327 |  |
| LexAop-P2X2 | Laboratory of Orie Shafer (via Rachel Wilson) |  |
| UAS-brp-D3::mCherry | Christiansen et al., J. Neurosci, 2011 (Flybase: FBtp0069949) |  |
| *per*<sup>01</sup> | Konopka and Benzer, PNAS, 1971 (Flybase: FBal0013649) |  |
| UAS-Rho1.Sph | BDSC | BDSC_7334 |
| UAS-Rho1-RNAi | BDSC | BDSC_9909 |
| R6-Gal4 | Helfrich-Forster et al., J. Comp. Neurol., 2007 (Flybase FBti0016844) |  |
| JRC_SS00645 | Laboratory of Gerry Rubin |  |
| C929-Gal4 | BDSC | BDSC_9909 |
| JRC_SS00681 | Laboratory of Gerry Rubin |  |
| UAS- PDF-RNAi | VDRC | 4380 |
| UAS-10x UAS t-PDF | Laboratory of Michael Nitabach |  |
| R23E05-Gal4 | BDSC | BDSC_49029 |
| R23E05-LexA | This paper |  |
| R23E05-Gal80 | This paper |  |
| UAS-RedStinger | BDSC | BDSC_8546 |
| PDF-Gal80 | Stoleru et al., Nature, 2004 (Flybase: Fbtp0019042) |  |
| LexAop-myr::GFP | BDSC (via Matt Pecot) | BDSC_32209 |
| Vglut-Gal4 (OK371) | BDSC | BDSC_26160 |
| GMR-hid[10] | Laboratory of<br>Andreas Bergmann |  |
| UAS-mGluRA-RNAi | BDSC | BDSC_34872 |
| UAS-CCHa1-RNAi | VDRC | 104794KK |
| UAS-CCHa1R-RNAi | VDRC | 103055KK |
| UAS-mCD8::GFP | Laboratory of<br>Michael Crickmore |  |
| UAS-Kir2.1 | Laboratory of<br>Michael Crickmore |  |

## References

1. Watabe-Uchida, M., Eshel, N. & Uchida, N. Neural Circuitry of Reward Prediction Error. Annu Rev Neurosci 40, 373–394, doi:10.1146/annurev-neuro-072116-031109 (2017).

2. Mirenowicz, J. & Schultz, W. Importance of unpredictability for reward responses in primate dopamine neurons. J Neurophysiol 72, 1024–1027, doi:10.1152/jn.1994.72.2.1024 (1994).

3. Schultz, W., Dayan, P. & Montague, P. R. A neural substrate of prediction and reward. Science 275, 1593–1599, doi:10.1126/science.275.5306.1593 (1997).

4. Vitaterna, M. H., Takahashi, J. S. & Turek, F. W. Overview of circadian rhythms. Alcohol Res Health 25, 85–93 (2001).

5. Golombek, D. A. & Rosenstein, R. E. Physiology of circadian entrainment. Physiol Rev 90, 1063–1102, doi:10.1152/physrev.00009.2009 (2010).

6. Rensing, L. & Ruoff, P. Temperature effect on entrainment, phase shifting, and amplitude of circadian clocks and its molecular bases. Chronobiol Int 19, 807–864, doi:10.1081/cbi-120014569 (2002).

7. Top, D. & Young, M. W. Coordination between Differentially Regulated Circadian Clocks Generates Rhythmic Behavior. Cold Spring Harb Perspect Biol 10, doi:10.1101/cshperspect.a033589 (2018).

8. Eckel-Mahan, K. & Sassone-Corsi, P. Metabolism and the circadian clock converge. Physiol Rev 93, 107–135, doi:10.1152/physrev.00016.2012 (2013).

9. Lu, B., Liu, W., Guo, F. & Guo, A. Circadian modulation of light-induced locomotion responses in Drosophila melanogaster. Genes Brain Behav 7, 730–739, doi:10.1111/j.1601-183X.2008.00411.x (2008).

10. Ryder, E. et al. The DrosDel collection: a set of P-element insertions for generating custom chromosomal aberrations in Drosophila melanogaster. Genetics 167, 797–813, doi:10.1534/genetics.104.026658 (2004).

11. Huber, R. et al. Sleep homeostasis in Drosophila melanogaster. Sleep 27, 628–639, doi:10.1093/sleep/27.4.628 (2004).

12. Dubowy, C. & Sehgal, A. Circadian Rhythms and Sleep in Drosophila melanogaster. Genetics 205, 1373–1397, doi:10.1534/genetics.115.185157 (2017).

13. Konopka, R. J. & Benzer, S. Clock mutants of Drosophila melanogaster. Proc Natl Acad Sci U S A 68, 2112–2116, doi:10.1073/pnas.68.9.2112 (1971).

14. Bargiello, T. A., Jackson, F. R. & Young, M. W. Restoration of circadian behavioural rhythms by gene transfer in Drosophila. Nature 312, 752–754, doi:10.1038/312752a0 (1984).

15. Zehring, W. A. et al. P-element transformation with period locus DNA restores rhythmicity to mutant, arrhythmic Drosophila melanogaster. Cell 39, 369–376, doi:10.1016/0092-8674(84)90015-1 (1984).

16. Tataroglu, O. & Emery, P. Studying circadian rhythms in Drosophila melanogaster. Methods 68, 140–150, doi:10.1016/j.ymeth.2014.01.001 (2014).

17. Blau, J. & Young, M. W. Cycling vrille expression is required for a functional Drosophila clock. Cell 99, 661–671, doi:10.1016/s0092-8674(00)81554-8 (1999).

18. Renn, S. C., Park, J. H., Rosbash, M., Hall, J. C. & Taghert, P. H. A pdf neuropeptide gene mutation and ablation of PDF neurons each cause severe abnormalities of behavioral circadian rhythms in Drosophila. Cell 99, 791–802 (1999).

19. Mohammad, F. et al. Optogenetic inhibition of behavior with anion channelrhodopsins. Nat Methods 14, 271–274, doi:10.1038/nmeth.4148 (2017).

20. Helfrich-Forster, C. The period clock gene is expressed in central nervous system neurons which also produce a neuropeptide that reveals the projections of circadian pacemaker cells within the brain of Drosophila melanogaster. Proc Natl Acad Sci U S A 92, 612–616, doi:10.1073/pnas.92.2.612 (1995).

21. Hyun, S. et al. Drosophila GPCR Han is a receptor for the circadian clock neuropeptide PDF. Neuron 48, 267–278, doi:10.1016/j.neuron.2005.08.025 (2005).

22. Schlichting, M., Diaz, M. M., Xin, J. & Rosbash, M. Neuron-specific knockouts indicate the importance of network communication to Drosophila rhythmicity. Elife 8, doi:10.7554/eLife.48301 (2019).

23. Lear, B. C., Zhang, L. & Allada, R. The neuropeptide PDF acts directly on evening pacemaker neurons to regulate multiple features of circadian behavior. PLoS Biol 7, e1000154, doi:10.1371/journal.pbio.1000154 (2009).

24. Zhang, L. et al. DN1(p) circadian neurons coordinate acute light and PDF inputs to produce robust daily behavior in Drosophila. Curr Biol 20, 591–599, doi:10.1016/j.cub.2010.02.056 (2010).

25. Pirez, N., Christmann, B. L. & Griffith, L. C. Daily rhythms in locomotor circuits in Drosophila involve PDF. J Neurophysiol 110, 700–708, doi:10.1152/jn.00126.2013 (2013).

26. Jenett, A. et al. A GAL4-driver line resource for Drosophila neurobiology. Cell Rep 2, 991–1001, doi:10.1016/j.celrep.2012.09.011 (2012).

27. Hamasaka, Y. et al. Glutamate and its metabotropic receptor in Drosophila clock neuron circuits. J Comp Neurol 505, 32–45, doi:10.1002/cne.21471 (2007).

28. Shafer, O. T., Helfrich-Forster, C., Renn, S. C. & Taghert, P. H. Reevaluation of Drosophila melanogaster’s neuronal circadian pacemakers reveals new neuronal classes. J Comp Neurol 498, 180–193, doi:10.1002/cne.21021 (2006).

29. Fujiwara, Y. et al. The CCHamide1 Neuropeptide Expressed in the Anterior Dorsal Neuron 1 Conveys a Circadian Signal to the Ventral Lateral Neurons in Drosophila melanogaster. Front Physiol 9, 1276, doi:10.3389/fphys.2018.01276 (2018).

30. Alpert, M. H. et al. A Circuit Encoding Absolute Cold Temperature in Drosophila. Curr Biol, doi:10.1016/j.cub.2020.04.038 (2020).

31. Ribeiro, I. M. A. et al. Visual Projection Neurons Mediating Directed Courtship in Drosophila. Cell 174, 607–621 e618, doi:10.1016/j.cell.2018.06.020 (2018).

32. Markow, T. A. & Manning, M. Mating success of photoreceptor mutants of Drosophila melanogaster. Behav Neural Biol 29, 276–280, doi:10.1016/s0163-1047(80)90612-3 (1980).

33. Choi, C. et al. Cellular dissection of circadian peptide signals with genetically encoded membrane-tethered ligands. Curr Biol 19, 1167–1175, doi:10.1016/j.cub.2009.06.029 (2009).

34. Choi, C. & Nitabach, M. N. Membrane-tethered ligands: tools for cell-autonomous pharmacological manipulation of biological circuits. Physiology (Bethesda) 28, 164–171, doi:10.1152/physiol.00056.2012 (2013).

35. Im, S. H., Li, W. & Taghert, P. H. PDFR and CRY signaling converge in a subset of clock neurons to modulate the amplitude and phase of circadian behavior in Drosophila. PLoS One 6, e18974, doi:10.1371/journal.pone.0018974 (2011).

36. Shafer, O. T. et al. Widespread receptivity to neuropeptide PDF throughout the neuronal circadian clock network of Drosophila revealed by real-time cyclic AMP imaging. Neuron 58, 223–237, doi:10.1016/j.neuron.2008.02.018 (2008).

37. Nicolai, L. J. et al. Genetically encoded dendritic marker sheds light on neuronal connectivity in Drosophila. Proc Natl Acad Sci U S A 107, 20553–20558, doi:10.1073/pnas.1010198107 (2010).

38. Zhang, Y. Q., Rodesch, C. K. & Broadie, K. Living synaptic vesicle marker: synaptotagmin- GFP. Genesis 34, 142–145, doi:10.1002/gene.10144 (2002).

39. Talay, M. et al. Transsynaptic Mapping of Second-Order Taste Neurons in Flies by trans- Tango. Neuron 96, 783–795 e784, doi:10.1016/j.neuron.2017.10.011 (2017).

40. Lima, S. Q. & Miesenbock, G. Remote control of behavior through genetically targeted photostimulation of neurons. Cell 121, 141–152, doi:10.1016/j.cell.2005.02.004 (2005).

41. Yao, Z., Macara, A. M., Lelito, K. R., Minosyan, T. Y. & Shafer, O. T. Analysis of functional neuronal connectivity in the Drosophila brain. J Neurophysiol 108, 684–696, doi:10.1152/jn.00110.2012 (2012).

42. Chen, T. W. et al. Ultrasensitive fluorescent proteins for imaging neuronal activity. Nature 499, 295–300, doi:10.1038/nature12354 (2013).

43. Fernandez, M. P., Berni, J. & Ceriani, M. F. Circadian remodeling of neuronal circuits involved in rhythmic behavior. PLoS Biol 6, e69, doi:10.1371/journal.pbio.0060069 (2008).

44. Sivachenko, A., Li, Y., Abruzzi, K. C. & Rosbash, M. The transcription factor Mef2 links the Drosophila core clock to Fas2, neuronal morphology, and circadian behavior. Neuron 79, 281–292, doi:10.1016/j.neuron.2013.05.015 (2013).

45. Gorostiza, E. A., Depetris-Chauvin, A., Frenkel, L., Pirez, N. & Ceriani, M. F. Circadian pacemaker neurons change synaptic contacts across the day. Curr Biol 24, 2161–2167, doi:10.1016/j.cub.2014.07.063 (2014).

46. Petsakou, A., Sapsis, T. P. & Blau, J. Circadian Rhythms in Rho1 Activity Regulate Neuronal Plasticity and Network Hierarchy. Cell 162, 823–835, doi:10.1016/j.cell.2015.07.010 (2015).

47. Agrawal, P. & Hardin, P. E. The Drosophila Receptor Protein Tyrosine Phosphatase LAR Is Required for Development of Circadian Pacemaker Neuron Processes That Support Rhythmic Activity in Constant Darkness But Not during Light/Dark Cycles. J Neurosci 36, 3860–3870, doi:10.1523/JNEUROSCI.4523-15.2016 (2016).

48. Prakash, P., Nambiar, A. & Sheeba, V. Oscillating PDF in termini of circadian pacemaker neurons and synchronous molecular clocks in downstream neurons are not sufficient for sustenance of activity rhythms in constant darkness. PLoS One 12, e0175073, doi:10.1371/journal.pone.0175073 (2017).

49. Kula, E., Levitan, E. S., Pyza, E. & Rosbash, M. PDF cycling in the dorsal protocerebrum of the Drosophila brain is not necessary for circadian clock function. J Biol Rhythms 21, 104–117, doi:10.1177/0748730405285715 (2006).

50. Muraro, N. I., Pirez, N. & Ceriani, M. F. The circadian system: plasticity at many levels. Neuroscience 247, 280–293, doi:10.1016/j.neuroscience.2013.05.036 (2013).

51. Fernandez, M. P. et al. Sites of Circadian Clock Neuron Plasticity Mediate Sensory Integration and Entrainment. Curr Biol, doi:10.1016/j.cub.2020.04.025 (2020).

52. Wu, Y., Cao, G. & Nitabach, M. N. Electrical silencing of PDF neurons advances the phase of non-PDF clock neurons in Drosophila. J Biol Rhythms 23, 117–128, doi:10.1177/0748730407312984 (2008).

53. Dajani, D. R. & Uddin, L. Q. Demystifying cognitive flexibility: Implications for clinical and developmental neuroscience. Trends Neurosci 38, 571–578, doi:10.1016/j.tins.2015.07.003 (2015).

54. Liu, Q. et al. Branch-specific plasticity of a bifunctional dopamine circuit encodes protein hunger. Science 356, 534–539, doi:10.1126/science.aal3245 (2017).

55. Hart, M. P. & Hobert, O. Neurexin controls plasticity of a mature, sexually dimorphic neuron. Nature 553, 165–170, doi:10.1038/nature25192 (2018).

56. Farris, S. M., Robinson, G. E. & Fahrbach, S. E. Experience- and age-related outgrowth of intrinsic neurons in the mushroom bodies of the adult worker honeybee. J Neurosci 21, 6395–6404 (2001).

57. Kim, S. S., Rouault, H., Druckmann, S. & Jayaraman, V. Ring attractor dynamics in the Drosophila central brain. Science 356, 849–853, doi:10.1126/science.aal4835 (2017).

58. Liang, X. et al. Morning and Evening Circadian Pacemakers Independently Drive Premotor Centers via a Specific Dopamine Relay. Neuron 102, 843–857 e844, doi:10.1016/j.neuron.2019.03.028 (2019).

59. Potdar, S. & Sheeba, V. Wakefulness Is Promoted during Day Time by PDFR Signalling to Dopaminergic Neurons in Drosophila melanogaster. eNeuro 5, doi:10.1523/ENEURO.0129-18.2018 (2018).

60. Gorska-Andrzejak, J., Chwastek, E. M., Walkowicz, L. & Witek, K. On Variations in the Level of PER in Glial Clocks of Drosophila Optic Lobe and Its Negative Regulation by PDF Signaling. Front Physiol 9, 230, doi:10.3389/fphys.2018.00230 (2018).

61. Park, J. H. et al. Differential regulation of circadian pacemaker output by separate clock genes in Drosophila. Proc Natl Acad Sci U S A 97, 3608–3613, doi:10.1073/pnas.070036197 (2000).

62. Ni, J. Q. et al. Vector and parameters for targeted transgenic RNA interference in Drosophila melanogaster. Nat Methods 5, 49–51, doi:10.1038/nmeth1146 (2008).

63. Pfeiffer, B. D. et al. Refinement of tools for targeted gene expression in Drosophila. Genetics 186, 735–755, doi:10.1534/genetics.110.119917 (2010).

64. Mauss, A. S., Busch, C. & Borst, A. Optogenetic Neuronal Silencing in Drosophila during Visual Processing. Sci Rep 7, 13823, doi:10.1038/s41598-017-14076-7 (2017).

65. Zhang, S. X., Rogulja, D. & Crickmore, M. A. Dopaminergic Circuitry Underlying Mating Drive. Neuron 91, 168–181, doi:10.1016/j.neuron.2016.05.020 (2016).

66. Boutros, C. L., Miner, L. E., Mazor, O. & Zhang, S. X. Measuring and Altering Mating Drive in Male Drosophila melanogaster. J Vis Exp, doi:10.3791/55291 (2017).

67. Tayler, T. D., Pacheco, D. A., Hergarden, A. C., Murthy, M. & Anderson, D. J. A neuropeptide circuit that coordinates sperm transfer and copulation duration in Drosophila. Proc Natl Acad Sci U S A 109, 20697–20702, doi:10.1073/pnas.1218246109 (2012).

68. Veenstra, J. A. & Ida, T. More Drosophila enteroendocrine peptides: Orcokinin B and the CCHamides 1 and 2. Cell Tissue Res 357, 607–621, doi:10.1007/s00441-014-1880-2 (2014).

69. Busza, A., Emery-Le, M., Rosbash, M. & Emery, P. Roles of the two Drosophila CRYPTOCHROME structural domains in circadian photoreception. Science 304, 1503–1506, doi:10.1126/science.1096973 (2004).

70. Berndt, A. et al. A novel photoreaction mechanism for the circadian blue light photoreceptor Drosophila cryptochrome. J Biol Chem 282, 13011–13021, doi:10.1074/jbc.M608872200 (2007).

71. VanVickle-Chavez, S. J. & Van Gelder, R. N. Action spectrum of Drosophila cryptochrome. J Biol Chem 282, 10561–10566, doi:10.1074/jbc.M609314200 (2007).

72. Hanai, S., Hamasaka, Y. & Ishida, N. Circadian entrainment to red light in Drosophila: requirement of Rhodopsin 1 and Rhodopsin 6. Neuroreport 19, 1441–1444, doi:10.1097/WNR.0b013e32830e4961 (2008).

73. Fogle, K. J., Parson, K. G., Dahm, N. A. & Holmes, T. C. CRYPTOCHROME is a blue-light sensor that regulates neuronal firing rate. Science 331, 1409–1413, doi:10.1126/science.1199702 (2011).

74. Fogle, K. J. et al. CRYPTOCHROME-mediated phototransduction by modulation of the potassium ion channel beta-subunit redox sensor. Proc Natl Acad Sci U S A 112, 2245–2250, doi:10.1073/pnas.1416586112 (2015).

75. Hendricks, J. C. et al. Rest in Drosophila is a sleep-like state. Neuron 25, 129–138 (2000).

76. Shaw, P. J., Cirelli, C., Greenspan, R. J. & Tononi, G. Correlates of sleep and waking in Drosophila melanogaster. Science 287, 1834–1837 (2000).

77. Klarsfeld, A., Leloup, J.-C. & Rouyer, F. Circadian rhythms of locomotor activity in Drosophila. Behavioural Processes 64, 161–175, doi:10.1016/s0376-6357(03)00133-5 (2003).

78. Zielinski, T., Moore, A. M., Troup, E., Halliday, K. J. & Millar, A. J. Strengths and limitations of period estimation methods for circadian data. PLoS One 9, e96462, doi:10.1371/journal.pone.0096462 (2014).

79. Houl, J. H., Ng, F., Taylor, P. & Hardin, P. E. CLOCK expression identifies developing circadian oscillator neurons in the brains of Drosophila embryos. BMC Neurosci 9, 119, doi:10.1186/1471-2202-9-119 (2008).

80. Shafer, O. T., Rosbash, M. & Truman, J. W. Sequential nuclear accumulation of the clock proteins period and timeless in the pacemaker neurons of Drosophila melanogaster. J Neurosci 22, 5946–5954, doi:20026628 (2002).

81. Siwicki, K. K., Eastman, C., Petersen, G., Rosbash, M. & Hall, J. C. Antibodies to the period gene product of Drosophila reveal diverse tissue distribution and rhythmic changes in the visual system. Neuron 1, 141–150, doi:10.1016/0896-6273(88)90198-5 (1988).

82. Zerr, D. M., Hall, J. C., Rosbash, M. & Siwicki, K. K. Circadian fluctuations of period protein immunoreactivity in the CNS and the visual system of Drosophila. J Neurosci 10, 2749–2762 (1990).

83. Shafer, O. T. & Taghert, P. H. RNA-interference knockdown of Drosophila pigment dispersing factor in neuronal subsets: the anatomical basis of a neuropeptide’s circadian functions. PLoS One 4, e8298, doi:10.1371/journal.pone.0008298 (2009).

84. Baines, R. A., Uhler, J. P., Thompson, A., Sweeney, S. T. & Bate, M. Altered electrical properties in Drosophila neurons developing without synaptic transmission. J Neurosci 21, 1523–1531 (2001).

